# Discovery of Potent Pyrazoline-Based Covalent SARS-CoV-2 Main Protease Inhibitors

**DOI:** 10.1101/2022.03.05.483025

**Authors:** Patrick Moon, Lydia Boike, Dustin Dovala, Nathaniel J. Henning, Mark Knapp, Jessica N. Spradlin, Carl C. Ward, Helene Wolleb, Charlotte M. Zammit, Daniel Fuller, Gabrielle Blake, Jason P. Murphy, Feng Wang, Yipin Lu, Stephanie A. Moquin, Laura Tandeske, Matthew J. Hesse, Jeffrey M. McKenna, John A. Tallarico, Markus Schirle, F. Dean Toste, Daniel K. Nomura

## Abstract

While vaccines and antivirals are now being deployed for the current SARS-CoV-2 pandemic, we require additional antiviral therapeutics to not only effectively combat SARS-CoV-2 and its variants, but also future coronaviruses. All coronaviruses have relatively similar genomes that provide a potential exploitable opening to develop antiviral therapies that will be effective against all coronaviruses. Among the various genes and proteins encoded by all coronaviruses, one particularly “druggable” or relatively easy-to-drug target is the coronavirus Main Protease (3CL^pro^ or Mpro), an enzyme that is involved in cleaving a long peptide translated by the viral genome into its individual protein components that are then assembled into the virus to enable viral replication in the cell. Inhibiting Mpro with a small-molecule antiviral would effectively stop the ability of the virus to replicate, providing therapeutic benefit. In this study, we have utilized activity-based protein profiling (ABPP)-based chemoproteomic approaches to discover and further optimize cysteine-reactive pyrazoline-based covalent inhibitors for the SARS-CoV-2 Mpro. Structure-guided medicinal chemistry and modular synthesis of di- and tri-substituted pyrazolines bearing either chloroacetamide or vinyl sulfonamide cysteine-reactive warheads enabled the expedient exploration of structure-activity relationships (SAR), yielding nanomolar potency inhibitors against Mpro from not only SARS-CoV-2, but across many other coronaviruses. Our studies highlight promising chemical scaffolds that may contribute to future pan-coronavirus inhibitors.

## Introduction

While vaccines are now being deployed for the current SARS-CoV-2 pandemic, we need more antiviral therapeutics that can effectively combat this COVID-19 pandemic and future coronaviruses that will inevitably arise. Furthermore, with the appearance of an increasing number of dangerous variants of SARS-CoV-2 and with all vaccines and antibody therapies targeting the viral Spike protein, which is prone to mutations, there is a need for a pan-coronavirus antiviral drug that will be effective against any variant of the SARS-CoV-2 virus as well as future coronaviruses.

All coronaviruses have several highly conserved genes that provide a potential exploitable opening to develop an antiviral therapy that could be effective against not only SARS-CoV-2, but also all past and future coronaviruses. Among the various genes and proteins encoded by coronaviruses, one particularly “druggable” or relatively easy-to-drug target is the coronavirus Main Protease (Mpro) also known as 3C-like protease, an enzyme that is involved in cleaving a long peptide translated by the viral genome into its individual protein components that are then assembled into the virus to enable viral replication in the cell. The replicase gene of SARS-CoV-2 encodes two overlapping polyproteins—pp1a and pp1ab—that are required for viral replication and transcription ^1^. Mpro is responsible for the majority of proteolytic processing events to release the functional polypeptides with at least 11 proteolytic cleavage events starting with the autolytic cleavage of Mpro itself from pp1a and pp1ab ^1^. Inhibiting Mpro with a small-molecule antiviral could effectively stop the ability of the virus to replicate, providing immediate and complete therapeutic benefit. Unlike the viral Spike protein that currently all vaccines and antibody therapeutics target, which is highly prone to mutations that give rise to resistant variants, Mpro protein sequence is highly conserved across all former coronaviruses indicating that making a potent and effective Mpro inhibitor for one coronavirus would result in an efficacious drug for all future coronaviruses. Because Mpro is a critical protein necessary for coronavirus replication, mutations in the core active site domain of this protein would likely impair its catalytic activity and the ability of the virus to replicate, and thus Mpro inhibitor antivirals would be unlikely to run into resistance through viral mutations. There has been considerable effort over the past two years by many academic and industrial labs to discover Mpro inhibitors **(Fig. 1A)** ^2,3^. For example, Pfizer has now developed an effective orally bioavailable covalently-acting peptide-based Mpro inhibitor Paxlovid that has been given emergency use authorization by the United States Food and Drug Administration **(Fig. 1A)** ^4^. However, there is still significant room for discovering additional Mpro inhibitor scaffolds that may be tractable for drug discovery and development efforts, leading to even more efficacious treatments. Additionally, there may be the emergence of resistance against Paxlovid once widely deployed in the clinic, necessitating additional inhibitors ^5,6^.

**Figure 1.**
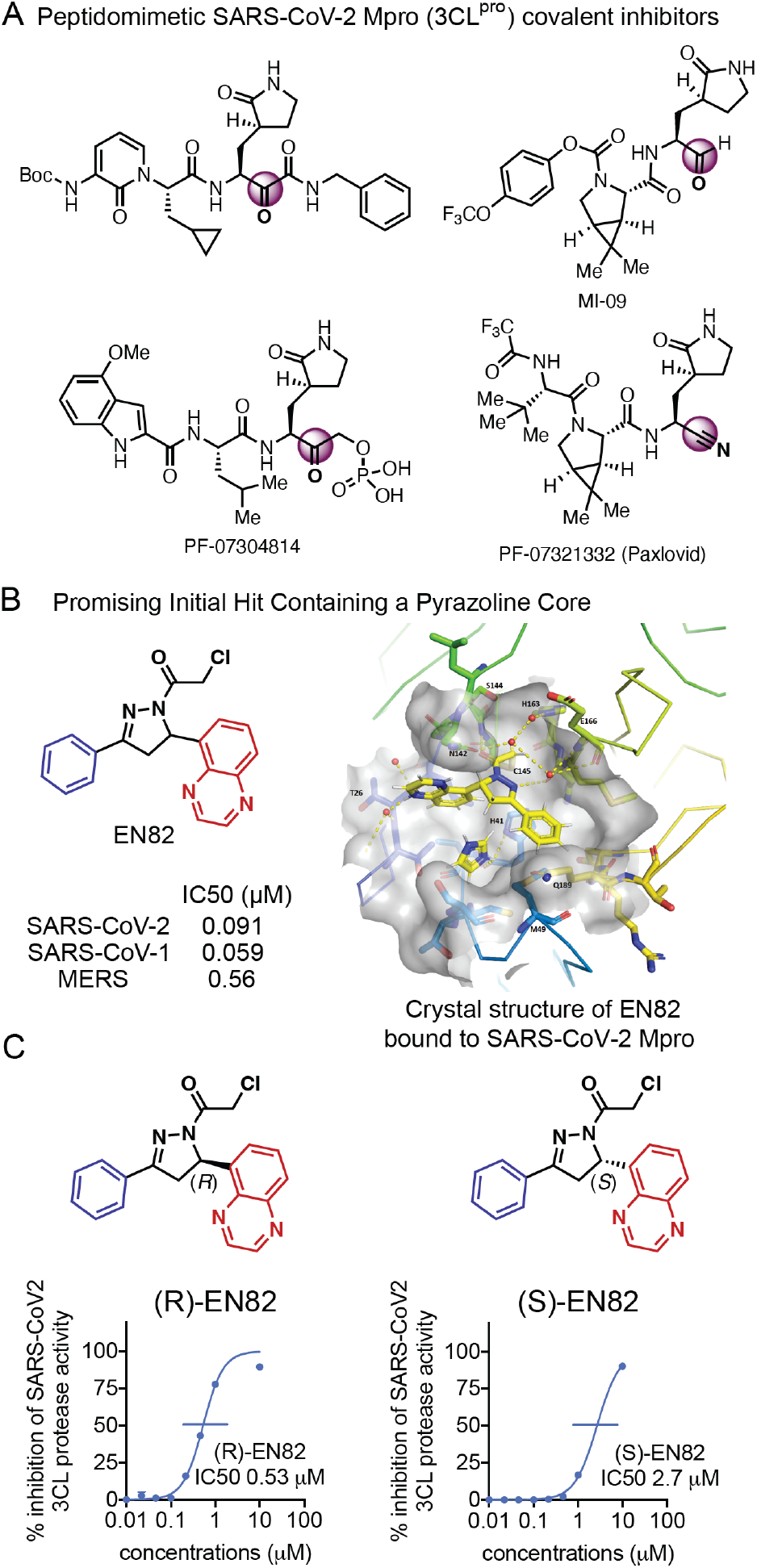
Discovery of pyrazoline-based SARS-CoV-2 main protease inhibitors. **(A)** Examples of peptidic SARS-CoV-2 Mpro inhibitors. **(B)** Structure of our top hit EN82 and potency of EN82 against SARS-CoV-2, SARS-CoV-1, and MERS-CoV Mpro. Apparent IC50s were derived from a rhodamine-based Mpro substrate peptide activity assay. On the right is a crystal structure of EN82 covalently bound to the catalytic C145 in the SARS-CoV-2 Mpro active site. **(C)** Potency of (*R*)- and (*S*)-EN82 against SARS-CoV-2 Mpro. Apparent IC50s were derived from averages of 3 biological replicates/group.

As a cysteine protease, the catalytic activity of Mpro is driven by its catalytic cysteine C145 ^1^. Therefore, chemoproteomic platforms and cysteine-reactive covalent ligand discovery approaches are particularly attractive for developing potent, selective, non-peptidic, more drug-like, and covalently-acting Mpro inhibitors that can irreversibly interact with the Mpro catalytic cysteine.

To rapidly identify cysteine-reactive inhibitors against SARS-CoV-2 Mpro, we screened a library of 582 acrylamides and chloroacetamides in a gel-based activity-based protein profiling (ABPP) screen, in which we competed these cysteine-reactive covalent ligands against the binding of a cysteine-reactive rhodamine-functionalized iodoacetamide probe (IA-rhodamine) using previously described gel-based ABPP approaches ^7,8^ **(Fig. S1-S2)**. The most promising hits clustered together on pyrazoline-based chloroacetamide ligands EN71, EN82, EN216, and EN223 that demonstrated dose-responsive inhibition of Mpro IA-rhodamine labeling **(Fig. S3A).** We subsequently tested these four compounds in a FRET-based activity assay employing an Mpro peptide substrate to identify compounds that inhibited Mpro activity (**Fig. S3B**). The most promising hit to arise from this screen was the pyrazoline EN82 which displayed encouraging potency against the SARS-CoV-2 Mpro with an apparent 50% inhibitory concentration (IC50) value for an incubation time of 30 min of 0.16 μM compared to 0.52-4.8 μM for EN216, EN71, and EN223 **(Fig. S3B).** The FRET-based peptide probe required significant concentrations of Mpro protein (115 nM) to obtain reliable Mpro activity readouts, hindering our ability to detect more potent inhibitors with IC50 values <100 nM. For subsequent Mpro activity assays, we switched over to a rhodamine-based Mpro substrate activity assay which required significantly less protein. Using this rhodamine-based assay, EN82 showed potent apparent IC50 values against Mpro from SARS-CoV-2 (0.091 μM), SARS-CoV-1 (0.059 μM) and MERS-CoV (0.56 μM) (**Fig. 1B**). We next performed chemoproteomic profiling of EN82 cysteine-reactivity in HEK293T cell lysate to assess whether this compound demonstrated some degree of selectivity or whether it was non-specific **(Fig. 2)**. Using a rapid mass spectrometry-based covalent chemoproteomics workflow, in which HEK293T cell lysates spiked with pure SARS-CoV-2 Mpro protein were pre-treated with vehicle or EN82 (10 μM), and subsequently labeled with acid-cleavable cysteine-reactive iodoacetamide-based enrichment probe for quantitative analysis of EN82-competed cysteine sites, we observed Mpro C145 as the primary target of EN82 with one additional off-target HMOX2 C282 out of >1000 distinct quantified probe-modified cysteines. Thus, as an initial hit compound, EN82 showed a high degree of proteome-wide selectivity with only one potential off-target **(Fig. 2)**. EN82 was a racemic mixture of two compounds. Upon enantioselective synthesis of each isomer, (*R*)-EN82 (0.53 μM) proved more active than (*S*)-EN82 (2.7 μM), consistent with a crystal structure obtained of (*R*)-EN82 bound to the catalytic cysteine C145 of SARS-CoV-2 Mpro **(Fig. 1B)**. With EN82 as an encouraging starting point, structure-activity relationship (SAR) studies around the central pyrazoline core were conducted to optimize for potency.

**Figure 2.**
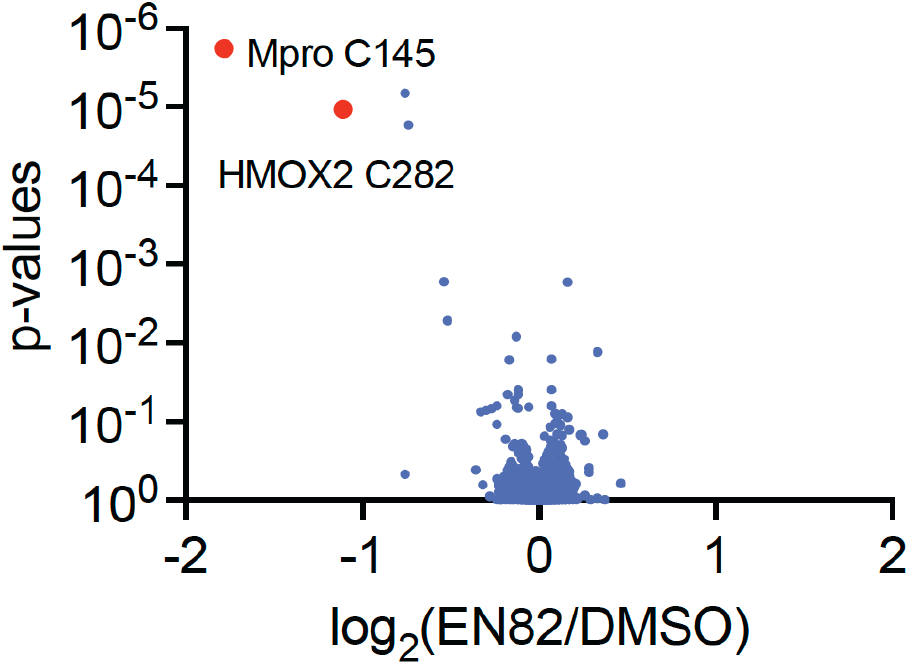
Proteome-wide cysteine-reactivity of EN82 in HEK293T cell lysate with spiked-in SARS-CoV-2 Mpro. SARS-CoV-2 Mpro was spiked into HEK293T cell lysate which was then treated with DMSO or EN82 (10 μM) in triplicate, followed by labeling of proteomes with an iodoacetamide-based enrichment probe followed by TMT-based quantitative mass spectrometry. Shown in red are Mpro C145 and HMOX C282 as the only two sites showing >2-fold competition and p<0.05 out of 1078 distinct quantified cysteine sites.

We next devised an expedient synthesis of 3,5-disubstituted pyrazolines starting with an aldol condensation between commercially available aldehyde and acetophenone precursors to form the corresponding chalcone, followed by hydrazine condensation to access the unprotected pyrazoline core, and subsequent N-acylation to yield the final pyrazoline chloroacetamide in three steps (**Fig. 3A**). A number of nitrogen-containing heterocycles were introduced at the pyrazoline C5-carbon (Ar^1^) in an attempt to improve potency. Differentially substituted quinoxalines, quinolines, indoles, azaindoles, benzotriazoles and pyridines were tested at this position, but no significant breakthrough was achieved (**Fig. 3B**). Several of these derivatives showed comparable potency to EN82, suggesting there is some flexibility for the C5-pyrazoline substituent. SAR at the pyrazoline C3-carbon (Ar^2^) was explored next (**Fig. 3C**). Inhibitors with larger 4-substituted arenes (LEB-2-187, LEB-2-175, LEB-2-174, LEB-2-173) showed decreased potency compared to smaller 3-substituted arenes (LEB-2-172, LEB-3-001, LEB-2-182, LEB-2-176, and LEB-3-004). This was consistent with the relatively shallow pocket in which the C3-arene is located in the EN82-bound Mpro crystal structure (**Fig. 1B**). Simple aliphatic tert-butyl and cyclopropyl substituents (HW-1-174, HW-1-185) at this position exhibited a significant loss of activity. Exploration of SAR was also examined using a 5-quinolinyl bearing pyrazoline scaffold (**Fig. 3C, bottom**), given that this scaffold displayed better overall solubility relative to the 5-quinoxalinyl derivatives. A consistent trend was observed with smaller 3- and 2-substituted arenes at the C3 position showing better activity. Ultimately, a 5-chloro-2-fluorophenyl C3-substituent (CMZ-53) proved optimal and provided a nearly three-fold improvement in potency compared to EN82.

**Figure 3.**
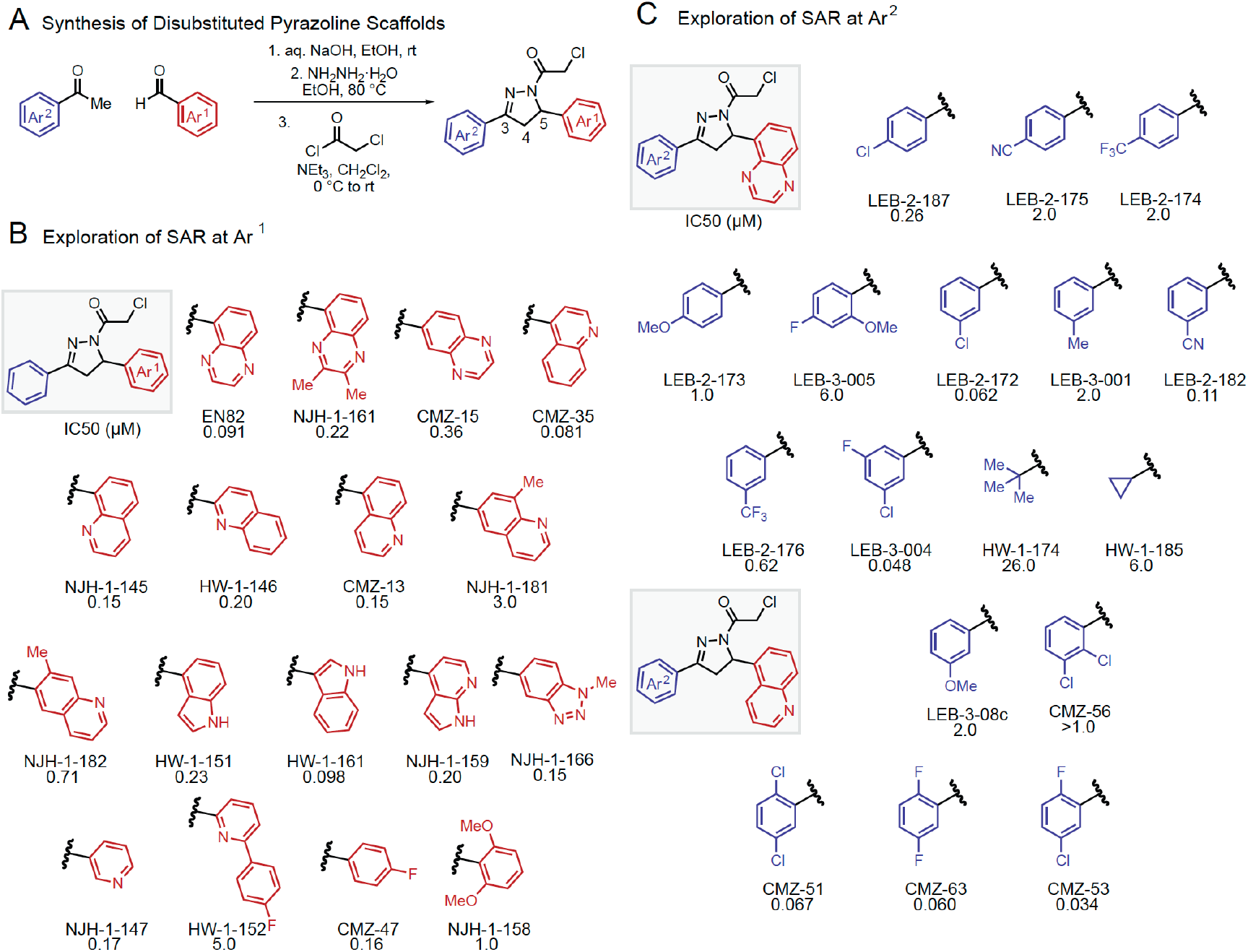
Exploration of 3,5-disubstituted pyrazoline SAR. (A) Synthesis of disubstituted pyrazoline scaffolds. **(B)** Exploration of SAR at Ar^1^. **(C)** Exploration of SAR at Ar^2^. Apparent IC50 values determined by rhodamine-based substrate activity assay for SARS-CoV-2 Mpro from n=3 biological replicates.

We next explored the possibility of adding a substituent at the C4 position of the pyrazoline core. Examination of the crystal structure of EN82 bound to the SARS-CoV-2 Mpro revealed a small pocket occupied by an imidazole that should be accessible with cis-trisubstituted pyrazolines (**Fig. 1B**). Consistent with this premise, the trisubstituted cis-pyrazoline PM-2-020B showed improved potency compared to EN82 and its disubstituted analog EN23 (**Fig. 4B**). In contrast, trans-pyrazoline PM-2-020A showed an approximate 30-fold lower potency compared to cis-PM-02-020B, highlighting the profound impact of relative stereochemistry on the activity of these trisubstituted pyrazolines. A crystal structure of PM-2-020B bound to the SARS-CoV-2 Mpro was obtained which corroborated the initial hypothesis, with the C4-phenyl group nestled in the previously empty subpocket. Trisubstituted analogs with C5-heterocycles were next tested while holding the C4-phenyl group constant. A smaller 3-pyridyl C5-substituent (cis-PM-2-043B) was slightly less potent, while a 5-quinoxalinyl substituent (cis-PM-2-027B) proved even less potent, perhaps due to unfavorable steric clash between the C4/C5 aryl substituents. Consistent with this hypothesis, a smaller isomeric 6-quinoxalinyl derivative (cis-HW-1-187B) proved to be our most potent inhibitor yet (Apparent IC50 = 0.022 μM). Other heteroaryl and alkyl C4-substituents were also tested but showed no significant improvement (HW-1-198B, HW-1-176B, HW-2-01B).

**Figure 4.**
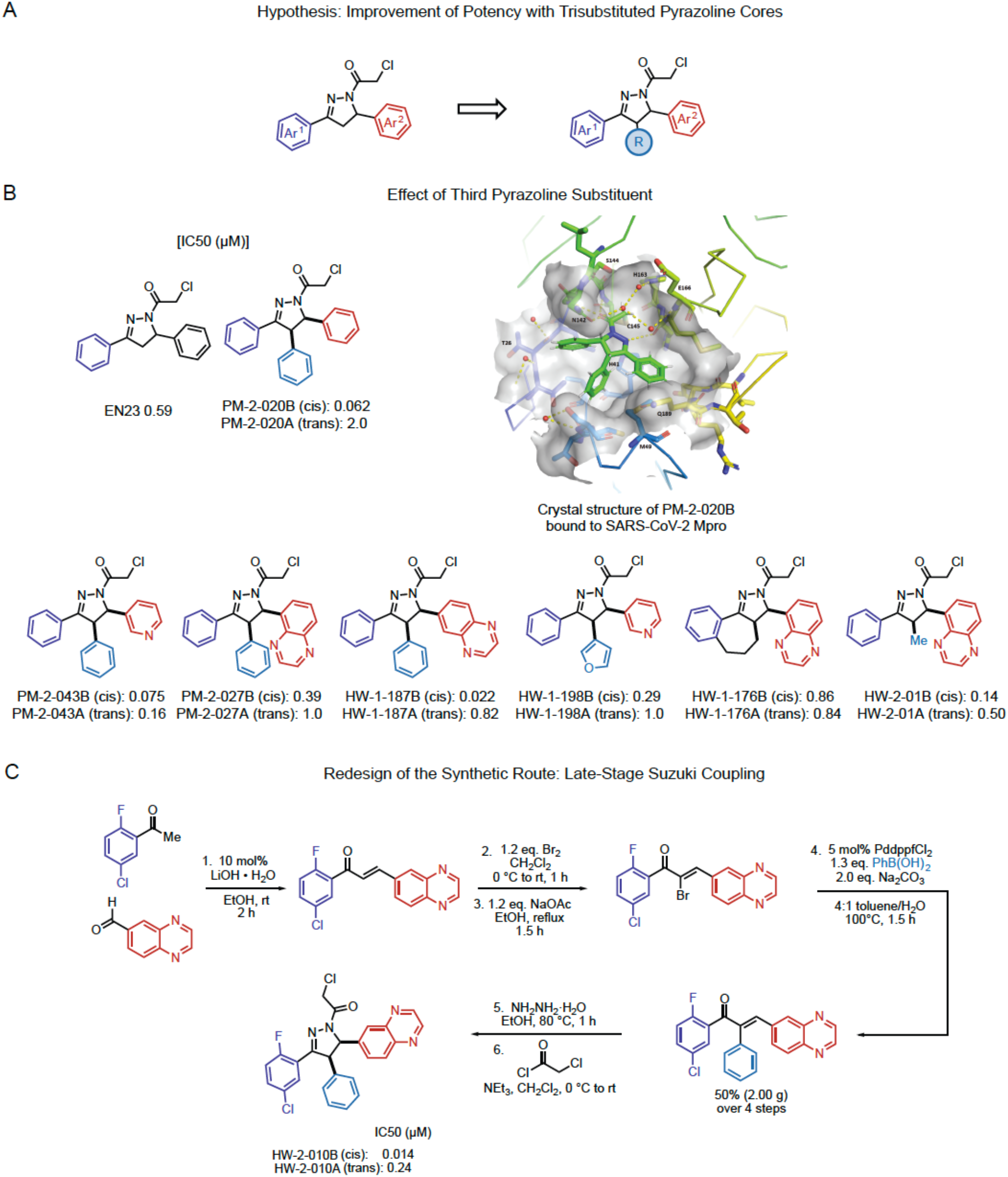
Exploration of trisubstituted pyrazoline inhibitors. **(A)** We hypothesized that potency would be improved through trisubstitution of the pyrazoline core. **(B)** Effect of a third pyrazoline substituent on inhibitory potency of compounds against SARS-CoV-2 Mpro. Shown is the crystal structure of PM-2-20B covalently bound to the catalytic C145 of the SARS-CoV-2 Mpro active site. Apparent IC50 values determined by rhodamine-based substrate activity assay for SARS-CoV-2 Mpro from n=3 biological replicates. **(C)** Redesign of the synthetic route involving a late-stage Suzuki coupling to yield HW-2-010B and HW-2-010A. Potencies against SARS-CoV-2 Mpro are shown from n=3 biological replicates.

A revised modular synthetic route was next designed such that each pyrazoline substituent originates from a different building block. Ideally, the C4-substituent could be installed at a later stage via cross-coupling to avoid having to be introduced in the first aldol condensation step from 2-arylacetophenone derivatives which are not readily available. Initial aldol condensation, followed by mono-bromination and Suzuki coupling led to a tri-arylated chalcone intermediate, which then led to the final pyrazoline core via hydrazine condensation and acylation (**Fig. 4C**). With this route, the optimal 5-chloro-2-fluorophenyl C3-substituent could be installed, leading to our most potent 0.014 μM chloroacetamide inhibitor (HW-2-010B). Along with exploring the SAR around the pyrazoline core, concurrent efforts were aimed at identifying an alternative cysteine reactive warhead given chloroacetamides could potentially present metabolic stability issues (**Fig. 5**). A vinylsulfonamide was ultimately identified as a more promising alternative warhead,^9,10^ culminating in the synthesis of PM-2-071B, a 0.091 μM inhibitor (**Fig. 5, Fig. 6**). Chiral SFC separation of PM-2-071B revealed that the (*S,S*)-enantiomer is most potent (0.035 μM), contrasting what is observed in the chloroacetamide series where the (*R*)-enantiomer of EN82 is most active **(Fig. 6)**. This suggests a different binding mode may be at play with vinylsulfonamides relative to chloroacetamides. Other vinylsulfonamide derivatives were also tested; a modest improvement in potency was obtained with a 3-methoxy C4-substituent (PM-2-167B) while loss in potency was observed with a larger 3-benzyl substituent.

**Figure 5.**
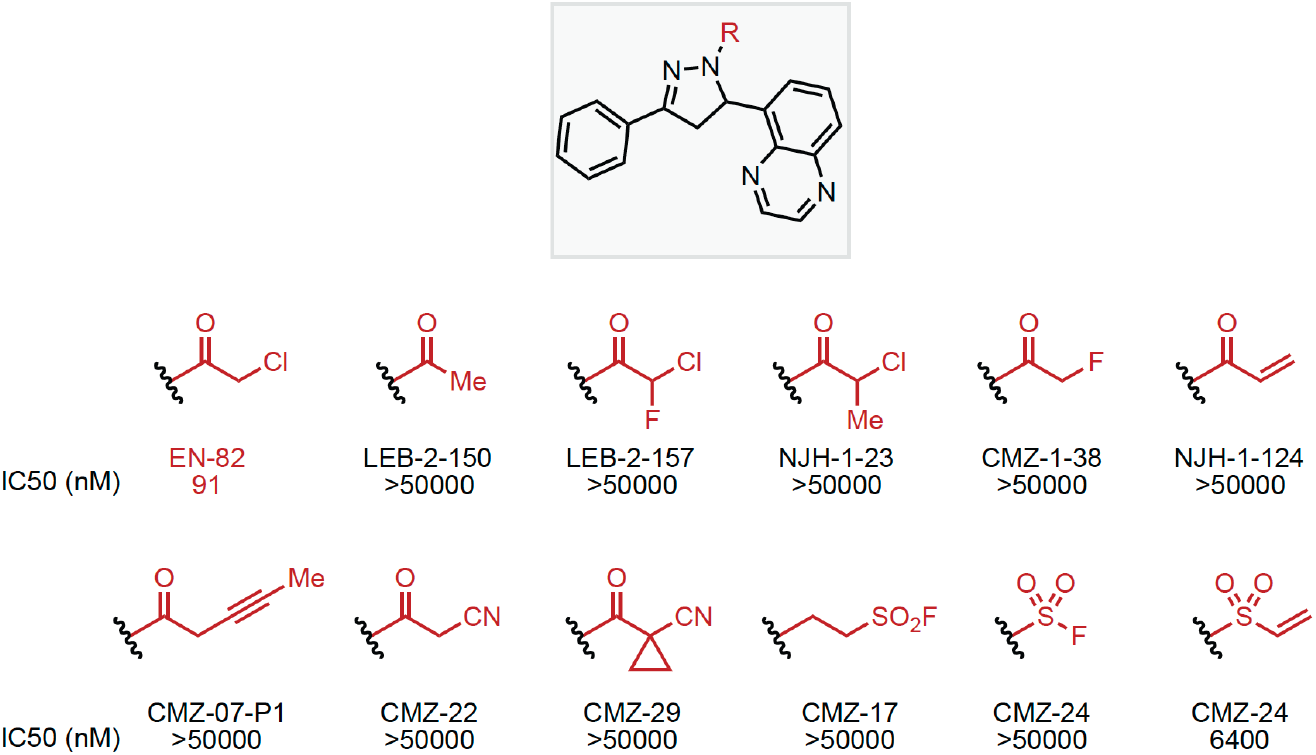
Exploration of Cysteine Reactive Warheads. Apparent IC50 values determined by rhodamine-based substrate activity assay for SARS-CoV-2 Mpro from n=3 biological replicates.

**Figure 6.**
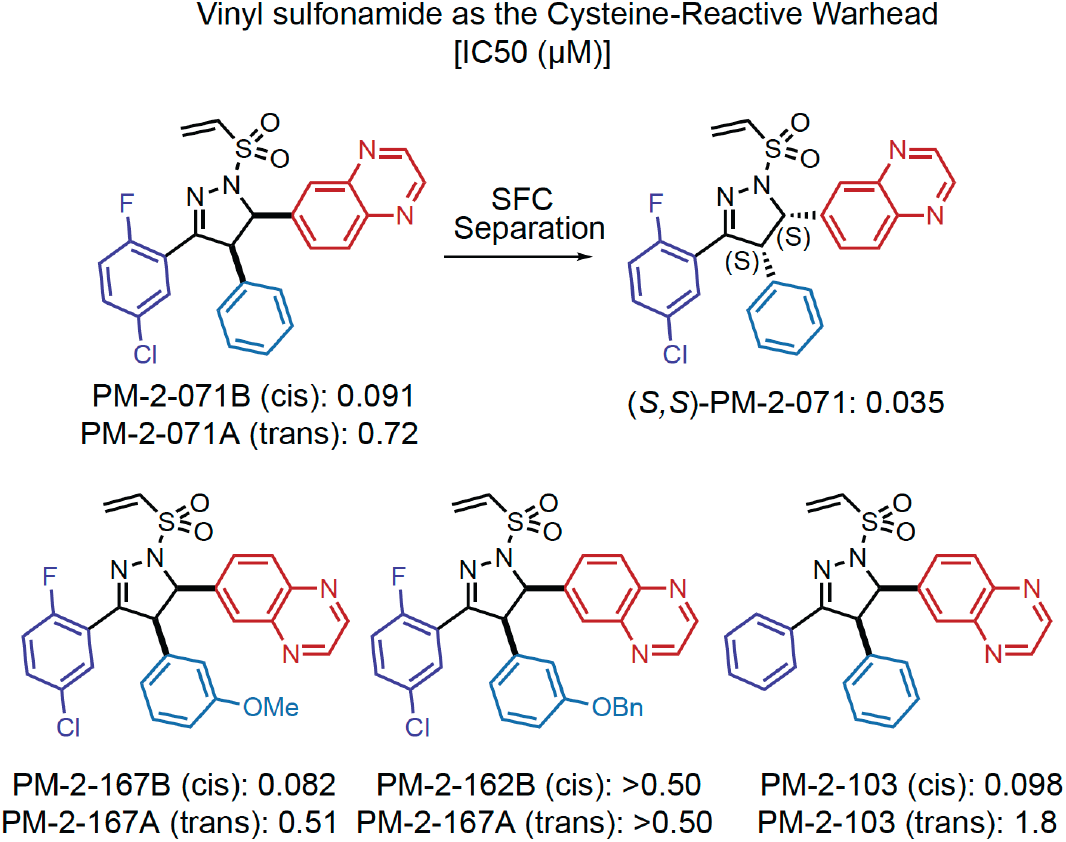
Exploration of trisubstituted pyrazoline inhibitors with a vinyl sulfonamide cysteine-reactive warhead. Apparent IC50 values determined by rhodamine-based substrate activity assay for SARS-CoV-2 Mpro from n=3 biological replicates.

We also tested several of the di- and tri-substituted compounds, including PM-2-071, from our study against a panel of Mpro enzymes from SARS-CoV-2 and other former coronaviruses, including SARS-CoV-1, HCoV-HKU1, HCoV-229E, MERS-CoV, PorCoV-HKU15, HCoV-NL63, avian infectious bronchitis virus (IBV), and HCoV-OC43. For these assays, we used an even more sensitive Mpro activity assay using an Agilent RapidFire-based substrate peptide activity assay that requires even less Mpro protein compared to the rhodamine-based activity assays to comparatively assess potencies of some of our best covalent ligands. In these assays, we show that most of the compounds tested inhibit Mpro from these other coronaviruses with apparent IC50 values in the nanomolar range **(Fig. S4, Fig. 7)**. Most notably, we demonstrate that PM-2-071 shows apparent IC50s <2 nM across SARS-CoV-2, HCoV-HKU1, and HCoV-OC43, and apparent IC50s <1 μM for SARS-CoV-1, HCoV-NL63, IBV, and HCoV-229E Mpro enzymes **(Fig. 7).** Thus, this chemical scaffold may represent a promising starting point for further optimization of pan-coronavirus Mpro inhibitors.

**Figure 7.**
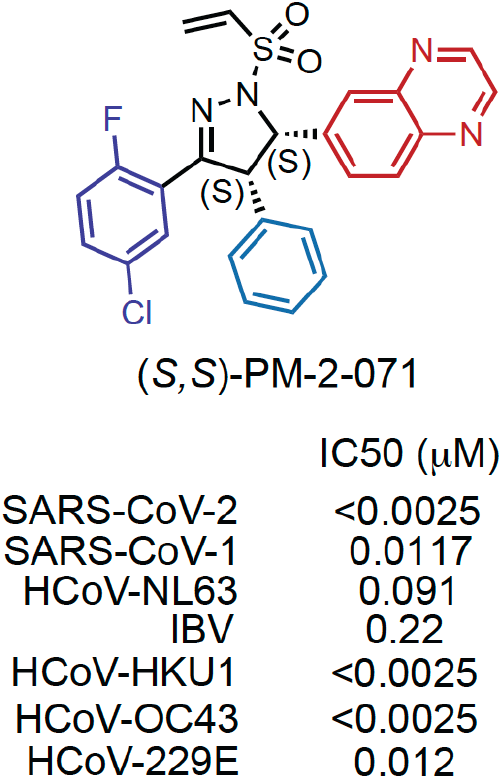
Exploration of trisubstituted pyrazoline inhibitors with a vinyl sulfonamide cysteinereactive warhead. Apparent IC50 values determined by MS-based substrate activity assay for SARS-CoV-2 Mpro.

In conclusion, in this study we have discovered highly potent SARS-CoV-2 Mpro inhibitors based on pyrazoline-based chloroacetamides and vinyl sulfonamides. We further demonstrate that many of the inhibitors developed here also show exceptional potency against Mpro from other coronaviruses, indicating the potential to develop a pan-coronavirus Mpro inhibitor. Exploration of SAR at the pyrazoline core revealed the importance of relative stereochemistry at the C4 and C5 carbons. We note that unfortunately our optimized inhibitors such as *(S,S)-*PM-2-071A appeared to have issues with solubility, metabolic stability, and cell permeability. We did not observe antiviral activity in cells with these series of compounds against SARS-CoV-2. As such, further medicinal chemistry is required to optimize these potent inhibitors for their drug-like properties to promote antiviral efficacy. Nonetheless, we believe that this general scaffold and the exploration of cysteine-reactive warheads here represents a good foundational starting point for the development of more advanced non-peptidic and more drug-like pan-coronavirus Mpro inhibitors in the future.

## Data availability

X-ray structure data have been deposited Structures have been deposited in the RCSB, accession codes: XXXX, YYYY

## Acknowledgements

We thank the members of the Nomura and Toste Research Groups and Novartis Institutes for BioMedical Research for critical reading of the manuscript and Pei-I Ho and Scott Busby for help with assay development. This work was supported by Novartis Institutes for BioMedical Research and the Novartis-Berkeley Center for Proteomics and Chemistry Technologies (NB-CPACT), FAST-GRANTS, and Sergey Brin Family Foundation.

