## Supporting Information for "Discovery of Potent Pyrazoline-Based Covalent SARS-CoV-2 Main Protease Inhibitors"

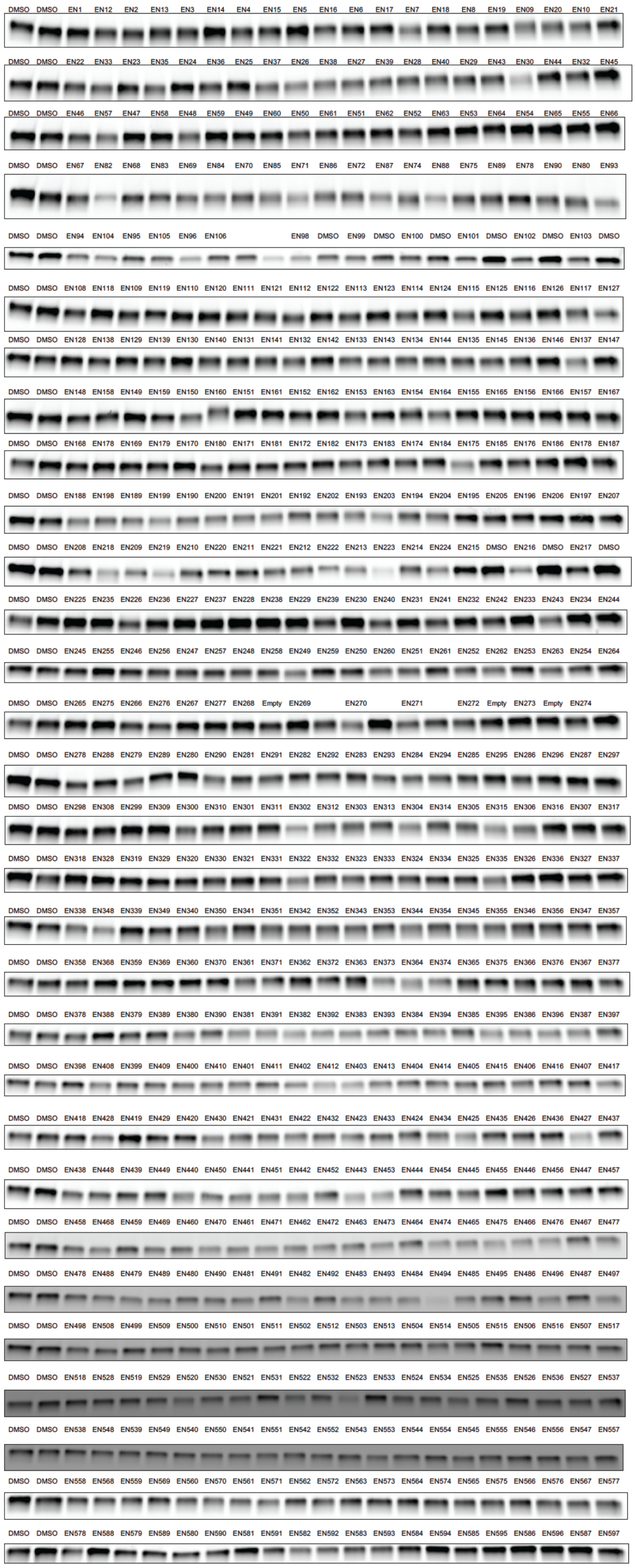


**Figure S1. Gel-based ABPP screen of cysteine-reactive covalent ligands against Mpro enzyme.** Pure SARS-CoV-2 Mpro protein was pre-incubated with DMSO vehicle or cysteine-reactive covalent ligands (50 μM, 30 min) prior to addition of a rhodamine-conjugated iodoacetamide (IA-rhodamine) probe (100 nM) for 1 hr after which reactions were quenched with SDS/PAGE loading buffer, resolved on SDS/PAGE and visualized by in-gel fluorescence.

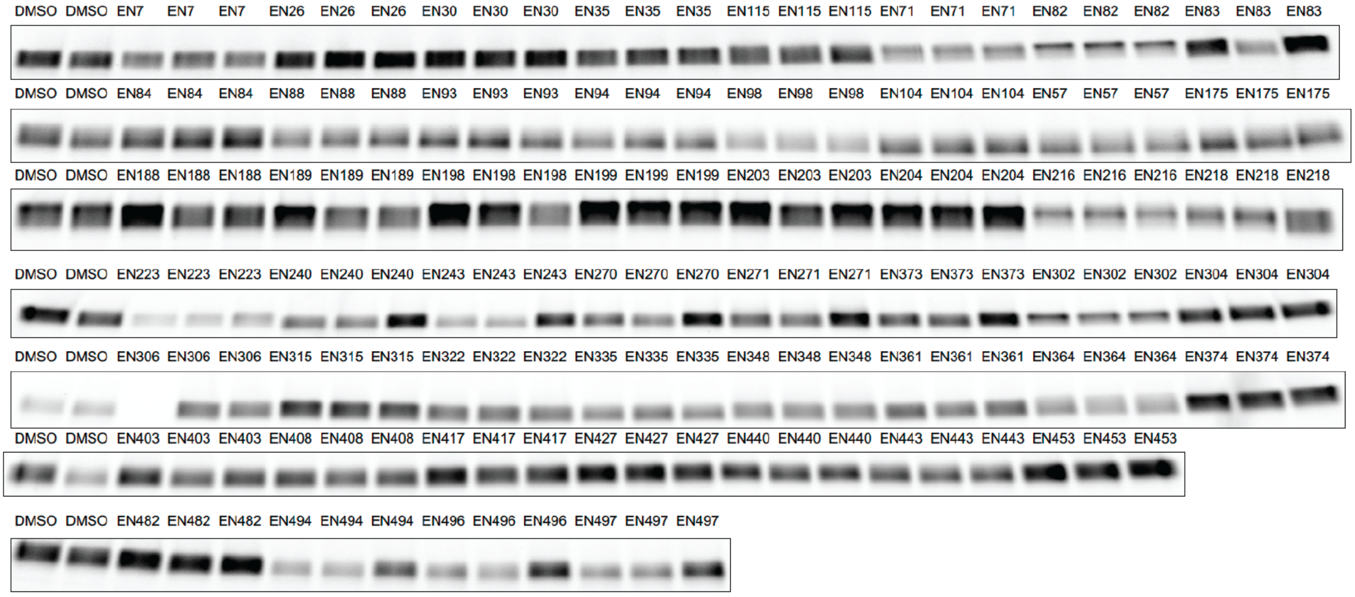


**Figure S2. Hit confirmation of initial gel-based ABPP screen.** Pure SARS-CoV-2 Mpro protein was pre-incubated with DMSO vehicle or cysteine-reactive covalent ligands (50 μM, 30 min) prior to addition of a rhodamine-conjugated iodoacetamide (IA-rhodamine) probe (100 nM) for 1hr min after which reactions were quenched with SDS/PAGE loading buffer, resolved on SDS/PAGE and visualized by in-gel fluorescence.

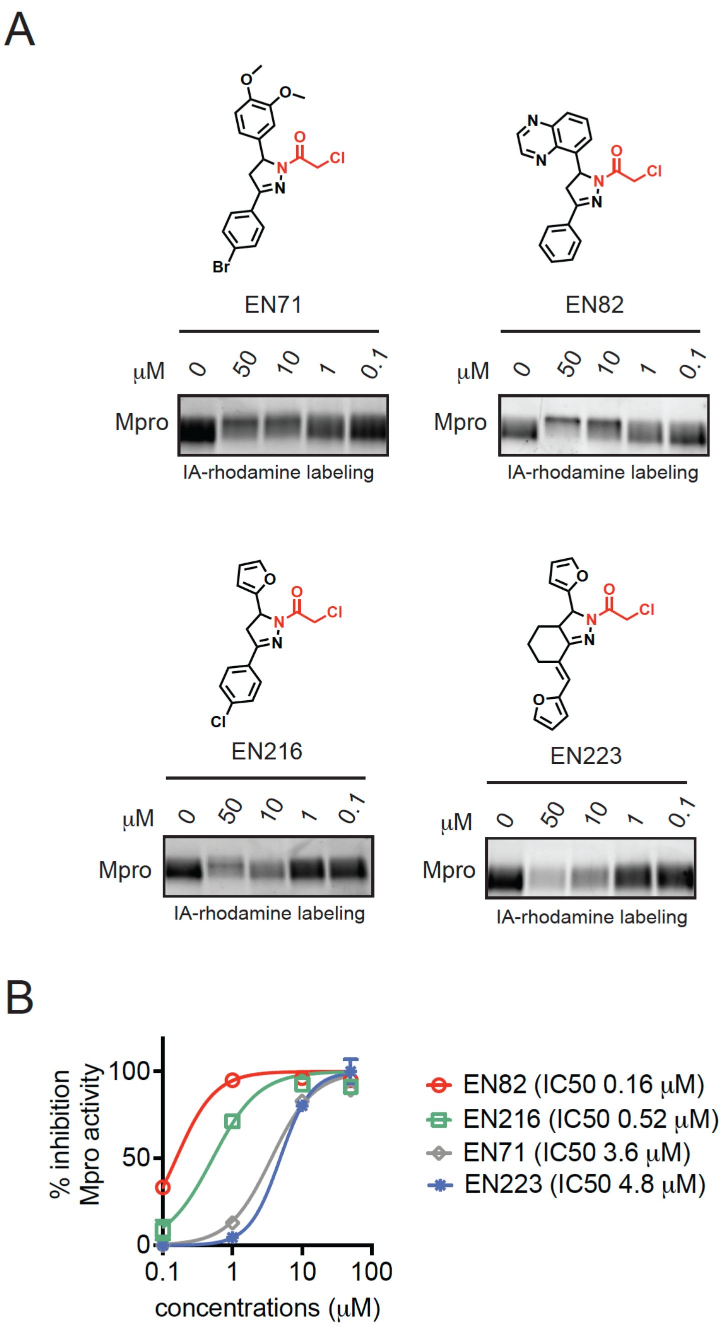


**Figure S3. Testing reproducible hit compounds in dose-response gel-based ABPP and Mpro substrate activity assays. (A)** Gel-based ABPP studies with reproducible hit compounds. Pure SARS-CoV-2 Mpro protein was pre-incubated with DMSO vehicle or cysteine-reactive covalent ligands (30 min) prior to addition of a rhodamine-conjugated iodoacetamide (IA-rhodamine) probe (100 nM) for 1 hr after which reactions were quenched with SDS/PAGE loading buffer, resolved on SDS/PAGE and visualized by in-gel fluorescence. **(B)** Mpro substrate activity assay using an FRET-based peptide probe. Activity assays in **(B)** are from n=3 biological replicates/group and values are expressed as average ± sem.

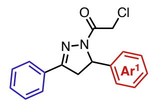


IC50 (μM)

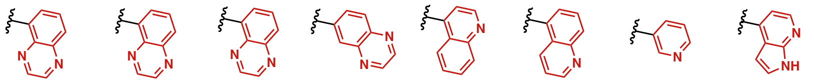

| | EN82 | (R)-EN82 | (S)-EN82 | CMZ-15 | CMZ-35 | CMZ-13 | NJH-1-147 | NJH-1-159 |
|---|---|---|---|---|---|---|---|---|
| SARS-CoV-2 | 0.025 | 0.30 | 0.17 | 0.010 | 0.0046 | 0.0027 | 0.058 | 0.02 |
| HCoV-HKU1 | 0.0066 | 0.0027 | 0.0052 | 0.0031 | 0.0023 | 0.0098 | 0.026 | 0.0089 |
| HCoV-229E | 0.029 | 0.015 | 0.053 | 0.011 | 0.0037 | 0.0045 | 0.054 | 0.019 |
| MERS-CoV | 1.56 | 0.67 | 2.4 | 0.18 | 0.14 | 0.31 | 7.2 | 0.96 |
| PorCoV-HKU15 | 10 | >25 | 7.4 | 8.2 | 1.6 | 5.0 | 11 | 9.3 |
| HCoV-NL63 | 0.24 | 0.13 | 0.55 | 0.11 | 0.016 | 0.0038 | 0.33 | 0.11 |

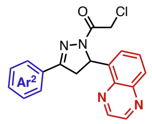


IC50 (μM)

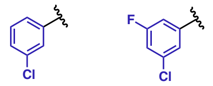


| | LEB-2-172 | LEB-3-004 |
|---|---|---|
| SARS-CoV-2 | 0.00089 | 0.0016 |
| HCoV-HKU1 | 0.00096 | 0.0023 |
| HCoV-229E | 0.0055 | 0.0053 |
| MERS-CoV | 0.038 | 0.10 |
| PorCoV-HKU15 | — | 2.1 |
| HCoV-NL63 | 0.011 | 0.025 |

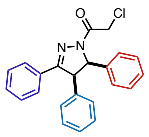

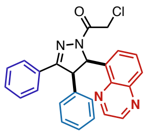

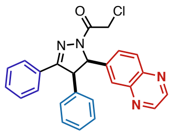

| | PM-2-20A *(trans)* | PM-2-20B *(cis)* | PM-2-27A *(trans)* | PM-2-27B *(cis)* | HW-1-87A *(trans)* | HW-1-187B *(cis)* |
|---|---|---|---|---|---|---|
| SARS-CoV-2 | 0.028 | 0.0053 | 0.11 | 0.071 | 0.085 | 0.0014 |
| HCoV-HKU1 | 0.036 | 0.20 | 0.060 | 0.22 | 0.040 | 0.016 |
| HCoV-229E | 0.0036 | 0.023 | 0.046 | 0.09 | 0.030 | 0.0023 |
| MERS-CoV | 3.7 | 2.8 | 3.3 | 4.3 | 0.34 | 1.3 |
| PorCoV-HKU15 | 4.8 | 1.1 | 5.4 | 7.5 | 4.9 | 6.3 |
| HCoV-NL63 | 0.067 | 0.35 | 0.092 | 0.59 | 0.14 | 0.056 |

**Figure S4. Potency (IC50, μM) of inhibitors against MPro enzymes from other coronaviruses.** The Mpro activity assays were performed with a MS-based substrate activity assay.

### Methods and Materials

#### Production of authentic SARS-CoV-2 Main Proteases

The coding sequence for SARS-CoV-2 main protease was codon-optimized for *E. coli* and synthesized by Integrated DNA Technologies. The sequence was amplified by PCR and cloned into the pGEX6P-1 vector, downstream of GST and an HRV 3C protease cleavage site, using the Gibson Assembly Master Mix kit (New England BioLabs, Inc). To ensure authentic termini, the amino acids AVLQ were added to the N-terminus of the main protease by addition of their coding sequence to the 5′ end of the gene product. This sequence reconstitutes the NSP4/5 cleavage site, resulting in auto-cleavage by the main protease protein product and removal of the GST tag. We also added a GP-6xHis tag (to enable IMAC purification) on the C-terminus (the GP completes a non-consensus 3C cleavage site along with the C-terminus of the main protease which allows for cleavage of the his tag after purification, resulting in an authentic C-terminus).

Hi Control BL21(DE3) cells were transformed with the expression plasmid using standard techniques. We used Hi Control cells as we observed expression of the main protease was toxic in other standard *E. coli* cell lines. A single colony was used to start an overnight culture in LB + carbenicillin media. This culture was used to inoculate 2 x 1 L cultures in Terrific Broth, supplemented with 50 mM sodium phosphate pH 7.0 and 100 µg/mL carbenicillin. These cultures grew in Fernbach flasks at 37 °C while shaking at 225 rpm, until the OD600 reached approximately 2.0, at which point the temperature was reduced to 20 °C and 0.5 mM IPTG (final) was added to each culture. The cells were allowed to grow overnight.

The next day, the cultures were centrifuged at 6,000 x *g* for 20 minutes at 4 °C, and the resulting cell pellets were resuspended in IMAC_A buffer (50 mM Tris pH 8.0, 400 mM NaCl, 1 mM TCEP). Cells were lysed with two passes through a cell homogenizer (Microfluidics model M-110P) at 18,000 psi. The lysate was clarified with centrifugation at 42,000 x *g* for 30 minutes and the cleared lysate was loaded onto 3 x 5 mL HiTrap Ni-NTA columns (GE) pre-equilibrated with IMAC_A buffer, using an AKTA Pure FPLC. After loading, the columns were washed with IMAC_A buffer until the A280 levels reached a sustained baseline. The protein was then eluted with a linear gradient with IMAC_B buffer (50 mM Tris pH 8.0, 400 mM NaCl, 500 mM imidazole, 1 mM TCEP) across 25 column volumes, while 2 mL fractions were collected automatically. Peak fractions were analyzed by SDS-PAGE and those containing SARS-CoV-2 main protease were pooled. Importantly, auto-cleavage of the N-terminal GST tag was observed and the eluted protein had a mass consistent with SARS-CoV-2 main protease along with the C-terminal GP-6xHis tag, as determined by ESI-LC/MS.

Pooled fractions were treated with HRV 3C protease (also known as “PreScission” protease) while dialyzing against IMAC_A buffer at room temperature (2 x 2 L dialyses). Room temperature dialysis was important as we observed a tendency for the main protease protein to precipitate with prolonged exposure to 4 °C. Cleavage of the C-terminal GP-6xHis tag was confirmed after 2 hours by ESI-LC/MS. The dialyzed and cleaved protein was then re-run through a 5 mL HiTrap Ni-NTA column pre-equilibrated with IMAC_A buffer. The main protease eluted in the flow-through as expected.

The protein was then concentrated to approximately 5 mL and loaded onto a Superdex 75 16/60 column pre-equilibrated with SEC Buffer (25 mM HEPES pH 7.5, 150 mM NaCl, 1 mM TCEP). The protein was run through the column at 1 mL/min and eluted as one large peak well in the included volume (at ~75 mL). Fractions from this peak were analyzed by SDS-PAGE and pure fractions were pooled and concentrated to 10 mg/mL, aliquotted, and stored at -80 °C. Final yield was typically in the realm of 60-70 mg/L of culture.

#### Crystallography of Compounds with Mpro

Co-crystallization of Covid-19$^{Mpro}$ with compounds EN82 and PM-02-20B was performed with 10mg/ml Covid-19$^{Mpro}$ (25 mM Hepes pH 7.5, 150 mM NaCl, 1 mM EDTA) was inhibited at 10X molar excess and incubated on ice for 1hr, solution was spun down for 10min at 10,000 rpm. Crystals were grown by hanging-drop vapor diffusion method at 18℃ by mixing 1:1, 1:2 and 2:1 ratio of protein to well solution. Crystal grew out of well solution composed of 25% w/v Peg 1500, 100 mM MIB buffer pH 7.0, from PACT screen (Nextal Biotechnologies). After 24 hr, crystals were harvested and cryo-protected using 20% Glycerol and well solution, flash-cooled in liquid nitrogen for data collection. EN82 was collected using in-house radiation source using an R-AXIS detector (Rigaku) and Cu Kα X-ray source (FR-E SuperBright High-Brilliance Rotating Anode Generator). Data was collected on a single crystal cooled to 100K. For PM-2-20B, data collection was completed at APS, IMCA-CAT beamline 17-ID-B.

**X-ray data collection, processing and structure refinement: EN82 PM-2-20b**

The diffraction images for both structures were processed using autoPROC, [Vonrhein, C., Flensburg, C., Keller, P., Sharff, A., Smart, O., Paciorek, W., Womack, T. & Bricogne, G. (2011). Data processing and analysis with the autoPROC toolbox. Acta Cryst. D67, 293-302.]. The Molecular Replacement solution was solved with Phaser (as implemented in CCP4I) using an in-house structure as the input model. This initial structure was built and refined by iterative cycles of manual rebuilding and subsequent structure refinement in Coot and autoBuster, respectively. Ligands were then placed and the structures further refined to convergence.

**Table 1** Data collection, processing and refinement statistics.

| | **EN-82 R stereoisomer** | **PM-20-20b** |
|---|---|---|
| PDB ID | | |
| **Data Collection** | | |
| Resolution range | 56.1 - 2.00 (2.10 - 2.00) | 48.31 - 1.88 (1.98-1.88) |
| Space group | C 1 2 1 | C 1 2 1 |
| Mol. in the ASU | 1 | 1 |
| Unit cell | 114.51 53.74 45.59 90.00 101.53 90.00 | 114.662 53.531 44.962 90.00 101.89 90.00 |
| Total reflections | 120409 (8386) | 63010 (2506) |
| Unique reflections | 17088 (1652) | 18050 (1065) |
| Multiplicity | 7.0 (5.1) | 3.5 (2.4) |
| Completeness (%) | 92.2 (62.1) | 82.7 (33.5) |
| Mean I/sigma(I) | 15.5 (1.3) | 18.6 (1.3) |
| Wilson B-factor | 39.19 | 36.09 |
| R-merge | 0.072 (0.967) | 0.032 (0.573) |
| R-meas | 0.084 (1.237) | 0.044 (0.811) |
| R-pim | 0.044 (0.755) | 0.030 (0.573) |
| CC1/2 | 0.999 (0.442) | 0.999 (0.623) |
| CC* | | |
| **Refinement** | | |
| Reflections used in refinement | 17064 (388) | 18025 (392) |
| Reflections used for R-free | 890 (27) | 929 (22) |
| R-work | 0.1936 (0.3067) | 0.1844 (0.3120) |
| R-free | 0.2359 (0.2941) | 0.2307 (0.3530) |
| CC(work) | 0.958 | 0.96 |
| CC(free) | 0.943 | 0.943 |
| Number of non-hydrogen atoms | 2467 | 2535 |
| protein atoms | 2310 | 2335 |
| solvent | 128 | 174 |
| Protein residues | 306 | 306 |
| RMS(bonds) | 0.008 | 0.008 |
| RMS(angles) | 0.95 | 0.99 |
| Ramachandran favored (%) | 99.01 | 99.01 |
| Ramachandran allowed (%) | 0.66 | 0 |
| Ramachandran outliers (%) | 0.33 | 0.99 |
| Rotamer outliers (%) | 0.4 | 0 |
| Clashscore | 4.37 | 5.82 |
| Average B-factor | 42.22 | 37.04 |

**Gel-Based ABPP Screens**

Recombinant Mpro (100 nM) was pre-treated with either DMSO vehicle or covalent ligand at room temperature for 30 min in 25 µL of PBS, and subsequently treated with Tetramethylrhodamine-5-iodoacetamide dihydroiodide (IA-rhodamine) (500 nM) (ThermoFisher Scientific) at room temperature for 1 h. The reaction was stopped by addition of 4×reducing Laemmli SDS sample loading buffer (Alfa Aesar). After boiling at 95 °C for 5 min, the samples were separated on precast 4−20% Criterion TGX gels (Bio-Rad). Probe-labeled proteins were analyzed by in-gel fluorescence using a ChemiDoc MP (Bio-Rad).

**SARS CoV2 MPro Activity Assay using a Fluorescent Substrate Peptide Probe (FRET-based assay)**

Compounds were made up in DMSO to 50X the desired screening concentration. DMSO was used as a solvent control. MPro protein was diluted in assay buffer (Tris buffered saline with 1 mM EDTA) to a concentration of 115 nM and was aliquoted to each well of a 96-well plate. Each well was treated with compound or vehicle and the plate was incubated for 30 min at room temperature. During the compound incubation the quenched fluorescent peptide probe (7-methoxycoumarin-4-ylacetyl) MCA-ABLQSGFR-Lys(2,4,-dinitrophenyl (Dnp))-Lys-NH was added from a 80 µM stock solution to a final concentration of 10 µM. Values were read-out on a Tecan Spark plate-reader.

**SARS CoV2 MPro Activity Assay using a Fluorescent Substrate Peptide Probe (Rhodamine-based assay)**

Compounds were made up in DMSO to 50X the desired screening concentration. DMSO was used as a solvent control. MPro protein was diluted in assay buffer (50 mM HEPES, pH 7.5, 150 mM NaCl, 1 mM EDTA, 0,01% pluronic acid F127) to a concentration of 30 nM and 24.5 µL of diluted protein was aliquoted to each well of a black 384 well plate (Corning 384-Well, Flat-Bottom Microplate). Each well was treated with 0.5 µL of compound or vehicle and the plate was incubated for 30 min at room temperature. During the compound incubation the peptide probe KTSAVLQ-(Rhodamine-110 (Rh-110))-gammaGlu (Biosyntan) was diluted from 5mM DMSO stock into assay buffer. After pre-incubation 5 µL of 75 µM Rh-110 probe was added to each well. RFU value was immediately measured on a Tecan Spark plate reader with an excitation wavelength of 488 nm and an emission wavelength of 535 nm at 30 °C for 30 min.

**SARS CoV2 MPro Activity Assay using an Agilent RapidFire Mass Spectrometer (MS-based assay)**

Compound $IC_{50}$ values for the Coranavirus main protease (Mpro) panel were determined in a Rapidfire-Mass spectrometry (RFMS) assay using an Agilent Rapidfire 365 autosampler coupled to Sciex 6500 triplequad (QQQ) mass spectrometer. Initially, Mpro was diluted in assay buffer (50 mM HEPES, pH 7.3, 150 mM NaCl, 1 mM EDTA, 0.01% pluronic acid F127) and added to a 384 well plate where it was pre-incubated with compound for 15 minutes prior to addition of substrate. To initiate the reaction, substrate was added to the compound plate and incubated for between 2-3 hrs depending on the Mpro (see table) at RT. Conditions were kept similar for each Mpro and were based off original conditions established for CoV2. Final incubation time was chosen when 10% substrate turnover was observed. After incubation the reaction was quenched with 2% acetic acid solution and then submitted to the RFMS. Samples were loaded onto a C18 SPE cartridge (Agilent) with $H_2O$ with 0.1% formic acid at a flow rate of 1.5 mL/min. They were then eluted with 75:20:5 ACN:$H_2O$:IPA with 0.1% formic acid at flow rate of 1.0 mL/min. MRM (multiple reaction monitoring) transitions corresponding to the substrate and the related products, along with $^{13}C_3$ $^{15}N$ AVLQ (for signal normalization) were monitored and peaks integrated using Sciex Multiquant. Corresponding peptides (Vivitide) monitored for each Mpro are listed in the table.

| Panel protein | Substrate | Product | Final enzyme conc. (nM) | Final substrate conc. (µM) | Incubation time (h) |
|---|---|---|---|---|---|
| IBV | SRLQAGFKKL | SRLQ | 5 | 10 | 2.5 |
| HCoV-NL63 | STLQSGLKKM | STLQ | 10 | 10 | 3 |
| HCoV-229E | STLQAGLRKM | STLQ | 10 | 10 | 3 |
| PorCoV-HKU15 | TKLQAGIKILL | TKLQ | 5 | 10 | 2.5 |
| HCoV-OC43 | SFLQSGIVKM | SFLQ | 5 | 5 | 2 |
| SARS-CoV-2 | AVLQSGFRKM | AVLQ | 5 | 5 | 2 |
| SARS-CoV-1 | AVLQSGFRKM | AVLQ | 5 | 5 | 2.5 |
| HCoV-HKU1 | SFLQSGIVKM | SFLQ | 5 | 5 | 2 |

**Cysteine profiling via rapid covalent chemoproteomics**

SARS-CoV-2 Mpro was spiked into HEK293T lysate (200ul at 5ug/ul) with a final concentration of 1uM followed by treatment with DMSO or EN82 at 10 μM for 6h in triplicate. This was followed by treatment with acid-cleavable biotin-PEG4-DADPS-C6-iodoacetamide probe at 100 μM for 1h at room temperature. Excess biotin probe was removed by cleanup through Zeba 7K MWCO columns. Lysates were denatured with 200 μL 8M urea, reduced with 10 mM DTT for 15 minutes and alkylated with 55 mM iodoacetamide for 1h. The denatured, alkylated proteins were digested with 20μg LysC/trypsin (Promega) for 4h at 37C. After dilution to 1M urea, biotinylated peptides were enriched by incubating with 100 μL ultralink streptavidin agarose (Thermo) for 1h at RT on a rotator. Beads were then transferred to 1.2 μM filter plate and washed 5x with 1 mL 0.1% SDS, 5x 1ml PBS and 5x with 1 mL water. 300 uL 10% formic acid were added and the samples incubated for 1h at RT to cleave the DADPS linker. Cysteinyl peptides were collected by centrifugation and dried by speedvac. After resuspension in 40 μL 50 mM HEPES pH 8, 60 μL acetonitrile and 50 μL TMT reagent in acetonitrile (Thermo) were added and the samples were incubated for 1hr at room temperature. TMT-labeled samples were pooled and 10% of the total material directly analyzed by nanoLC-SPS- using an Easy-nLC 1200 high-performance liquid chromatography system (Thermo) interfaced with an Orbitrap Eclipse Tribrid mass spectrometer (Thermo). A Kasil-fritted trapping column (75 μm x 15 mm) packed with 5 um ReproSil-Pur 120 C18-AQ, was used together with a fused silica spraying capillary pulled to a tip diameter of 8-10 um using a P-2000 capillary puller (Sutter Instruments). The capillary tubing (75 μm I.D.) was packed with a 120 mm separation column comprised of 3 um ReproSil-Pur C18 AQ. Samples (10 μL) were injected onto the trapping column using 0.1% formic acid/2% acetonitrile in water at a flow rate of 2.5 μL/min. Trapped peptides were then introduced into the separation column and eluted at 300 nL/min using a gradient of 3-45% mobile phase B (98% acetonitrile + 0.1% formic acid in water) over 90 min (mobile phase A: 2% acetonitrile + 0.1% formic acid in water). SPS-MS3 analysis of the TMT labeled peptides was performed using MS1 scans that were acquired from m/z 400-1600 at 120,000 mass resolution with a triggering intensity threshold of 400,000 ions. MS2 scans were acquired using Turbo CID with 1.2 Da isolation and AGC target of 1E4 and a collision energy of 30%. For SPS the top 10 ions were selected for MS3 analysis using a collision energy of 55% with an orbitrap resolution of 60,000. Raw files were processed using Proteome Discoverer 2.4. Data were searched against the Uniprot human protein database including the SARS-CoV-2 Mpro sequence using Mascot.

**General Synthetic Methods**

All non-aqueous reactions were performed under an inert atmosphere of dry nitrogen in flame dried glassware sealed with a rubber septum unless stated otherwise. Nitrogen was supplied through a glass manifold. Reactions were stirred magnetically and monitored by thin layer chromatography (TLC). Analytical thin layer chromatography was performed using MERCK Silica Gel 60 F254 TLC glass plates and visualized by ultraviolet light (UV). Additionally, TLC plates were stained with aqueous potassium permanganate ($KMnO_4$) [1.5 g $KMnO_4$, 200 mL $H_2O$, 10 g $K_2CO_3$, 1.25 mL 10% NaOH]. Concentration under reduced pressure was performed by rotator evaporation at 40 °C at the appropriate pressure. Chromatographic purification was performed as flash chromatography on MERCK silica gel 60 Å (230 x 400 mesh) at 0.2–0.5 bar overpressure. Purified compounds were dried further under high vacuum (0.01-0.1 mbar). Yields refer to the purified compound.

**Chemicals**

All chemicals and solvents were used as received from the commercial supplier without further purification unless mentioned otherwise. DCM was purified by passage through an activated alumina column under an atmosphere of dry argon.

**Analytics**

Nuclear Magnetic Resonance (NMR) spectra were recorded on BRUKER AV (600 MHz and 300 MHz), AVB (400 MHz), AVQ (400 MHz) and NEO (500 MHz) spectrometers. Measurements were carried out at ambient temperature. Chemical shifts ($\delta$) are reported in ppm with the residual solvent signal as internal standard (chloroform at 7.26 and 77.00 ppm for $^{1}$H NMR and $^{13}$C NMR spectroscopy, respectively). The data is reported as (s = singlet, d = doublet, t = triplet, q = quartet, p = quintet, m = multiplet or unresolved, br = broad signal, coupling constant(s) in Hz, integration). $^{13}$C NMR spectra were recorded with broadband $^{1}$H decoupling.

Mass spectrometry (MS) analyses were obtained at the Catalysis Center at the College of Chemistry, University of California, Berkeley.

### General Procedures

#### Synthesis of α,β-Unsaturated Ketones

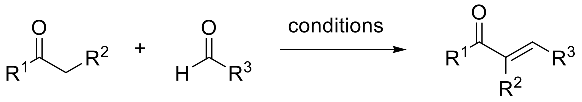

#### General Procedure A

The ketone (1.0 equiv) and aldehyde (2.0 equiv) were taken up in EtOH (0.6 M) and cooled to 0 °C. 40% NaOH (10 equiv) was added dropwise, and the resulting reaction mixture was stirred at ambient temperature for 4 h. The precipitated solid was filtered, washed with cold water and cold EtOH and dried under high vacuum. Where necessary, the crude product was recrystallized from hot EtOH to afford the corresponding α,β-unsaturated ketone.

#### General Procedure B

$LiOH \cdot H_2O$ (10 mol%) was added to a solution of the ketone (1.0 equiv) in EtOH (1 M) and the resulting mixture was stirred for 10 min at ambient temperature. The aldehyde (1.0 equiv) was then added, and the reaction mixture was stirred at ambient temperature for 30 min to 1 h before it was concentrated under reduced pressure. Purification by column chromatography afforded the corresponding the corresponding α,β-unsaturated ketone.

#### General Procedure C

A mixture of the aldehyde (1.0 equiv) and deoxybenzoin (3.0 equiv) in toluene (0.4 M) was treated with acetic acid (0.9 equiv) and piperidine (0.2 equiv). Powdered 4 Å molecular sieves (500 mg/mmol) was added, and the resulting reaction mixture was stirred at reflux temperature for 7 to 16 h. It was then allowed to cool to ambient temperature, quenched with $NaHCO_3$ solution (sat. aqueous) and extracted with EtOAc (3 x). The combined organic phases were washed with NaCl solution (sat. aqueous), dried over $Na_2SO_4$, filtered and concentrated under reduced pressure. Purification by column chromatography afforded the corresponding α,β-unsaturated ketone.

#### General Procedure D

A mixture of the aldehyde (1.0 equiv) and propiophenone (1.0 equiv) in EtOH (0.3 M) was heated gently until both starting materials dissolved. A solution of NaOH (1.2 equiv) in EtOH/$H_2O$ (1:1 v/v, 0.3 M) was added dropwise, and the resulting reaction mixture was stirred at ambient temperature for 1 h, then at 60 °C for 16 h. The reaction mixture was allowed to cool to ambient temperature, diluted with water and extracted with DCM (3 x). The combined organic phases were washed with NaCl solution (sat. aqueous), dried over $Na_2SO_4$, filtered and concentrated under reduced pressure. Purification by column chromatography afforded the corresponding α,β-unsaturated ketone.

#### General Procedure E

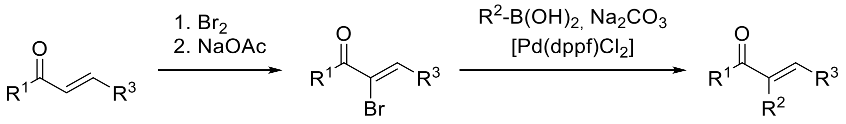

A solution of the chalcone (1.0 equiv) in DCM (0.1 M) was cooled to 0 °C. A solution of bromine (1.2 equiv) in DCM (1 M) was added dropwise over 3 min, and the resulting reaction mixture was stirred at 0 °C for 15 min, then at ambient temperature for 15 min. It was then quenched with $Na_2S_2O_3$ solution (sat. aqueous) and the resulting mixture was extracted with DCM (3 x). The combined organic phases were washed with NaCl solution (sat. aqueous), dried over $Na_2SO_4$, filtered and concentrated under reduced pressure.

The crude dibromide was dissolved in EtOH (0.3 M) and sodium acetate (1.2 equiv) was added. The resulting reaction mixture was stirred at reflux temperature for 1.5 h before the solvent was removed under reduced pressure. The residue was partitioned between DCM and $NaHCO_3$ solution (sat. aqueous), the phases were separated and the aqueous phase was extracted with DCM (2 x). The combined organic phases were washed with NaCl solution (sat. aqueous), dried over $Na_2SO_4$, filtered and concentrated under reduced pressure.

A mixture of the crude vinyl bromide (1.0 equiv), the boronic acid (1.3 equiv), sodium carbonate (2.0 equiv) and $Pd(dppf)Cl_2$ (5 mol%) was taken up in toluene/$H_2O$ (4:1 v/v, 0.2 M). The resulting suspension was sparged with nitrogen for 10 min. The flask was sealed and the reaction mixture was stirred at 100 °C for 1 h. It was then allowed to cool to ambient temperature and partitioned between DCM and 1 M NaOH. The phases were separated and the aqueous phase was extracted with DCM (2 x). The combined organic phases were washed with NaCl solution (sat. aqueous), dried over $Na_2SO_4$, filtered and concentrated under reduced pressure. Purification by column chromatography afforded the corresponding α,β-unsaturated ketone.

**Synthesis of Pyrazoline Chloroacetamides**

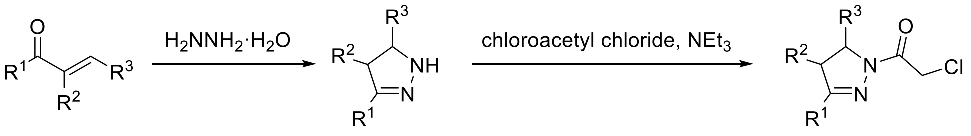


**General Procedure F**

Hydrazine monohydrate (2.0 equiv) was added to a suspension of the α,β-unsaturated ketone (1.0 equiv) in EtOH (0.3 M). The resulting reaction mixture was stirred at reflux temperature for 2.5 to 4 h before it was concentrated under reduced pressure. The crude pyrazoline was then dissolved in DCM (0.2 M) and cooled to 0 °C. Triethylamine (3.0 equiv) was added dropwise, followed by chloroacetyl chloride (1.5 equiv). The resulting reaction mixture was stirred at ambient temperature for 30 min before it was diluted with DCM. The organic phase was sequentially washed with $NaHCO_3$ solution (sat. aqueous) and NaCl solution (sat. aqueous), dried over $Na_2SO_4$, filtered and concentrated under reduced pressure. Purification by column chromatography afforded the corresponding chloroacetamide.

**Synthesis of Pyrazoline Vinyl Sulfonamides**

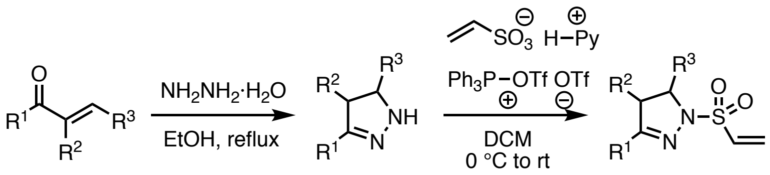


**General Procedure G**

*Step 1*: Hydrazine monohydrate (2.0 equiv) was added to a suspension of the α,β-unsaturated ketone (1.0 equiv) in EtOH (0.3 M) at rt. The reaction mixture was then stirred at reflux for 2h, then allowed to cool to rt and concentrated under reduced pressure to give the crude pyrazoline. This was used in the next step without further purification.

*Step 2*: A solution of triphenylphosphine oxide (2.2 equiv) in dichloromethane (0.25 M) was degassed at rt. Trifluoromethanesulfonic anhydride (1.0 equiv) was added at rt and the resultant mixture was stirred at this temperature for 15 min. In a separate flask, pyridine (1.0 equiv) was added to a solution of vinyl sulfonic acid (1.0 equiv) in dichloromethane (0.15 M) at rt and the mixture was then concentrated *in vacuo* to give a white solid. A solution of the vinyl sulfonic acid-pyridinium salt in dichloromethane (0.30 M) was added to the reaction mixture at rt and the resultant mixture was stirred at this temperature for 30 min. A solution of the crude pyrazoline (1.0 equiv) and triethylamine (2.0 equiv) in dichloromethane (0.20 M) was then added to the reaction mixture at 0 °C and the resultant mixture was allowed to warm to rt and stirred at rt for 16 h. The mixture was then diluted with dichloromethane and washed sequentially with 2.0 M aq HCl, sat aq $NaHCO_3$ and NaCl solution (sat. aqueous), then dried and concentrated under reduced pressure. Purification *via* flash column chromatography using both normal-phase (gradient elution, EtOAc in hexane) and reverse-phase (gradient elution, acetonitrile in water) conditions afforded the vinyl sulfonamides.

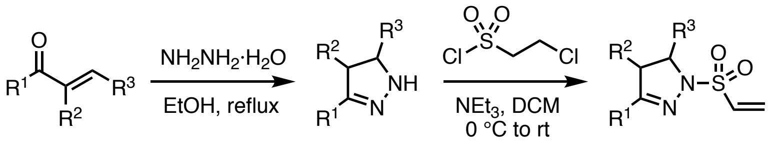


**General Procedure H**

Hydrazine monohydrate (2.0 equiv) was added to a suspension of the α,β-unsaturated ketone (1.0 equiv) in EtOH (0.3 M). The resulting reaction mixture was stirred at reflux temperature for 1 to 4 h before it was concentrated under reduced pressure. The crude pyrazoline. The crude pyrazoline was dissolved in DCM (0.2 M) and cooled to 0 °C. Triethylamine (3.0 equiv) was added dropwise, followed by 2-chloroethanesulfonyl chloride (1.2–1.5 equiv). The resulting reaction mixture was stirred at room temperature 1 h before it was diluted with DCM. The organic phase was sequentially washed with 1M HCl, sat. aq. $NaHCO_3$ and NaCl solution (sat. aqueous), dried over $Na_2SO_4$, filtered and concentrated under reduced pressure. Purification by column chromatography or preparative TLC afforded the corresponding vinyl sulfonamide.

**Stereoselective Synthesis of Pyrazoline Chloroacetamides**

**General Procedure I**

(*E*)-1-phenyl-3-(quinoxalin-5-yl)prop-2-en-1-one (1.0 equiv), tert-butyl carbazate (1.1 equiv), potassium phosphate (1.3 equiv), indicated catalyst (10 mol%) were combined in THF (0.5 M) under nitrogen and stirred at 0 °C for 18h. The mixture was diluted with EtOAc, filtered to remove salts, concentrated and purified by silica gel chromatography to provide the stereoenriched pyrazoline.

**General Procedure J**

Boc-protected pyrazoline (1.0 eq) was dissolved in DCM (0.2 M), cooled to 0 °C, and 4.0 M HCl in dioxane (10.0 equiv) was added under nitrogen. The mixture was stirred at room temperature for several hours (2-3) until starting material was consumed, then additional DCM was added (reducing HCl to 0.1 M). At 0 °C, chloroacetyl chloride (3.0 equiv) was added followed by triethylamine (13.0 equiv), and the reaction allowed to warm to room temperature and stirred for 1h. Water was added and the mixture extracted with DCM. Extracts were combined, washed with NaCl solution (sat. aqueous), dried over $Na_2SO_4$, concentrated under vacuum and purified by silica gel chromatography.

**Formylation of Aryl Bromides**

**General Procedure K**

Aryl bromide (1.0 Equiv) was dissolved in THF (to 0.2 M) and cooled to -78 ºC. n-BuLi (2.5 M in hexanes) was added dropwise, the reaction turned red-brown, and was stirred for 20 minutes at -78 ºC. Then, DMF (2.0 Equiv) was added dropwise. The reaction was removed from the -78 ºC bath and allowed to stir for 2h at room temperature. To quench the reaction the flask was cooled to -78 ºC quickly, sat. $NH_4Cl$ (2 mL) was added dropwise, and the reaction allowed to warm to rt again. Water was added and the mixture was extracted with ethyl acetate. Combined organic extracts were washed with NaCl solution (sat. aqueous), dried over sodium sulfate, concentrated, and purified by silica gel chromatography to provide the resulting aldehyde.

**Characterization Data**

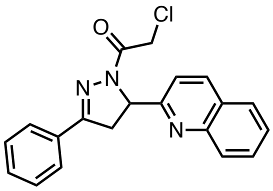

**HW-01-146**

**2-Chloro-1-(3-phenyl-5-(quinolin-2-yl)-4,5-dihydro-1*H*-pyrazol-1-yl)ethan-1-one.** General Procedure B was followed starting from acetophenone (0.30 mL, 2.6 mmol) and quinoline-2-carbaldehyde (445 mg, 2.83 mmol). Purification by column chromatography (EtOAc/hexane, 15:85) afforded the corresponding chalcone (326 mg) as a white solid.

General Procedure F was followed starting from the above product (0.20 g, 0.77 mmol). Purification by column chromatography (EtOAc/hexane, 30:70) afforded the corresponding chloroacetamide (158 mg, 29% over three steps) as a white solid.

**$^1$H NMR** (400 MHz, $CDCl_3$): $\delta$ 8.15 (d, $J$ = 8.5 Hz, 1H), 8.01 (dd, $J$ = 8.5, 1.0 Hz, 1H), 7.80 (ddd, $J$ = 6.9, 4.0, 1.6 Hz, 3H), 7.68 (ddd, $J$ = 8.4, 6.9, 1.5 Hz, 1H), 7.55–7.49 (m, 2H), 7.49–7.43 (m, 3H), 5.88 (dd, $J$ = 10.9, 6.1 Hz, 1H), 4.64 (d, $J$ = 2.1 Hz, 2H), 3.89 – 3.71 (m, 2H); **$^{13}$C NMR** (151 MHz, $CD_2Cl_2$) δ 164.5, 159.6, 156.5, 148.3, 137.4, 131.3, 131.1, 130.1, 129.5, 129.2, 128.0, 128.0, 127.3, 127.0, 119.9, 62.7, 42.9, 40.6. **HRMS** (ESI): exact mass calculated for $C_{20}H_{17}ClN_3O$ [(M+H)$^+$] 350.1055, found 350.1048.

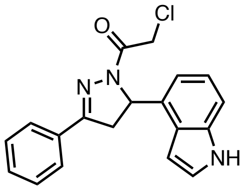

**HW-01-151**

**1-(5-(1*H*-Indol-4-yl)-3-phenyl-4,5-dihydro-1*H*-pyrazol-1-yl)-2-chloroethan-1-one.** General Procedure B was followed starting from acetophenone (0.30 mL, 2.6 mmol) and indole-4-carbaldehyde (373 mg, 2.57 mmol). Purification by column chromatography (EtOAc/hexane, 30:70) afforded the corresponding chalcone (375 mg) as a yellow solid.

General Procedure F was followed starting from the above product (0.20 g, 0.81 mmol). Purification by column chromatography (EtOAc/hexane, 30:70) afforded the corresponding chloroacetamide (100 mg, 20% over three steps) as a yellowish solid.

**$^1$H NMR** (600 MHz, $CDCl_3$): $\delta$ 8.27 (s, br, 1H), 7.83–7.71 (m, 2H), 7.52–7.39 (m, 3H), 7.32 (dt, $J$ = 8.2, 1.0 Hz, 1H), 7.19 (dd, $J$ = 3.3, 2.5 Hz, 1H), 7.14 (t, $J$ = 7.7 Hz, 1H), 7.02 (d, $J$ = 7.2 Hz, 1H), 6.43 (ddd, $J$ = 3.2, 1.9, 0.9 Hz, 1H), 5.94 (dd, $J$ = 11.9, 5.4 Hz, 1H), 4.74–4.51 (m, 2H), 3.85 (dd, $J$ = 17.8, 12.0 Hz, 1H), 3.38 (dd, $J$ = 17.8, 5.4 Hz, 1H); **$^{13}$C NMR** (151 MHz, $CDCl_3$): $\delta$ 164.0, 155.8, 136.4, 132.0, 131.0, 130.7, 128.8, 126.8, 124.4, 124.3, 122.1, 117.4, 111.0, 100.2, 59.9, 42.3, 41.5; **HRMS** (ESI): exact mass calculated for $C_{19}H_{17}ClN_3O$ [(M+H)$^+$] 338.1055, found 338.1026.

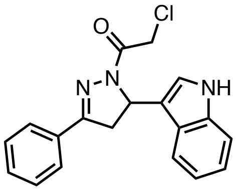


**HW-01-161**

**1-(5-(1*H*-Indol-3-yl)-3-phenyl-4,5-dihydro-1*H*-pyrazol-1-yl)-2-chloroethan-1-one.** General Procedure B was followed starting from acetophenone (0.10 mL, 0.86 mmol) and indole-3-carbaldehyde (124 mg, 0.857 mmol). The stirring time was prolonged to 22 h at ambient temperature followed by 1 h at 40 °C. Purification by column chromatography (EtOAc/hexane, 35:65) afforded the corresponding chalcone (22 mg) as a yellow solid.

General Procedure F was followed starting from the above product (22 mg, 89 µmol). Purification by column chromatography (EtOAc/hexane, 40:60) afforded the corresponding chloroacetamide (19 mg, 6% over three steps) as a white solid.

**$^1$H NMR** (600 MHz, $CDCl_3$): $\delta$ 8.48 (s, br, 1H), 7.84–7.77 (m, 2H), 7.53–7.43 (m, 3H), 7.34 (ddd, $J$ = 8.9, 8.0, 1.0 Hz, 2H), 7.20–7.07 (m, 2H), 7.01 (ddd, $J$ = 8.0, 7.0, 1.0 Hz, 1H), 5.92 (dd, $J$ = 11.8, 4.8 Hz, 1H), 4.63 (d, $J$ = 13.8 Hz, 1H), 4.57 (d, $J$ = 13.8 Hz, 1H), 3.76 (dd, $J$ = 17.8, 11.8 Hz, 1H), 3.48 (dd, $J$ = 17.8, 4.8 Hz, 1H); **$^{13}$C NMR** (151 MHz, $CDCl_3$): $\delta$ 164.1, 156.1, 136.9, 131.0, 130.8, 128.9, 126.9, 124.2, 123.5, 122.2, 119.8, 118.5, 114.6, 111.8, 54.6, 42.5, 40.4; **HRMS** (ESI): exact mass calculated for $C_{19}H_{17}ClN_3O$ [(M+H)$^+$] 338.1055, found 338.1032.

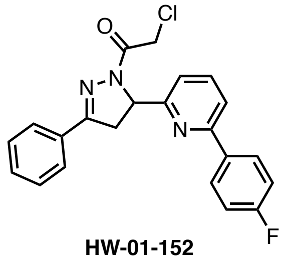


**HW-01-152**

**2-Chloro-1-(5-(6-(4-fluorophenyl)pyridin-2-yl)-3-phenyl-4,5-dihydro-1*H*-pyrazol-1-yl)-ethan-1-one.** General Procedure B was followed starting from acetophenone (50 µL, 0.43 mmol) and 6-(4-fluorophenyl)picolinaldehyde (86 mg, 0.43 mmol). Purification by column chromatography (EtOAc/hexane, 15:85) afforded the corresponding chalcone (108 mg) as a white solid.

General Procedure F was followed starting from the above product (60 mg, 0.20 mmol). Purification by column chromatography (EtOAc/hexane, 30:70) afforded the corresponding chloroacetamide (57 mg, 61% over three steps) as a white solid.

**$^1$H NMR** (400 MHz, $CDCl_3$): $\delta$ 7.98–7.88 (m, 2H), 7.84–7.76 (m, 2H), 7.72 (t, $J$ = 7.7 Hz, 1H), 7.59 (d, $J$ = 7.8 Hz, 1H), 7.51–7.40 (m, 3H), 7.32 (d, $J$ = 7.6 Hz, 1H), 7.08 (t, $J$ = 8.7 Hz, 2H), 5.76 (dd, $J$ = 10.9, 5.5 Hz, 1H), 4.63 (d, $J$ = 1.1 Hz, 2H), 3.84–3.63 (m, 2H); **$^{13}$C NMR** (151 MHz, $CD_2Cl_2$) δ 164.4, 164.0 (d, $J$ = 247.8 Hz), 158.8, 156.6, 156.4, 138.1, 135.6 (d, $J$ = 3.2 Hz), 131.5, 131.1, 129.2, 129.1 (d, $J$ = 8.4 Hz), 127.3, 120.5, 119.5, 115.8 (d, $J$ = 21.6 Hz), 62.1, 42.9, 40.7. **HRMS** (ESI): exact mass calculated for $C_{22}H_{18}ClFN_3O$ [(M+H)$^+$] 394.1117, found 394.1099.

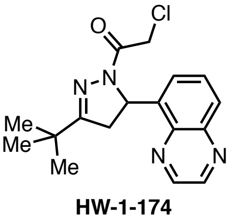


**HW-1-174**

**1-(3-(*tert*-Butyl)-5-(quinoxalin-5-yl)-4,5-dihydro-1*H*-pyrazol-1-yl)-2-chloroethan-1-one.** General Procedure A was followed starting from pinacolone (20 µL, 0.16 mmol) and quinoxaline-5-carbaldehyde (50 mg, 0.32 mmol). The crude α,β-unsaturated ketone (27 mg) was used in the following transformation without further purification.
General Procedure F was followed starting from the above product (26 mg, 0.11 mmol). Purification by column chromatography (EtOAc/hexane, 40:60) afforded the corresponding chloroacetamide (18 mg, 35% over three steps) as a white solid.
**$^1$H NMR** (300 MHz, $CDCl_3$): *δ* 8.88 (d, *J* = 1.8 Hz, 1H), 8.83 (d, *J* = 1.8 Hz, 1H), 8.04 (d, *J* = 8.6 Hz, 1H), 7.81–7.66 (m, 1H), 7.48 (d, *J* = 7.1 Hz, 1H), 6.46 (dd, *J* = 11.7, 4.9 Hz, 1H), 4.69–4.43 (m, 2H), 3.63 (dd, *J* = 18.1, 11.7 Hz, 1H), 2.79 (dd, *J* = 18.1, 5.0 Hz, 1H), 1.20 (s, 9H); **$^{13}$C NMR** (151 MHz, $CD_2Cl_2$) δ 168.4, 164.1, 145.6, 144.4, 143.9, 140.5, 139.5, 130.2, 129.5, 125.9, 56.91, 42.8, 42.0, 34.5, 28.1. **HRMS** (ESI): exact mass calculated for $C_{17}H_{20}ClN_4O$ [$(M+H)^+$] 331.1320, found 331.1343.

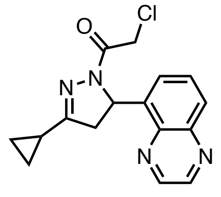

**HW-01-185**

**2-Chloro-1-(3-cyclopropyl-5-(quinoxalin-5-yl)-4,5-dihydro-1*H*-pyrazol-1-yl)ethan-1-one.** General Procedure A was followed starting from cyclopropyl methyl ketone (20 µL, 0.20 mmol) and quinoxaline-5-carbaldehyde (64 mg, 0.40 mmol). Upon completion of the reaction, the reaction mixture was diluted with water (10 mL) and extracted with diethyl ether (3 x 20 mL). The combined organic phases were washed with NaCl solution (20 mL, sat. aqueous), dried over $Na_2SO_4$, filtered and concentrated under reduced pressure. Purification by column chromatography (EtOAc/hexane, 30:70) afforded the corresponding α,β-unsaturated ketone (53 mg) as a white solid.
General Procedure F was followed starting from the above product (23 mg, 0.10 mmol). Purification by column chromatography (EtOAc/hexane, 50:50) afforded the corresponding chloroacetamide (16 mg, 50% over three steps) as a white solid.
**$^1$H NMR** (300 MHz, $CDCl_3$): *δ* 8.87 (d, *J* = 1.8 Hz, 1H), 8.82 (d, *J* = 1.8 Hz, 1H), 8.03 (dd, *J* = 8.5, 1.4 Hz, 1H), 7.73 (dd, *J* = 8.5, 7.2 Hz, 1H), 7.56–7.42 (m, 1H), 6.47 (dd, *J* = 11.7, 4.9 Hz, 1H), 4.69–4.36 (m, 2H), 3.49 (dd, *J* = 18.0, 11.6 Hz, 1H), 2.57 (dd, *J* = 18.0, 4.9 Hz, 1H), 1.81 (ddd, *J* = 13.3, 8.4, 5.0 Hz, 1H), 1.04–0.66 (m, 4H); **$^{13}$C NMR** (151 MHz, $CD_2Cl_2$) δ 163.6, 163.5, 145.6, 144.4, 143.9, 140.4, 139.3, 130.2, 129.5, 125.8, 56.3, 42.9, 42.7, 11.7, 7.4, 7.0. **HRMS** (ESI): exact mass calculated for $C_{16}H_{15}ClN_4NaO$ [$(M+MeCN+Na)^+$] 378.1082, found 378.1055.

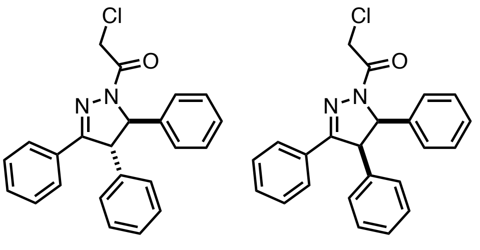

**PM-02-20A** **PM-02-20B**

**2-chloro-1-(3,4,5-triphenyl-4,5-dihydro-1*H*-pyrazol-1-yl)ethan-1-one.** General Procedure C was followed starting from deoxybenzoin (235 mg, 1.2 mmol) and benzaldehyde (0.12 mL, 1.2 mmol). Purification by column chromatography (EtOAc/hexane, 10:90) afforded the corresponding α,β-unsaturated ketone (126 mg) as a mixture with benzaldehyde. This mixture was used in the following transformation without further purification.

General Procedure F was followed starting from the above product (120 mg, 0.42 mmol). Purification by column chromatography (EtOAc/hexane, 0:100 to 20:80) afforded the *trans*-substituted chloroacetamide **PM-02-20A** (13 mg, 2% over three steps) as a colorless film and the *cis*-substituted chloroacetamide **PM-02-20B** (8 mg, 1% over three steps) as a colorless film.
**PM-02-20A [trans] : $^1$H NMR** (300 MHz, $CDCl_3$) δ 7.71 – 7.63 (m, 2H), 7.40 – 7.27 (m, 9H), 7.25 – 7.10 (m, 4H), 5.36 (d, *J* = 3.6 Hz, 1H), 4.68 (d, *J* = 2.7 Hz, 2H), 4.55 (d, *J* = 3.5 Hz, 1H). **$^{13}$C NMR** (126 MHz, $CDCl_3$) δ 164.2, 157.2, 140.2, 139.3, 130.6, 130.2, 129.7, 129.3, 128.8, 128.3, 128.2, 127.6, 127.1, 125.5, 70.7, 61.5, 42.3. **HRMS** (ESI): exact mass calculated for $C_{23}H_{20}ClN_2O$ [$(M+H)^+$] 375.1259, found 375.1253.

**PM-02-20B [cis]:** **$^1$H NMR** (300 MHz, $C_6D_6$) δ 7.56 (dd, *J* = 6.9, 2.8 Hz, 2H), 6.96 (t, *J* = 3.4 Hz, 3H), 6.89 – 6.69 (m, 5H), 6.69 – 6.60 (m, 3H), 6.45 (dd, *J* = 6.3, 2.7 Hz, 2H), 5.46 (d, *J* = 11.8 Hz, 1H), 4.54 – 4.40 (m, 2H), 4.34 (d, *J* = 11.7 Hz, 1H). **$^{13}$C NMR** (151 MHz, $C_6D_6$) δ 164.4, 155.8, 136.2, 135.3, 131.2, 130.1, 129.9, 128.6, 127.2, 127.2, 127.2, 66.9, 57.2, 42.2. **HRMS** (ESI): exact mass calculated for $C_{23}H_{20}ClN_2O$ [(M+H)$^+$] 375.1259, found 375.1280.

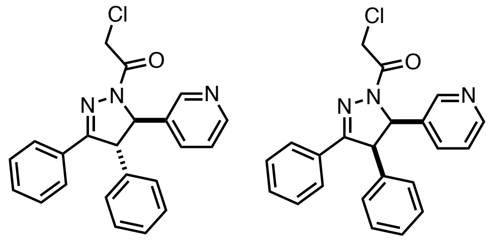

**PM-02-43A** **PM-02-43B**

**2-chloro-1-(3,4-diphenyl-5-(pyridin-3-yl)-4,5-dihydro-1*H*-pyrazol-1-yl)ethan-1-one.** General procedure E was followed starting from 1-phenyl-3-(pyridin-3-yl)prop-2-en-1-one (630 mg, 3.0 mmol, 1 equiv.). The intermediate product 2-bromo-1-phenyl-3-(pyridin-3-yl)prop-2-en-1-one was obtained as a light orange oil (896 mg, 99%). **$^1$H NMR** (300 MHz, $CDCl_3$) δ 8.86 (d, *J* = 2.3 Hz, 1H), 8.65 (dd, *J* = 4.8, 1.6 Hz, 1H), 8.38 (dt, *J* = 7.8, 1.9 Hz, 1H), 7.87 – 7.80 (m, 2H), 7.65 (s, 1H), 7.65 – 7.58 (m, 1H), 7.51 (m, 2H), 7.46 – 7.37 (m, 1H).

A mixture of 2-bromo-1-phenyl-3-(pyridin-3-yl)prop-2-en-1-one (58 mg, 0.20 mmol, 1.0 equiv), phenylboronic acid (32 mg, 0.26 mmol, 1.3 equiv), sodium carbonate (43 mg, 0.40 mmol, 2.0 equiv) and Pd(dppf)$Cl_2$ (7 mg, 10 µmol, 5 mol%) was taken up in toluene/$H_2O$ (4:1 v/v, 1 mL). The resulting suspension was sparged with nitrogen for 10 min. The flask was sealed and the reaction mixture was stirred at 100 °C for 1 h. The reaction mixture was then allowed to cool to ambient temperature and partitioned between DCM (20 mL) and 1 M NaOH (10 mL). The phases were separated and the aqueous phase was extracted with DCM (2 x 20 mL). The combined organic phases were washed with NaCl solution (20 mL, sat. aqueous), dried over $Na_2SO_4$, filtered and concentrated under reduced pressure. Purification by column chromatography (EtOAc/hexane, 40:60 to 100:0) afforded the corresponding phenyl-substituted α,β-unsaturated ketone (41 mg, 72%). **$^1$H NMR** (300 MHz, $CDCl_3$) δ 8.54 – 8.32 (m, 2H), 7.96 (dt, *J* = 8.4, 1.2 Hz, 1H), 7.85 (dt, *J* = 7.0, 1.3 Hz, 1H), 7.59 – 7.50 (m, 1H), 7.50 – 7.40 (m, 2H), 7.40 – 7.22 (m, 6H), 7.14 (d, *J* = 2.7 Hz, 1H), 7.06 (dd, *J* = 8.6, 4.3 Hz, 1H).

General Procedure F was followed starting from the above product (41 mg, 0.14 mmol). Purification by column chromatography (7% to 9% MeOH/DCM) afforded the *trans*-substituted chloroacetamide **PM-02-43A** (18 mg, 25% over three steps) as a colorless film. Further purification (EtOAc/hexane, 20:80 to 100:0) afforded the *cis*-substituted chloroacetamide **PM-02-43B** (8 mg, 11% over three steps) as a colorless film.

**PM-02-43A [trans] :** **$^1$H NMR** (600 MHz, $C_6D_6$) δ 8.61 (d, *J* = 2.3 Hz, 1H), 8.45 (dd, *J* = 4.8, 1.6 Hz, 1H), 7.56 – 7.48 (m, 2H), 7.11 (dt, *J* = 8.0, 2.0 Hz, 1H), 6.94 (ddt, *J* = 11.0, 8.6, 4.4 Hz, 6H), 6.83 – 6.77 (m, 2H), 6.66 (dd, *J* = 7.9, 4.7 Hz, 1H), 5.31 (d, *J* = 3.9 Hz, 1H), 4.39 – 4.26 (m, 2H), 4.19 (d, *J* = 4.0 Hz, 1H). **$^{13}$C NMR** (151 MHz, $C_6D_6$) δ 164.0, 156.2, 149.9, 148.2, 139.4, 136.2, 132.8, 130.5, 130.5, 129.9, 128.8, 128.3, 127.8, 127.2, 123.8, 68.9, 61.3, 41.8. **HRMS** (ESI): exact mass calculated for $C_{22}H_{19}ClN_3O$ [(M+H)$^+$] 376.1211, found 376.1196.

**PM-02-43B [cis] :** **$^1$H NMR** (500 MHz, $C_6D_6$) δ 8.29 (d, *J* = 2.5 Hz, 1H), 8.15 (dd, *J* = 4.8, 1.7 Hz, 1H), 7.56 – 7.48 (m, 2H), 7.04 – 6.89 (m, 3H), 6.78 (dt, *J* = 7.9, 2.1 Hz, 1H), 6.65 (dd, *J* = 5.2, 2.8 Hz, 3H), 6.44 – 6.34 (m, 3H), 5.30 (d, *J* = 11.6 Hz, 1H), 4.47 – 4.36 (m, 2H), 4.30 (d, *J* = 11.7 Hz, 1H). **$^{13}$C NMR** (126 MHz, $C_6D_6$) δ 164.8, 155.8, 149.1, 148.8, 134.9, 133.7, 131.7, 130.9, 130.3, 129.6, 128.7, 128.6, 128.6, 127.5, 122.5, 64.8, 56.9, 42.2. **HRMS** (ESI): exact mass calculated for $C_{22}H_{19}ClN_3O$ [(M+H)$^+$] 376.1211, found 376.1211.

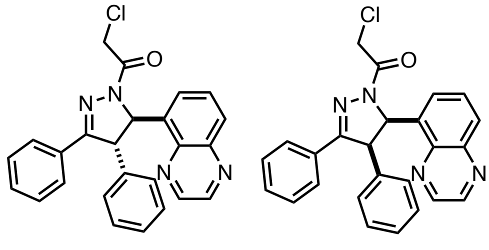

**PM-02-27A** **PM-02-27B**

**2-chloro-1-(3,4-diphenyl-5-(quinoxalin-5-yl)-4,5-dihydro-1*H*-pyrazol-1-yl)ethan-1-one.** General Procedure C was followed starting from deoxybenzoin (39 mg, 0.20 mmol) and quinoxaline-6-carbaldehyde (32 mg, 0.20 mmol). Purification by column chromatography (EtOAc/hexane, 1:4) afforded the corresponding α,β-unsaturated ketone (22 mg) as a mixture with quinoxaline-6-carbaldehyde. This mixture was used in the following transformation without further purification.

General Procedure F was followed starting from the above product (22 mg, 0.066 mmol). Purification by column chromatography (EtOAc/hexane, 0:100 to 40:60) afforded the *trans*-substituted chloroacetamide **PM-02-27A** (10 mg, 12% over three steps) as a colorless film and the *cis*-substituted chloroacetamide **PM-02-27B** (1.4 mg, 2% over three steps) as a colorless film.

**PM-02-27A [trans]:** $^{1}$**H NMR** (300 MHz, $CD_2Cl_2$) δ 8.87 (dd, *J* = 22.7, 1.8 Hz, 2H), 8.08 (d, *J* = 8.4 Hz, 1H), 7.81 – 7.61 (m, 3H), 7.51 (d, *J* = 7.3 Hz, 1H), 7.46 – 7.13 (m, 8H), 6.47 (d, *J* = 3.0 Hz, 1H), 4.92 – 4.71 (m, 2H), 4.61 (d, *J* = 3.1 Hz, 1H). $^{13}$**C NMR** (151 MHz, $CD_2Cl_2$) δ 164.3, 158.4, 145.8, 144.5, 144.1, 140.6, 139.5, 137.8, 130.9, 130.8, 130.2, 129.9, 129.6, 129.0, 128.3, 128.0, 127.8, 125.8, 66.8, 61.3, 42.7.**HRMS** (ESI): exact mass calculated for $C_{25}H_{20}ClN_4O$ [$(M+H)^+$] 427.1320, found 427.1293.

**PM-02-27B [cis]:** $^{1}$**H NMR** (300 MHz, $C_6D_6$) δ 8.26 – 8.16 (m, 2H), 7.71 (d, *J* = 8.4 Hz, 1H), 7.60 (dd, *J* = 6.8, 2.9 Hz, 1H), 7.24 (d, *J* = 7.2 Hz, 1H), 7.05 – 6.87 (m, 5H), 6.43 (m, 4H), 5.00 (d, *J* = 11.7 Hz, 1H), 4.68 (d, *J* = 12.8 Hz, 1H), 4.40 (d, *J* = 12.8 Hz, 1H). $^{13}$**C NMR** (126 MHz, $C_6D_6$) δ 165.2, 156.2, 144.8, 143.4, 143.2, 140.3, 135.9, 135.3, 131.2, 130.1, 129.3, 129.2, 128.9, 128.6, 128.6, 127.7, 127.6, 127.1, 62.6, 57.1, 42.6. **HRMS** (ESI): exact mass calculated for $C_{25}H_{20}ClN_4O$ [$(M+H)^+$] 427.1320, found 427.1338.

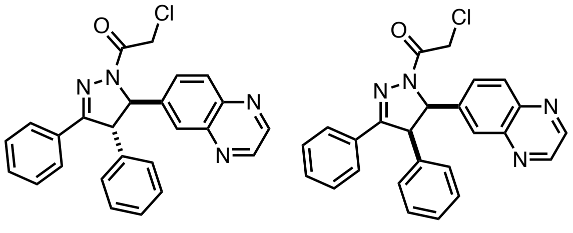

**HW-01-187A** **HW-01-187B**

**2-Chloro-1-(3,4-diphenyl-5-(quinoxalin-6-yl)-4,5-dihydro-1*H*-pyrazol-1-yl)ethan-1-one.** General Procedure C was followed starting from deoxybenzoin (118 mg, 0.599 mmol) and quinoxaline-6-carbaldehyde (32 mg, 0.20 mmol). Purification by column chromatography (EtOAc/hexane, 40:60) afforded the corresponding α,β-unsaturated ketone (57 mg) containing impurities. This product was used in the following transformation without further purification. $^{1}$**H NMR** (300 MHz, $CDCl_3$) δ 8.83 – 8.71 (m, 2H), 8.02 (d, *J* = 7.5 Hz, 2H), 7.96 – 7.82 (m, 2H), 7.74 – 7.29 (m, 10H).

General Procedure F was followed starting from the above product (57 mg). Purification by column chromatography (EtOAc/hexane, 45:55) afforded the *trans*-substituted chloroacetamide **HW-01-187A** (33 mg, 39% over three steps) as a white solid and the *cis*-substituted chloroacetamide **HW-01-187B** (6 mg, 7% over three steps) as a colorless film.

**HW-01-187A [trans]:** $^{1}$**H NMR** (600 MHz, $CDCl_3$): *δ* 8.84 (s, 2H), 8.15 (d, *J* = 8.7 Hz, 1H), 7.97 (d, *J* = 2.0 Hz, 1H), 7.68 (ddd, *J* = 8.8, 4.9, 1.6 Hz, 3H), 7.42–7.27 (m, 6H), 7.24–7.16 (m, 2H), 5.61 (d, *J* = 3.8 Hz, 1H), 4.75 (d, *J* = 13.4 Hz, 1H), 4.71 (d, *J* = 13.4 Hz, 1H), 4.64 (d, *J* = 3.8 Hz, 1H); $^{13}$**C NMR** (151 MHz, $CDCl_3$): *δ* 164.3, 156.9, 145.4, 145.2, 143.1, 142.7, 142.1, 138.7, 130.9, 130.7, 129.8, 129.8, 128.7, 128.3, 127.7, 127.6, 127.0, 125.8, 70.3, 61.5, 42.0; **HRMS** (ESI): exact mass calculated for $C_{25}H_{20}ClN_4O$ [$(M+MeCN+H)^+$] 468.1585, found 468.1574.

**HW-01-187B [cis]:** $^{1}$**H NMR** (600 MHz, $CDCl_3$): *δ* 8.75 (d, *J* = 1.8 Hz, 1H), 8.73 (d, *J* = 1.8 Hz, 1H), 7.74–7.70 (m, 2H), 7.64–7.60 (m, 2H), 7.37–7.33 (m, 1H), 7.31–7.27 (m, 2H), 7.20 (dd, *J* = 8.8, 2.0 Hz, 1H), 6.87 (s, 3H), 6.78 (s, 2H), 6.08 (d, *J* = 11.7 Hz, 1H), 5.34 (d, *J* = 11.7 Hz, 1H), 4.80 (d, *J* = 13.5 Hz, 1H), 4.67 (d, *J* = 13.5 Hz, 1H); $^{13}$**C NMR** (151 MHz, $CD_2Cl_2$) δ 165.4, 157.0, 145.6, 145.5, 142.8, 142.4, 138.7, 134.7, 130.8, 130.7, 130.0 (2C), 129.2 (2C), 128.9, 128.6, 128.1, 127.7, 66.8, 57.7, 43.0. **HRMS** (ESI): exact mass calculated for $C_{25}H_{19}ClN_4NaO$ [$(M+MeCN+Na)^+$] 490.1405, found 490.1400.

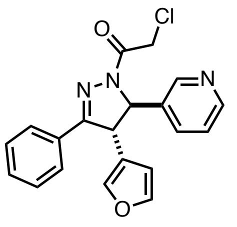
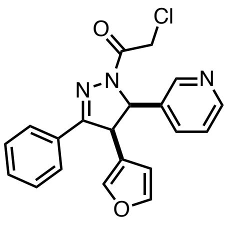

HW-01-198A HW-01-198B

**2-Chloro-1-(4-(furan-3-yl)-3-phenyl-5-(pyridin-3-yl)-4,5-dihydro-1*H*-pyrazol-1-yl)ethan-1-one.** General procedure E was followed starting from 1-phenyl-3-(pyridin-3-yl)prop-2-en-1-one (630 mg, 3.0 mmol, 1 equiv.). The intermediate product 2-bromo-1-phenyl-3-(pyridin-3-yl)prop-2-en-1-one was obtained as a light orange oil (896 mg, 99%).

A mixture of 2-bromo-1-phenyl-3-(pyridin-3-yl)prop-2-en-1-one (58 mg, 0.20 mmol, 1.0 equiv), 3-furanylboronic acid (29 mg, 0.26 mmol, 1.3 equiv), sodium carbonate (43 mg, 0.40 mmol, 2.0 equiv) and Pd(dppf)$Cl_2$ (7 mg, 10 µmol, 5 mol%) was taken up in toluene/$H_2O$ (4:1 v/v, 1 mL). The resulting suspension was sparged with nitrogen for 10 min. The flask was sealed and the reaction mixture was stirred at 100 °C for 1 h. More portions of 3-furanylboronic acid (29 mg, 0.26 mmol, 1.3 equiv), sodium carbonate (22 mg, 0.20 mmol, 1.0 equiv) and Pd(dppf)$Cl_2$ (7 mg, 10 µmol, 5 mol%) were then added, the mixture was sparged again with nitrogen for 10 min, the flask was sealed and stirring was continued at 100°C for 2 h. The reaction mixture was then allowed to cool to ambient temperature and partitioned between DCM (20 mL) and 1 M NaOH (10 mL). The phases were separated and the aqueous phase was extracted with DCM (2 x 20 mL). The combined organic phases were washed with NaCl solution (20 mL, sat. aqueous), dried over $Na_2SO_4$, filtered and concentrated under reduced pressure. Purification by column chromatography (EtOAc/hexane, 40:60) afforded the corresponding furyl-substituted α,β-unsaturated ketone (46 mg, d.r. = 2:1).

General Procedure F was followed starting from the above product (43 mg, 0.16 mmol). Purification by column chromatography (EtOAc/hexane, 50:50 to 75:25) afforded the *trans*-substituted chloroacetamide **HW-01-198A** (27 mg, 39% over three steps) as a yellowish oil. Further purification by pTLC (EtOAc/hexane, 80:20) afforded the *cis*-substituted chloroacetamide **HW-01-198B** (4 mg, 6% over three steps) as a colorless film.

**HW-01-198A [trans]: $^{1}$H NMR** (300 MHz, $CDCl_3$): *δ* 8.56 (dd, *J* = 5.0, 1.8 Hz, 2H), 7.73 (dd, *J* = 8.1, 1.7 Hz, 2H), 7.53 (dt, *J* = 7.9, 2.0 Hz, 1H), 7.44–7.27 (m, 6H), 6.26 (dd, *J* = 1.9, 0.9 Hz, 1H), 5.40 (d, *J* = 3.6 Hz, 1H), 4.63 (s, 2H), 4.54 (d, *J* = 3.6 Hz, 1H); **$^{13}$C NMR** (151 MHz, $CDCl_3$): *δ* 164.3, 156.3, 149.5, 147.3, 144.6, 139.5, 135.3, 133.3, 131.0, 129.6, 128.8, 127.4, 124.0, 122.9, 108.8, 67.3, 51.4, 41.8; **HRMS** (ESI): exact mass calculated for $C_{20}H_{17}ClN_3O_2$ [$(M+MeCN+H)^+$] 407.1269, found 407.1246.

**HW-01-198B [cis]: $^{1}$H NMR** (600 MHz, $CDCl_3$): *δ* 8.39 (dd, *J* = 4.9, 1.6 Hz, 1H), 8.32 (d, *J* = 2.3 Hz, 1H), 7.70–7.66 (m, 2H), 7.42–7.38 (m, 1H), 7.37–7.32 (m, 2H), 7.30 (d, *J* = 8.0 Hz, 1H), 7.13 (dd, *J* = 7.9, 4.8 Hz, 1H), 7.03 (d, *J* = 1.5 Hz, 2H), 5.78 (d, *J* = 11.4 Hz, 1H), 5.59 (s, 1H), 5.19 (d, *J* = 11.4 Hz, 1H), 4.73 (d, *J* = 13.5 Hz, 1H), 4.57 (d, *J* = 13.4 Hz, 1H); **$^{13}$C NMR** (151 MHz, $CDCl_3$): *δ* 165.0, 156.1, 148.0, 147.6, 143.7, 141.4, 135.1, 134.9, 130.8, 129.8, 128.7, 127.5, 123.2, 119.0, 110.5, 63.8, 47.9, 42.1; **HRMS** (ESI): exact mass calculated for $C_{20}H_{17}ClN_3O_2$ [$(M+MeCN+H)^+$] 407.1269, found 407.1242.

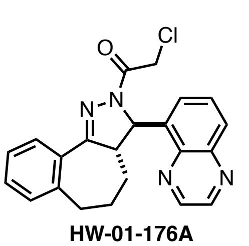
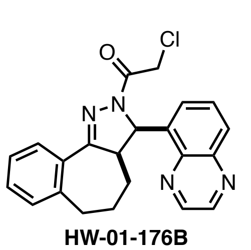

HW-01-176A HW-01-176B

**2-Chloro-1-(3-(quinoxalin-5-yl)-3a,4,5,6-tetrahydrobenzo[6,7]cyclohepta[1,2-c]pyrazol-2(3*H*)-yl)ethan-1-one.** General Procedure A was followed starting from 1-benzosuberone (30 µL, 0.20 mmol) and quinoxaline-5-carbaldehyde (63 mg, 0.40 mmol). The crude α,β-unsaturated ketone (42 mg) was used in the following transformation without further purification.

General Procedure F was followed starting from the above product (41 mg, 0.14 mmol). Purification by column chromatography (EtOAc/hexane, 40:60) afforded the *trans*-substituted chloroacetamide **HW-01-176A** (9 mg, 17% over three steps) as a colorless film and the *cis*-substituted chloroacetamide **HW-01-176B** (5 mg, 9% over three steps) as a colorless film.

**HW-01-176A [trans]: $^{1}$H NMR** $^{1}$H NMR (600 MHz, $CD_2Cl_2$) δ 8.86 (m, 2H), 8.03 (dd, *J* = 8.5, 1.4 Hz, 1H), 7.82 – 7.71 (m, 2H), 7.58 – 7.49 (m, 1H), 7.28 (m, 2H), 7.17 (dd, *J* = 7.4, 1.4 Hz, 1H), 6.23 (d, *J* = 4.4 Hz, 1H), 4.74 (d, *J* = 13.6 Hz, 1H), 4.56 (d, *J* = 13.6 Hz, 1H), 3.21 (dt, *J* = 12.3, 5.0 Hz, 1H), 2.95 – 2.80 (m, 2H), 2.74 (m, 1H), 2.12 – 1.93 (m, 2H), 1.74 (m, 1H). **$^{13}$C NMR** (151 MHz, $CD_2Cl_2$) δ 164.3, 162.5, 145.7, 144.5,

143.9, 142.7, 140.8, 138.8, 132.2, 130.7, 130.5, 130.1, 129.5, 129.0, 126.8, 125.9, 64.5, 57.9, 42.8, 36.7, 34.2, 26.2. **HRMS** (ESI): exact mass calculated for $C_{22}H_{20}ClN_4O$ [$(M+H)^+$] 391.1320, found 391.1312.

**HW-01-176B [cis]:** $^1$**H NMR** (600 MHz, $CDCl_3$): *δ* 8.88 (d, *J* = 6.7 Hz, 2H), 8.03 (td, *J* = 7.6, 6.9, 1.5 Hz, 2H), 7.71 (t, *J* = 7.8 Hz, 1H), 7.46 (d, *J* = 7.2 Hz, 1H), 7.33 (dtd, *J* = 18.5, 7.5, 1.6 Hz, 2H), 7.16–7.11 (m, 1H), 7.02 (d, *J* = 11.3 Hz, 1H), 4.73 (d, *J* = 13.0 Hz, 1H), 4.57 (d, *J* = 13.0 Hz, 1H), 4.03 (ddd, *J* = 13.0, 11.3, 3.5 Hz, 1H), 3.00 (ddd, *J* = 14.9, 10.6, 4.4 Hz, 1H), 2.65 (ddd, *J* = 14.7, 6.2, 3.9 Hz, 1H), 1.70 (ddp, *J* = 21.4, 10.7, 3.6 Hz, 2H), 1.32 (ddd, *J* = 13.5, 6.4, 3.3 Hz, 1H), 0.97 (ddt, *J* = 13.4, 9.6, 6.7 Hz, 1H); $^{13}$**C NMR** (151 MHz, $CDCl_3$): *δ* 164.0, 162.0, 145.0, 144.2, 143.0, 141.4, 140.8, 135.0, 130.9, 130.8, 130.5, 129.8, 129.3, 128.9, 126.8, 126.7, 59.4, 50.0, 42.1, 34.2, 25.8, 24.7; **HRMS** (ESI): exact mass calculated for $C_{22}H_{20}ClN_4O$ [$(M+H)^+$] 391.1320, found 391.1295.

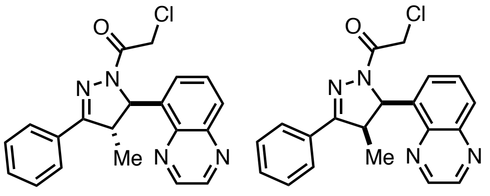

**HW-02-01A** **HW-02-01B**

**2-Chloro-1-(4-methyl-3-phenyl-5-(quinoxalin-5-yl)-4,5-dihydro-1*H*-pyrazol-1-yl)ethan-1-one.** General Procedure D was followed starting from propiophenone (50 µL, 0.37 mmol) and quinoxaline-5-carbaldehyde (59 mg, 0.37 mmol). Purification by column chromatography (EtOAc/hexane, 25:75) afforded the corresponding α,β-unsaturated ketone (49 mg).
General Procedure F was followed starting from the above product (46 mg, 0.17 mmol). Purification by column chromatography (EtOAc/hexane, 35:65 to 40:60) afforded the *trans*-substituted chloroacetamide **HW-02-01A** (13 mg, 10% over three steps) as a white solid. Further purification by pTLC (EtOAc/hexane, 70:30) afforded the *cis*-substituted chloroacetamide **HW-02-01B** (2.5 mg, 2% over three steps, containing 9% of the *trans* product **xx**) as a white solid.
**HW-02-01A [trans]:** $^1$**H NMR** (600 MHz, $CDCl_3$): *δ* 8.90 (d, *J* = 1.8 Hz, 1H), 8.88 (d, *J* = 1.8 Hz, 1H), 8.04 (dd, *J* = 8.4, 1.3 Hz, 1H), 7.75–7.71 (m, 2H), 7.69 (dd, *J* = 8.4, 7.2 Hz, 1H), 7.49–7.35 (m, 4H), 6.38 (d, *J* = 2.9 Hz, 1H), 4.78 (d, *J* = 13.2 Hz, 1H), 4.63 (d, *J* = 13.1 Hz, 1H), 3.55 (qd, *J* = 7.2, 3.0 Hz, 1H), 1.64 (d, *J* = 7.2 Hz, 3H); $^{13}$**C NMR** (151 MHz, $CDCl_3$): *δ* 164.3, 160.8, 144.9, 143.9, 143.4, 140.6, 137.5, 130.6, 130.0, 129.1, 128.8, 127.2, 125.1, 64.5, 50.0, 41.9, 18.9; **HRMS** (ESI): exact mass calculated for $C_{20}H_{18}ClN_4O$ [$(M+H)^+$] 365.1164, found 365.1144.
**HW-02-01B [cis]:** $^1$**H NMR** (500 MHz, $CDCl_3$): *δ* 8.91 (d, *J* = 1.8 Hz, 1H), 8.88 (d, *J* = 1.8 Hz, 1H), 8.07 (dd, *J* = 8.5, 1.4 Hz, 1H), 7.80–7.75 (m, 3H), 7.55 (dt, *J* = 7.2, 0.9 Hz, 1H), 7.45 (dd, *J* = 5.0, 1.9 Hz, 3H), 6.81 (d, *J* = 11.3 Hz, 1H), 4.81 (d, *J* = 13.4 Hz, 1H), 4.54 (d, *J* = 13.4 Hz, 1H), 4.38 (dq, *J* = 11.2, 7.5 Hz, 1H), 0.64 (d, *J* = 7.5 Hz, 3H); **HRMS** (ESI): exact mass calculated for $C_{20}H_{18}ClN_4O$ [$(M+H)^+$] 365.1164, found 365.1140.

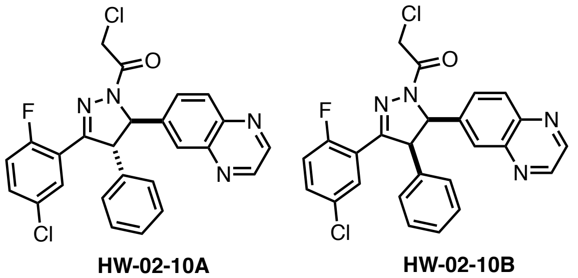

**HW-02-10A** **HW-02-10B**

**2-Chloro-1-(3-(5-chloro-2-fluorophenyl)-4-phenyl-5-(quinoxalin-6-yl)-4,5-dihydro-1*H*-pyrazol-1-yl)ethan-1-one.** General procedure E was followed starting from 1-(5-chloro-2-fluorophenyl)-3-(quinoxalin-6-yl)prop-2-en-1-one (2.15g, 6.9 mmol, 1 equiv.). The intermediate product 2-bromo-1-(5-chloro-2-fluorophenyl)-3-(quinoxalin-6-yl)prop-2-en-1-one was obtained as a beige solid (2.66g, 99%). $^1$**H NMR** (300 MHz, $CDCl_3$) δ 8.90 (s, 2H), 8.63 (s, 1H), 8.18 (m, 2H), 7.96 (s, 1H), 7.64 – 7.45 (m, 2H), 7.17 (t, *J* = 8.8, 1H).

The vinyl bromide from above (2.66 g, 6.8 mmol, 1.0 equiv), phenylboronic acid (1.08 g, 8.8 mmol, 1.3 equiv), sodium carbonate (1.4 g, 0.14 mmol, 2.0 equiv) and Pd(dppf)$Cl_2$ (277 mg, 0.34 mmol, 5 mol%) was taken up in toluene/$H_2O$ (4:1 v/v, 34 mL). The resulting suspension was sparged with nitrogen for 10 min. The flask was sealed and the reaction mixture was stirred at 100 °C for 1.5 h. The reaction mixture was then allowed to cool to ambient temperature and partitioned between EtOAc (100 mL) and 1 M NaOH (30 mL). The phases were separated and the aqueous phase was extracted with EtOAc (3 x 70 mL). The combined organic phases were washed with NaCl solution (50 mL, sat. aqueous), dried over $Na_2SO_4$, filtered and concentrated under reduced pressure. Purification by column chromatography (EtOAc/hexane, 4:1 to 1:1 Hexane/EtOAc) afforded the corresponding phenyl-substituted α,β-unsaturated ketone (2.0 g, 76%) as a light yellow oil. **$^1$H NMR** (500 MHz, $CDCl_3$) *major isomer*, δ 8.82 – 8.77 (m, 2H), 7.90 (d, *J* = 1.9 Hz, 1H), 7.83 (d, *J* = 8.8 Hz, 1H), 7.59 (dd, *J* = 5.8, 2.7 Hz, 1H), 7.52 (s, 1H), 7.50 – 7.43 (m, 2H), 7.39 (dt, *J* = 4.9, 2.6 Hz, 3H), 7.34 (dd, *J* = 8.9, 2.1 Hz, 1H), 7.27 (d, *J* = 2.1 Hz, 1H), 7.07 (t, *J* = 8.9 Hz, 1H).

General Procedure F was followed starting from the corresponding chalcone (200 mg, 0.51 mmol). Purification by column chromatography (EtOAc/hexane, 50:50) afforded the *trans*-substituted chloroacetamide **HW-02-10A** (103 mg, 42% over two steps) as a yellow oil. Further purification by pTLC (EtOAc/hexane, 70:30) and column chromatography (MeOH/DCM, 1:99) afforded the *cis*-substituted chloroacetamide **HW-01-10B** (22 mg, 9% over three steps) as a colorless film.

**HW-02-10A [trans]: $^1$H NMR** (600 MHz, $CDCl_3$): *δ* 8.86 (s, 2H), 8.17 (d, *J* = 8.7 Hz, 1H), 7.96 (d, *J* = 2.0 Hz, 1H), 7.93 (dd, *J* = 6.2, 2.7 Hz, 1H), 7.67 (dd, *J* = 8.7, 2.1 Hz, 1H), 7.38–7.28 (m, 4H), 7.15–7.11 (m, 2H), 6.93 (dd, *J* = 10.6, 8.8 Hz, 1H), 5.63 (d, *J* = 4.0 Hz, 1H), 4.76 (dd, *J* = 4.0, 2.9 Hz, 1H), 4.74 (d, *J* = 13.4 Hz, 1H), 4.66 (d, *J* = 13.4 Hz, 1H); **$^{13}$C NMR** (151 MHz, $CDCl_3$): *δ* 164.5, 159.1 (d, *J* = 254.5 Hz), 153.1 (d, *J* = 3.9 Hz), 145.5, 145. 2, 143.1, 142.7, 141.9, 138.0, 132.2 (d, *J* = 8.9 Hz), 130.9, 129.9 (d, *J* = 3.3 Hz), 129.6, 129.1 (d, *J* = 3.2 Hz), 128.4, 127.7, 127.0, 125.8, 119.5 (d, *J* = 12.9 Hz), 118.1 (d, *J* = 24.3 Hz), 70.2, 62.6 (d, *J* = 6.1 Hz), 41.9; **HRMS** (ESI): exact mass calculated for $C_{25}H_{18}Cl_2FN_4O$ [$(M+MeCN+H)^+$] 520.1101, found 520.1101.

**HW-01-10B [cis]: $^1$H NMR** (600 MHz, $CDCl_3$): *δ* 8.75 (d, *J* = 1.9 Hz, 1H), 8.73 (d, *J* = 1.8 Hz, 1H), 7.89 (dd, *J* = 6.1, 2.7 Hz, 1H), 7.73 (d, *J* = 8.7 Hz, 1H), 7.69 (d, *J* = 2.0 Hz, 1H), 7.29 (ddd, *J* = 8.8, 4.2, 2.7 Hz, 1H), 7.17 (dd, *J* = 8.7, 2.0 Hz, 1H), 6.90–6.82 (m, 4H), 6.76–6.65 (m, 2H), 6.09 (d, *J* = 12.0 Hz, 1H), 5.46 (dd, *J* = 12.0, 2.6 Hz, 1H), 4.78 (d, *J* = 13.5 Hz, 1H), 4.62 (d, *J* = 13.5 Hz, 1H); **$^{19}$F NMR** (376 MHz, $CDCl_3$): *δ* –112.4; **$^{13}$C NMR** (151 MHz, $CDCl_3$): *δ* 165.2, 158.9 (d, *J* = 254.3 Hz), 153.5 (d, *J* = 3.4 Hz), 144.9, 144.7, 142.1, 141.9, 137.9, 133.2, 132.0 (d, *J* = 8.8 Hz), 129.9 (d, *J* = 3.0 Hz), 129.2, 129.1 (d, *J* = 3.3 Hz), 128.9, 128.2, 127.5, 126.9, 120.1 (d, *J* = 13.5 Hz), 118.0 (d, *J* = 23.9 Hz), 66.2, 58.6 (d, *J* = 5.3 Hz), 42.1; **HRMS** (ESI): exact mass calculated for $C_{25}H_{18}Cl_2FN_4O$ [$(M+MeCN+H)^+$] 520.1101, found 520.1088.

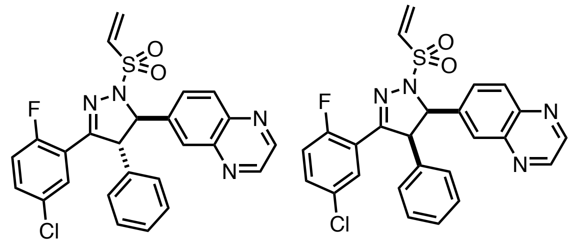


**PM-02-71A** **PM-02-71B**

**6-(3-(5-chloro-2-fluorophenyl)-4-phenyl-1-(vinylsulfonyl)-4,5-dihydro-1*H*-pyrazol-5-yl)quinoxaline.** General procedure E was followed starting from 1-(5-chloro-2-fluorophenyl)-3-(quinoxalin-6-yl)prop-2-en-1-one (2.15g, 6.9 mmol, 1 equiv.). The intermediate product 2-bromo-1-(5-chloro-2-fluorophenyl)-3-(quinoxalin-6-yl)prop-2-en-1-one was obtained as a beige solid (2.66g, 99%). **$^1$H NMR** (300 MHz, $CDCl_3$) δ 8.90 (s, 2H), 8.63 (s, 1H), 8.18 (m, 2H), 7.96 (s, 1H), 7.64 – 7.45 (m, 2H), 7.17 (t, *J* = 8.8, 1H).

The vinyl bromide from above (2.66 g, 6.8 mmol, 1.0 equiv), phenylboronic acid (1.08 g, 8.8 mmol, 1.3 equiv), sodium carbonate (1.4 g, 0.14 mmol, 2.0 equiv) and Pd(dppf)$Cl_2$ (277 mg, 0.34 mmol, 5 mol%) was taken up in toluene/$H_2O$ (4:1 v/v, 34 mL). The resulting suspension was sparged with nitrogen for 10 min. The flask was sealed and the reaction mixture was stirred at 100 °C for 1.5 h. The reaction mixture was then allowed to cool to ambient temperature and partitioned between EtOAc (100 mL) and 1 M NaOH (30 mL). The phases were separated and the aqueous phase was extracted with EtOAc (3 x 70 mL). The combined organic phases were washed with NaCl solution (50 mL, sat. aqueous), dried over $Na_2SO_4$, filtered and concentrated under reduced pressure. Purification by column chromatography (EtOAc/hexane, 4:1 to 1:1 Hexane/EtOAc) afforded the corresponding phenyl-substituted α,β-unsaturated ketone (2.0 g, 76%) as a light yellow oil. **$^1$H NMR** (500 MHz, $CDCl_3$) *major isomer*, δ 8.82 – 8.77 (m, 2H), 7.90 (d, *J* = 1.9 Hz, 1H), 7.83 (d, *J* = 8.8 Hz, 1H), 7.59 (dd, *J* = 5.8, 2.7 Hz, 1H), 7.52 (s, 1H), 7.50 – 7.43 (m, 2H), 7.39 (dt, *J* = 4.9, 2.6 Hz, 3H), 7.34 (dd, *J* = 8.9, 2.1 Hz, 1H), 7.27 (d, *J* = 2.1 Hz, 1H), 7.07 (t, *J* = 8.9 Hz, 1H).

General Procedure H was followed starting from the above product (267 mg, 0.69 mmol, 1.0 equiv.). Purification by column chromatography (EtOAc:hexane, 50:50 to 100:0) afforded the *trans*-substituted vinyl sulfonamide **PM-02-71A** (73 mg, 21%) as a white solid and the *cis*-substituted vinylsulfonamide **PM-02-71B** (68 mg, 20%) as a white solid.

**PM-02-71A [trans] : $^1$H NMR** (600 MHz, $CD_2Cl_2$) δ 8.87 – 8.83 (m, 2H), 8.17 (d, *J* = 8.7 Hz, 1H), 7.94 (d, *J* = 2.1 Hz, 1H), 7.87 (dd, *J* = 6.1, 2.7 Hz, 1H), 7.80 (dd, *J* = 8.7, 2.1 Hz, 1H), 7.31 (m, 4H), 7.05 – 7.03 (m, 2H), 6.91 (dd, *J* = 10.4, 8.8 Hz, 1H), 6.77 (dd, *J* = 16.6, 10.0 Hz, 1H), 6.40 (d, *J* = 16.7 Hz, 1H), 6.25 (d, *J* = 10.0 Hz, 1H), 5.13 (d, *J* = 8.6 Hz, 1H), 4.83 (dd, *J* = 8.6, 3.1 Hz, 1H). **$^{13}$C NMR** (151 MHz, $CD_2Cl_2$) δ 159.1 (d, *J* =

253.5 Hz), 154.5 (d, *J* = 3.6 Hz), 146.2, 146.0, 143.3, 142.3, 138.1, 132.8, 132.5 (d, *J* = 8.8 Hz), 131.7, 131.0, 130.2 (d, *J* = 3.1 Hz), 129.9 (d, *J* = 3.4 Hz), 129.8, 128.7, 128.5, 128.3, 127.8, 120.2 (d, *J* = 13.8 Hz), 118.3 (d, *J* = 24.2 Hz), 75.2, 64.6 (d, *J* = 5.0 Hz). **HRMS** (ESI): exact mass calculated for $C_{25}H_{18}ClFN_4O_2S$ [(M+MeCN+H)$^+$] 534.1161, found 534.1164.

**PM-02-71B [cis] : $^1$H NMR** (600 MHz, $CD_2Cl_2$) δ 8.77 – 8.71 (m, 2H), 8.06 (dd, *J* = 6.3, 2.7 Hz, 1H), 7.88 (d, *J* = 2.0 Hz, 1H), 7.69 (d, *J* = 8.7 Hz, 1H), 7.40 (dd, *J* = 8.7, 2.0 Hz, 1H), 7.35 (ddd, *J* = 8.8, 4.3, 2.7 Hz, 1H), 7.01 – 6.93 (m, 4H), 6.86 – 6.78 (m, 2H), 6.74 (dd, *J* = 16.7, 10.0 Hz, 1H), 6.40 (d, *J* = 16.7 Hz, 1H), 6.30 (d, *J* = 9.9 Hz, 1H), 5.46 (d, *J* = 10.8 Hz, 1H), 5.18 (dd, *J* = 10.8, 2.5 Hz, 1H). **$^{13}$C NMR** (151 MHz, $CD_2Cl_2$) δ 159.6 (d, *J* = 252.2 Hz), 156.0 (d, *J* = 3.5 Hz), 145.7 (d, *J* = 8.5 Hz), 142.7, 142.6, 138.1, 133.0, 132.8, 132.8, 132.5, 131.7, 130.3, 129.9, 129.7 (d, *J* = 3.3 Hz), 129.5, 128.9, 128.8, 128.7, 128.3, 120.1, 118.5 (d, *J* = 24.3 Hz), 71.0, 60.4 (d, *J* = 7.0 Hz). **HRMS** (ESI): exact mass calculated for $C_{25}H_{18}ClFN_4O_2S$ [(M+MeCN+H)$^+$] 534.1161, found 534.1159.

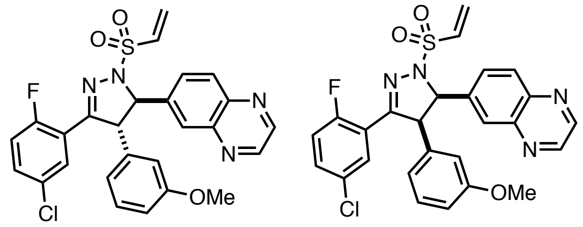

**PM-02-167A** **PM-02-167B**

**6-(3-(5-chloro-2-fluorophenyl)-4-(3-methoxyphenyl)-1-(vinylsulfonyl)-4,5-dihydro-1*H*-pyrazol-5-yl)quinoxaline.** General procedure E was followed starting from 1-(5-chloro-2-fluorophenyl)-3-(quinoxalin-6-yl)prop-2-en-1-one (2.15g, 6.9 mmol, 1 equiv.). The intermediate product 2-bromo-1-(5-chloro-2-fluorophenyl)-3-(quinoxalin-6-yl)prop-2-en-1-one was obtained as a beige solid (2.66g, 99%).**$^1$H NMR** (300 MHz, $CDCl_3$) δ 8.90 (s, 2H), 8.63 (s, 1H), 8.18 (m, 2H), 7.96 (s, 1H), 7.64 – 7.45 (m, 2H), 7.17 (t, *J* = 8.8, 1H).

The vinyl bromide from above (200 mg, 0.51 mmol, 1.0 equiv), 3-methoxyphenylboronic acid (58 mg, 0.61 mmol, 1.2 equiv), sodium carbonate (162 mg, 1.5 mmol, 2.0 equiv) and Pd(dppf)$Cl_2$ (19 mg, 0.026 mmol, 5 mol%) was taken up in toluene/$H_2O$ (4:1 v/v, 2.5 mL). The resulting suspension was sparged with nitrogen for 10 min. The flask was sealed and the reaction mixture was stirred at 100 °C for 2 h. The reaction mixture was then allowed to cool to ambient temperature and partitioned between DCM (20 mL) and 1 M NaOH (10 mL). The phases were separated and the aqueous phase was extracted with DCM (2 x 20 mL). The combined organic phases were washed with NaCl solution (20 mL, sat. aqueous), dried over $Na_2SO_4$, filtered and concentrated under reduced pressure. Purification by column chromatography (EtOAc/hexane, 10:90 to 60:40) afforded the corresponding phenyl-substituted α,β-unsaturated ketone (174 mg, 81%). **$^1$H NMR** (300 MHz, $CDCl_3$) *major isomer* δ 8.75 (s, 2H), 7.89 (t, *J* = 2.0 Hz, 1H), 7.80 (d, *J* = 8.8 Hz, 1H), 7.54 (dd, *J* = 5.8, 2.7 Hz, 1H), 7.46 (s, 1H), 7.41 (ddd, *J* = 8.8, 4.4, 2.7 Hz, 1H), 7.33 (dd, *J* = 8.9, 2.0 Hz, 1H), 7.30 – 7.20 (m, 1H), 7.04 (t, *J* = 8.9 Hz, 1H), 6.92 – 6.87 (m, 1H), 6.84 – 6.73 (m, 2H), 3.70 (s, 3H).

General Procedure H was followed starting from the above product (35 mg, 0.08 mmol, 1.0 equiv.). Purification by column chromatography (EtOAc:hexane, 30:70 to 100:0) afforded the *trans*-substituted vinyl sulfonamide **PM-02-167A** (22 mg, 54%) as a white solid and the *cis*-substituted vinylsulfonamide **PM-02-167B** (6 mg, 14%) as a white solid.

**PM-02-167A [trans] : $^1$H NMR** (600 MHz, $CD_2Cl_2$) δ 8.86 (q, *J* = 1.9 Hz, 2H), 8.18 (d, *J* = 8.7 Hz, 1H), 7.97 (d, *J* = 2.0 Hz, 1H), 7.86 (dd, *J* = 6.2, 2.7 Hz, 1H), 7.81 (dd, *J* = 8.7, 2.1 Hz, 1H), 7.32 (ddd, *J* = 8.8, 4.3, 2.7 Hz, 1H), 7.24 (t, *J* = 7.9 Hz, 1H), 6.93 (dd, *J* = 10.4, 8.8 Hz, 1H), 6.83 (ddd, *J* = 8.3, 2.6, 0.9 Hz, 1H), 6.78 (dd, *J* = 16.7, 10.0 Hz, 1H), 6.63 (dt, *J* = 7.6, 1.2 Hz, 1H), 6.57 (t, *J* = 2.2 Hz, 1H), 6.41 (d, *J* = 16.6 Hz, 1H), 6.25 (d, *J* = 9.9 Hz, 1H), 5.17 (d, *J* = 8.5 Hz, 1H), 4.80 (dd, *J* = 8.5, 3.0 Hz, 1H), 3.71 (s, 3H). **$^{13}$C NMR** (151 MHz, $CD_2Cl_2$) δ 160.7, 159.2 (d, *J* = 253.4 Hz), 154.4 (d, *J* = 4.0 Hz), 146.1, 145.9, 143.3 (d, *J* = 5.4 Hz), 142.4, 139.6, 132.9, 132.5 (d, *J* = 9.0 Hz), 131.6, 131.0, 130.8, 130.2 (d, *J* = 3.6 Hz), 129.8 (d, *J* = 4.0 Hz), 128.6, 127.8, 120.4, 120.3 (d, *J* = 13.9 Hz), 118.3 (d, *J* = 24.0 Hz), 114.4, 113.6, 75.0, 64.5 (d, *J* = 5.6 Hz), 55.6. **HRMS** (ESI): exact mass calculated for $C_{26}H_{21}ClFN_4O_3S$ [(M+MeCN+H)$^+$] 564.1267, found 564.1267.

**PM-02-167B [cis] : $^1$H NMR** (400 MHz, CD2Cl2) δ 8.76 (d, *J* = 5.9 Hz, 2H), 8.04 (dd, *J* = 6.3, 2.7 Hz, 1H), 7.90 (s, 1H), 7.73 (d, *J* = 8.8 Hz, 1H), 7.45 (dd, *J* = 8.7, 2.0 Hz, 1H), 7.35 (dt, *J* = 7.3, 3.1 Hz, 1H), 7.03 – 6.94 (m, 1H), 6.90 (t, *J* = 8.0 Hz, 1H), 6.73 (dd, *J* = 16.6, 10.0 Hz, 1H), 6.54 – 6.23 (m, 5H), 5.45 (d, *J* = 10.8 Hz, 1H), 5.15 (dd, *J* = 10.9, 2.5 Hz, 1H), 3.48 (s, 3H). **$^{13}$C NMR** (151 MHz, $CD_2Cl_2$) δ 160.1, 159.6 (d, *J* = 254.2 Hz), 155.9 (d, *J* = 3.8 Hz), 145.7, 145.6, 142.7, 142.6, 138.1, 134.5, 132.8 (d, *J* = 8.9 Hz), 132.4, 131.7, 130.3 (d, *J* = 3.2 Hz), 129.9, 129.7 (d, *J* = 3.2 Hz), 128.8, 128.66, 121.8, 120.2 (d, *J* = 12.8 Hz), 118.5 (d, *J* = 24.3

Hz), 115.4, 113.6, 70.9, 60.4 (d, *J* = 6.9 Hz), 55.5. **HRMS** (ESI): exact mass calculated for $C_{26}H_{20}ClFN_4NaO_3S$ [(M+MeCN+Na)$^+$] 586.1087, found 586.1109.

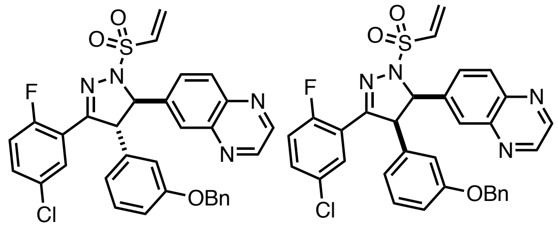

PM-2-162A PM-2-162B

**6-(4-(3-(benzyloxy)phenyl)-3-(5-chloro-2-fluorophenyl)-1-(vinylsulfonyl)-4,5-dihydro-1*H*-pyrazol-5-yl)quinoxaline.** General procedure E was followed starting from 1-(5-chloro-2-fluorophenyl)-3-(quinoxalin-6-yl)prop-2-en-1-one (2.15g, 6.9 mmol, 1 equiv.). The intermediate product 2-bromo-1-(5-chloro-2-fluorophenyl)-3-(quinoxalin-6-yl)prop-2-en-1-one was obtained as a beige solid (2.66g, 99%). **$^1$H NMR** (300 MHz, $CDCl_3$) δ 8.90 (s, 2H), 8.63 (s, 1H), 8.18 (m, 2H), 7.96 (s, 1H), 7.64 – 7.45 (m, 2H), 7.17 (t, *J* = 8.8, 1H).

The vinyl bromide from above (200 mg, 0.51 mmol, 1.0 equiv), 3-benzyloxyphenylboronic acid (139 mg, 0.61 mmol, 1.2 equiv), sodium carbonate (162 mg, 1.5 mmol, 2.0 equiv) and Pd(dppf)$Cl_2$ (19 mg, 0.026 mmol, 5 mol%) was taken up in toluene/$H_2O$ (4:1 v/v, 2.5 mL). The resulting suspension was sparged with nitrogen for 10 min. The flask was sealed and the reaction mixture was stirred at 100 °C for 2 h. The reaction mixture was then allowed to cool to ambient temperature and partitioned between DCM (20 mL) and 1 M NaOH (10 mL). The phases were separated and the aqueous phase was extracted with DCM (2 x 20 mL). The combined organic phases were washed with NaCl solution (20 mL, sat. aqueous), dried over $Na_2SO_4$, filtered and concentrated under reduced pressure. Purification by column chromatography (EtOAc/hexane, 10:90 to 60:40) afforded the corresponding phenyl-substituted α,β-unsaturated ketone (203 mg, 80%). **$^1$H NMR** (300 MHz, $CDCl_3$) δ 8.81 (s, 2H), 7.92 (d, *J* = 1.9 Hz, 1H), 7.82 (d, *J* = 8.8 Hz, 1H), 7.58 (dd, *J* = 5.8, 2.7 Hz, 1H), 7.49 (s, 1H), 7.40 – 7.27 (m, 8H), 7.09 (t, *J* = 8.9 Hz, 2H), 6.92 – 6.83 (m, 2H), 5.00 (s, 2H).

General Procedure H was followed starting from the above product (41 mg, 0.08 mmol, 1.0 equiv.). Purification by column chromatography (EtOAc:hexane, 20:80 to 100:0) afforded the *trans*-substituted vinyl sulfonamide **PM-02-162A** (28 mg, 59%) as a white solid and the *cis*-substituted vinylsulfonamide **PM-02-162B** (9 mg, 20%) as a white solid.

**PM-02-162A [trans] : $^1$H NMR** (600 MHz, CD2Cl2) δ 8.85 (m, 2H), 8.17 (d, *J* = 8.7 Hz, 1H), 7.96 (d, *J* = 2.0 Hz, 1H), 7.85 (dd, *J* = 6.1, 2.7 Hz, 1H), 7.79 (dd, *J* = 8.7, 2.1 Hz, 1H), 7.38 – 7.35 (m, 4H), 7.34 – 7.29 (m, 2H), 7.27 – 7.21 (m, 1H), 6.95 – 6.89 (m, 2H), 6.75 (dd, *J* = 16.6, 9.9 Hz, 1H), 6.66 – 6.63 (m, 2H), 6.40 (d, *J* = 16.6 Hz, 1H), 6.22 (d, *J* = 9.9 Hz, 1H), 5.16 (d, *J* = 8.5 Hz, 1H), 4.98 (d, *J* = 1.6 Hz, 2H), 4.80 (dd, *J* = 8.5, 3.0 Hz, 1H). **$^{13}$C NMR** (151 MHz, $CD_2Cl_2$) δ 159.8, 159.2 (d, *J* = 253.8 Hz), 154.3 (d, *J* = 3.9 Hz), 146.1, 145.9, 143.3 (d, *J* = 1.9 Hz), 142.3, 139.7, 137.1, 132.9, 132.5 (d, *J* = 9.4 Hz), 131.6, 131.0, 130.9, 130.2 (d, *J* = 3.8 Hz), 129.8 (d, *J* = 3.8 Hz), 128.9, 128.5, 128.5, 127.9, 127.8, 120.7, 120.2 (d, *J* = 14.0 Hz), 118.3 (d, *J* = 24.0 Hz), 115.1, 114.8, 75.0, 70.5, 64.5 (d, *J* = 5.5 Hz). **HRMS** (ESI): exact mass calculated for $C_{32}H_{24}ClFN_4NaO_3S$ [(M+Na)$^+$] 621.1134, found 621.1110.

**PM-02-162B [cis] : $^1$H NMR** (600 MHz, $CD_2Cl_2$) δ 8.78 – 8.73 (m, 2H), 8.02 (dd, *J* = 6.3, 2.7 Hz, 1H), 7.90 (d, *J* = 2.0 Hz, 1H), 7.69 (d, *J* = 8.8 Hz, 1H), 7.38 – 7.27 (m, 6H), 7.23 (d, *J* = 7.4 Hz, 2H), 6.99 (dd, *J* = 10.7, 8.8 Hz, 1H), 6.73 (dd, *J* = 16.7, 10.0 Hz, 1H), 6.57 (dd, *J* = 8.4, 2.5 Hz, 1H), 6.45 – 6.37 (m, 3H), 6.29 (d, *J* = 10.0 Hz, 1H), 5.45 (d, *J* = 10.8 Hz, 1H), 5.13 (dd, *J* = 10.9, 2.4 Hz, 1H), 4.85 – 4.62 (m, 2H). **$^{13}$C NMR** (151 MHz, $CD_2Cl_2$) δ 159.6 (d, *J* = 254.3 Hz), 159.2, 155.8 (d, *J* = 4.4 Hz), 145.7, 145.6, 142.7, 142.6, 138.1, 137.2, 134.5, 132.8 (d, *J* = 9.3 Hz), 132.4, 131.7, 130.3 (d, *J* = 3.5 Hz), 129.9, 129.8, 129.6 (d, *J* = 4.0 Hz), 128.9, 128.8, 128.6 (d, *J* = 3.3 Hz), 128.3, 127.7, 122.1 (d, *J* = 5.6 Hz), 120.1 (d, *J* = 13.1 Hz), 118.5 (d, *J* = 24.5 Hz), 114.8, 71.0, 70.3, 60.4 (d, *J* = 7.1 Hz). **HRMS** (ESI): exact mass calculated for $C_{32}H_{24}ClFN_4NaO_3S$ [(M+Na)$^+$] 621.1134, found 621.1121.

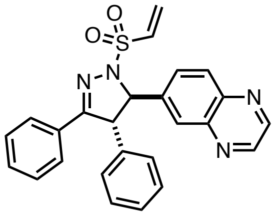

**PM-02-103A**

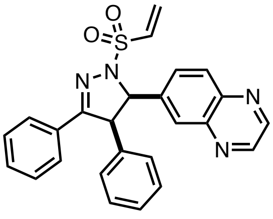

**PM-02-103B**

**6-(3,4-diphenyl-1-(vinylsulfonyl)-4,5-dihydro-1*H*-pyrazol-5-yl)quinoxaline.** General Procedure C was followed starting from deoxybenzoin (2.9 g, 15 mmol) and quinoxaline-6-carbaldehyde (790 mg, 5.0 mmol). Purification by column chromatography (EtOAc/hexane, 1:4 to 1:0) afforded the corresponding α,β-unsaturated ketone (1.5 g, 89%). **$^1$H NMR** (300 MHz, $CDCl_3$) δ 8.83 – 8.71 (m, 2H), 8.02 (d, *J* = 7.5 Hz, 2H), 7.96 – 7.82 (m, 2H), 7.74 – 7.29 (m, 10H).

General Procedure H was followed starting from the above product (108 mg, 0.32 mmol, 1.0 equiv.). Purification by column chromatography (EtOAc:hexane, 40:60 to 100:0) afforded the *trans*-substituted vinyl sulfonamide **PM-02-103A** (70 mg, 49% over three steps) as a white solid and the *cis*-substituted vinylsulfonamide **PM-02-103B** (19 mg, 13% over three steps) as a white solid.

**PM-02-103A [trans] : $^1$H NMR** (600 MHz, $CD_2Cl_2$) δ 8.85 (d, *J* = 1.7 Hz, 2H), 8.16 (d, *J* = 8.7 Hz, 1H), 7.95 (d, *J* = 2.0 Hz, 1H), 7.80 (dd, *J* = 8.7, 2.0 Hz, 1H), 7.64 – 7.58 (m, 2H), 7.38 – 7.32 (m, 4H), 7.32 – 7.25 (m, 2H), 7.13 (dd, *J* = 8.0, 1.6 Hz, 2H), 6.76 (dd, *J* = 16.6, 10.0 Hz, 1H), 6.37 (d, *J* = 16.7 Hz, 1H), 6.20 (d, *J* = 9.9 Hz, 1H), 5.13 (d, *J* = 8.0 Hz, 1H), 4.76 (d, *J* = 8.0 Hz, 1H). **$^{13}$C NMR** (151 MHz, $CD_2Cl_2$) δ 157.9, 146.1, 145.9, 143.3, 143.3, 142.7, 139.2, 133.0, 131.1, 131.0, 130.9, 130.3, 130.0, 128.9, 128.6, 128.6, 128.3, 128.1, 127.7, 74.9, 63.6. **HRMS** (ESI): exact mass calculated for $C_{25}H_{21}N_4O_2S$ [$(M+H)^+$] 441.1380, found 441.1390.

**PM-02-103B [cis] : 1H** NMR (600 MHz, $CD_2Cl_2$) δ 8.75 (d, *J* = 9.1 Hz, 2H), 7.90 (d, *J* = 1.9 Hz, 1H), 7.77 – 7.67 (m, 3H), 7.44 (dd, *J* = 8.7, 2.0 Hz, 1H), 7.42 – 7.38 (m, 1H), 7.35 (t, *J* = 7.5 Hz, 2H), 7.05 – 6.94 (m, 3H), 6.89 (s, 2H), 6.77 (dd, *J* = 16.7, 10.0 Hz, 1H), 6.38 (d, *J* = 16.7 Hz, 1H), 6.27 (d, *J* = 10.0 Hz, 1H), 5.43 (d, *J* = 10.5 Hz, 1H), 5.07 (d, *J* = 10.6 Hz, 1H). **$^{13}$C NMR** (151 MHz, $CD_2Cl_2$) δ 159.9, 145.7, 145.6, 142.7, 142.6, 138.5, 133.7, 132.6, 132.2, 131.7, 131.2, 130.4, 130.0, 129.5, 129.1, 129.0, 128.7, 128.2, 128.0, 71.4, 59.2. **HRMS** (ESI): exact mass calculated for C25H21N4O2S [$(M+H)^+$] 441.1380, found 441.1400.

**(*R*)-EN-82:**

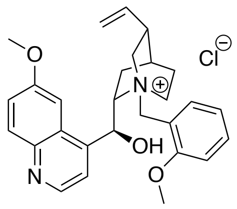

Catalyst A

**Catalyst A (1S,2S,4S,5R)-2-((*R*)-hydroxy(6-methoxyquinolin-4-yl)methyl)-1-(2-methoxybenzyl)-5-vinylquinuclidin-1-ium chloride** (for (*R*)-EN-82): Thionyl chloride (1.0 mL, 14 mmol) was added dropwise to a solution of 2-methoxybenzyl alcohol (967 mg, 7.0 mmol) in DCM (20 mL) at 0 °C, and the solution was then stirred at rt for 4h. Volatiles were then evaporated to provide the benzyl chloride. Quinidine (324 mg, 1.0 mmol) was added to a solution of 2-methoxybenzyl chloride (313 mg, 2.0 mmol) in toluene (5 mL) and the mixture was stirred at 80 °C for 19h. After cooling to rt, the mixture was concentrated and and purified by silica gel chromatography (0-4% MeOH/DCM) to provide the title compound (150 mg, 0.31 mmol, 31%) as a maroon foam. **LC/MS** calc. 445.2, found 455.2. **$^1$H NMR** (400 MHz, DMSO-$d_6$) δ 8.82 (d, *J* = 4.6 Hz, 1H), 8.02 (d, *J* = 9.2 Hz, 1H), 7.77 (d, *J* = 4.6 Hz, 1H), 7.67 (d, *J* = 7.7 Hz, 1H), 7.58 (t, *J* = 7.9 Hz, 1H), 7.49 (d, *J* = 9.3 Hz, 1H), 7.39 (s, 1H), 7.25 (d, *J* = 8.3 Hz, 1H), 7.15 (t, *J* = 7.4 Hz, 1H), 6.98 (d, *J* = 3.6 Hz, 1H), 6.56 (s, 1H), 6.05 (ddd, *J* = 17.5, 10.7, 7.3 Hz, 1H), 5.77 (s, 1H), 5.29 – 5.17 (m, 2H), 5.06 (d, *J* = 12.6 Hz, 1H), 4.75 (d, *J* = 12.5 Hz, 1H), 4.21 (t, *J* = 10.5 Hz, 1H), 4.09 (s, 3H), 3.90 (s, 3H), 3.47 (t, *J* = 11.6 Hz, 1H), 2.95 (q, *J* = 10.1 Hz, 1H), 2.65 (d, *J* = 9.2 Hz, 1H), 2.38 (t, *J* = 11.6 Hz, 1H), 2.09 (s, 1H), 1.88 (s, 1H), 1.82 – 1.71 (m, 2H), 1.10 – 0.97 (m, 1H). Procedure followed from Mahé *et al.* Enantioselective Phase-Transfer Catalysis: Synthesis of Pyrazolines. *Angewandte Chemie International Edition* **2010**, *49* (39), 7072–7075. https://doi.org/10.1002/anie.201002485.

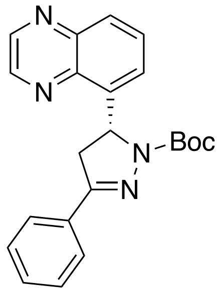


***tert*-butyl (*R*)-3-phenyl-5-(quinoxalin-5-yl)-4,5-dihydro-1*H*-pyrazole-1-carboxylate**: Following general procedure I and Catalyst A, (*E*)-1-phenyl-3-(quinoxalin-5-yl)prop-2-en-1-one (130 mg, 0.50 mmol) were combined in THF (1 mL) under nitrogen and stirred at 0 °C for 18 h. The mixture was diluted with EtOAc, filtered to remove salts, concentrated and purified by silica gel chromatography (0-35% EtOAc/Hex) to provide the pyrazoline (61 mg, 0.16 mmol, 31%) as a white solid. **LC/MS** calc. 375.15, found 375.2. **$^{1}$H NMR** (400 MHz, CDCl3) δ 8.97 – 8.91 (m, 2H), 8.09 (d, *J* = 8.4 Hz, 1H), 7.84 – 7.75 (m, 3H), 7.69 (d, *J* = 7.3 Hz, 1H), 7.44 – 7.39 (m, 3H), 6.65 (dd, *J* = 12.2, 5.4 Hz, 1H), 4.03 (dd, *J* = 17.3, 12.0 Hz, 1H), 3.17 (dd, *J* = 17.3, 5.4 Hz, 1H), 1.30 (s, 9H).

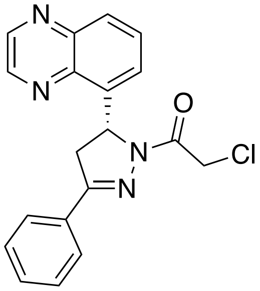


(*R*)-EN82

**(*R*)-EN-82 - (*R*)-2-chloro-1-(3-phenyl-5-(quinoxalin-5-yl)-4,5-dihydro-1*H*-pyrazol-1-yl)ethan-1-one** - Following general procedure J, tert-butyl (*R*)-3-phenyl-5-(quinoxalin-5-yl)-4,5-dihydro-1H-pyrazole-1-carboxylate (40 mg, 0.11 mmol) was converted to **(*R*)-EN-82** (22 mg, 0.064 mmol, 58%) as a white solid. **HRMS** calc. 351.1007, found 351.1007. **$^{1}$H NMR** (400 MHz, CDCl3) δ 8.92 (dd, *J* = 16.1, 1.7 Hz, 2H), 8.09 (d, *J* = 8.5 Hz, 1H), 7.84 – 7.71 (m, 3H), 7.59 (d, *J* = 7.2 Hz, 1H), 7.52 – 7.42 (m, 3H), 6.72 (dd, *J* = 11.9, 5.1 Hz, 1H), 4.81 (d, *J* = 13.4 Hz, 1H), 4.64 (d, *J* = 13.5 Hz, 1H), 4.07 (dd, *J* = 17.9, 11.9 Hz, 1H), 3.25 (dd, *J* = 17.9, 5.1 Hz, 1H). **$^{13}$C NMR** (151 MHz, DMSO) δ 163.8, 156.8, 146.3, 145.3, 143.1, 139.7, 139.5, 131.2, 131.2, 130.5, 129.2, 129.0, 127.4, 125.8, 56.6, 43.0, 42.3.

**(*S*)-EN082:**

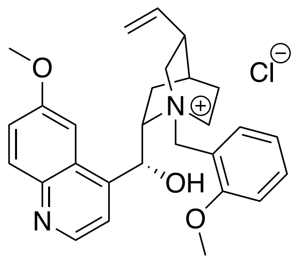


Catalyst B

**Catalyst B (1*S*,2*S*,4*S*,5*R*)-2-((*S*)-hydroxy(6-methoxyquinolin-4-yl)methyl)-1-(2-methoxybenzyl)-5-vinylquinuclidin-1-ium chloride (for *S*-EN082)** was synthesized from quinine, otherwise identically to Catalyst A, according to Mahé *et al.* Enantioselective Phase-Transfer Catalysis: Synthesis of Pyrazolines. *Angewandte Chemie International Edition* **2010**, *49* (39), 7072–7075. https://doi.org/10.1002/anie.201002485.

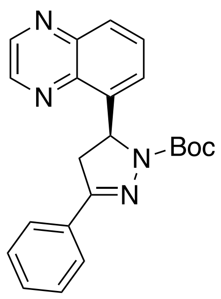

***tert*-butyl (*S*)-3-phenyl-5-(quinoxalin-5-yl)-4,5-dihydro-1*H*-pyrazole-1-carboxylate:** General Procedure I was followed from (*E*)-1-phenyl-3-(quinoxalin-5-yl)prop-2-en-1-one (130 mg, 0.50 mmol), using Catalyst B to provide the title compound (122 mg, 0.32 mmol, 65%) as a white solid. **LC/MS** calc. 375.17, found 375.2. **$^{1}$H NMR** (400 MHz, CDCl3) δ 8.94 (d, *J* = 10.0 Hz, 2H), 8.09 (d, *J* = 8.5 Hz, 1H), 7.84 – 7.76 (m, 3H), 7.69 (d, *J* = 7.3 Hz, 1H), 7.45 – 7.35 (m, 3H), 6.64 (dd, *J* = 12.2, 5.3 Hz, 1H), 4.02 (dd, *J* = 18.0, 12.7 Hz, 1H), 3.17 (d, *J* = 18.0 Hz, 1H), 1.30 (s, 9H).

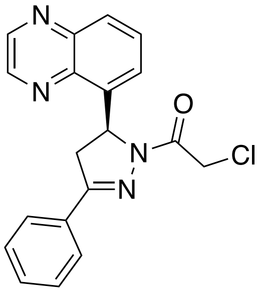

(S)-EN82

**(*S*)-EN-82 - (*S*)-2-chloro-1-(3-phenyl-5-(quinoxalin-5-yl)-4,5-dihydro-1*H*-pyrazol-1-yl)ethan-1-one -** Following general procedure J, tert-butyl (*R*)-3-phenyl-5-(quinoxalin-5-yl)-4,5-dihydro-1H-pyrazole-1-carboxylate (40 mg, 0.11 mmol) was converted to **(*S*)-EN-82** (21.3 mg, 0.061 mmol, 47%) as a white solid. **HRMS** calc. 351.1007, found 351.1002. **$^{1}$H NMR** (400 MHz, DMSO-$d_6$) δ 9.04 (d, *J* = 3.9 Hz, 2H), 8.06 (d, *J* = 8.5 Hz, 1H), 7.85 – 7.78 (m, 3H), 7.56 (d, *J* = 7.2 Hz, 1H), 7.50 – 7.43 (m, 3H), 6.56 (dd, *J* = 11.9, 4.9 Hz, 1H), 4.92 (d, *J* = 13.9 Hz, 1H), 4.78 (d, *J* = 13.9 Hz, 1H), 4.08 (dd, *J* = 18.2, 12.0 Hz, 1H), 3.25 (dd, *J* = 18.1, 5.0 Hz, 1H). **$^{13}$C NMR** (151 MHz, DMSO) δ 163.8, 156.8, 146.3, 145.3, 143.1, 139.7, 139.5, 131.2, 131.2, 130.6, 129.3, 129.0, 127.4, 125.8, 56.6, 43.0, 42.3.

**NJH-01-123:**

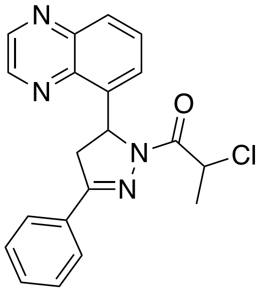

NJH-01-123

**2-chloro-1-(3-phenyl-5-(quinoxalin-5-yl)-4,5-dihydro-1H-pyrazol-1-yl)propan-1-one:** (*E*)-1-phenyl-3-(quinoxalin-5-yl)prop-2-en-1-one (55mg, 0.20 mmol) was converted via General Procedure F and purified by silica gel chromatography (0-25% EtOAc/Hex) to give **NJH-01-123** (43 mg, 0.12 mmol, 59%) as a cream-colored solid. **$^{1}$H NMR** (400 MHz, CDCl3) δ 8.93 (dd, *J* = 11.0, 1.9 Hz, 2H), 8.09 (dd, *J* = 8.5, 1.4 Hz, 1H), 7.83 – 7.79 (m, 2H), 7.77 (d, *J* = 7.7 Hz, 1H), 7.65 (d, *J* = 7.1 Hz, 1H), 7.50 – 7.43 (m, 3H), 6.77 (dd, *J* = 11.9, 4.9 Hz, 1H), 5.60 (q, *J* = 6.8 Hz, 1H), 4.07 (dd, *J* = 17.9, 11.9 Hz, 1H), 3.22 (dd, *J* = 17.9, 4.9 Hz, 1H), 1.79 (d, *J* = 6.8 Hz, 3H). **HRMS** calc. 365.1164, found 365.1173.

**NJH-01-124:**

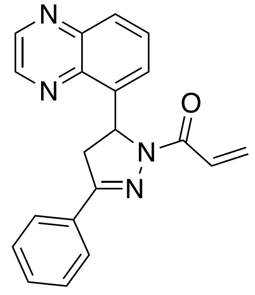

NJH-01-124

**1-(3-phenyl-5-(quinoxalin-5-yl)-4,5-dihydro-1H-pyrazol-1-yl)prop-2-en-1-one (NJH-01-124):** (*E*)-1-phenyl-3-(quinoxalin-5-yl)prop-2-en-1-one (55 mg, 0.20 mmol) was converted via General Procedure F and purified by silica gel chromatography (0-50% EtOAc/Hex) to give **NJH-01-124** (31 mg, 0.094 mmol, 47%) as a pale yellow solid. **HRMS** calc. 329.1397, found 329.1403. $^{1}$**H NMR** (400 MHz, CDCl3) δ 8.93 (d, *J* = 7.1 Hz, 2H), 8.07 (d, *J* = 8.6 Hz, 1H), 7.86 – 7.70 (m, 3H), 7.60 – 7.39 (m, 5H), 6.82 (dd, *J* = 11.8, 4.9 Hz, 1H), 6.51 (d, *J* = 17.2 Hz, 1H), 5.86 (d, *J* = 9.7 Hz, 1H), 4.06 (dd, *J* = 17.9, 12.0 Hz, 1H), 3.21 (dd, *J* = 17.8, 5.0 Hz, 1H). $^{13}$**C NMR** (151 MHz, DMSO) δ 162.5, 156.1, 146.3, 145.2, 143.1, 140.0, 139.7, 131.5, 131.0, 130.6, 129.2, 128.8, 128.5, 127.3, 125.7, 56.3, 42.1.

**NJH-01-145:**

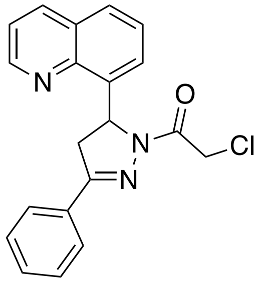

NJH-01-145

**2-chloro-1-(3-phenyl-5-(quinolin-8-yl)-4,5-dihydro-1H-pyrazol-1-yl)ethan-1-one (NJH-01-145)**: General Procedure A was followed starting from quinoline-8-carbaldehyde (157 mg, 1.0 mmol). After stirring at room temperature for 15 h, the solution was diluted with water and extracted with EtOAc (3x5mL). Extracts were combined, washed with NaCl solution (sat. aqueous), dried over $Na_2SO_4$, concentrated and purified by silica gel chromatography (0-60% EtOAc/Hex) to obtain the chalcone (85 mg, 0.33 mmol, 33%) as an orange oil.

This chalcone ((*E*)-1-phenyl-3-(quinolin-8-yl)prop-2-en-1-one) (81 mg, 0.31 mmol) was converted via General Procedure F and purified by silica gel chromatography (0-50% EtOAc/Hex) to give **NJH-01-145** (35 mg, 0.10 mmol, 32%) as a white solid. **HRMS** calc. 350.1055, found 350.1057. $^{1}$**H NMR** (400 MHz, CDCl3) δ 8.98 (dd, *J* = 4.2, 1.8 Hz, 1H), 8.21 (dd, *J* = 8.3, 1.8 Hz, 1H), 7.79 (d, *J* = 8.2 Hz, 3H), 7.57 – 7.39 (m, 6H), 6.78 (dd, *J* = 11.7, 4.9 Hz, 1H), 4.75 (dd, *J* = 73.5, 13.2 Hz, 2H), 4.10 (dd, *J* = 18.0, 11.8 Hz, 1H), 3.26 (dd, *J* = 17.9, 4.9 Hz, 1H). $^{13}$**C NMR** (151 MHz, DMSO) δ 163.7, 156.8, 150.4, 144.9, 138.7, 137.0, 131.3, 131.1, 129.2, 128.8, 128.1, 127.4, 126.8, 124.8, 122.1, 57.1, 43.0, 42.5.

**NJH-01-147:**

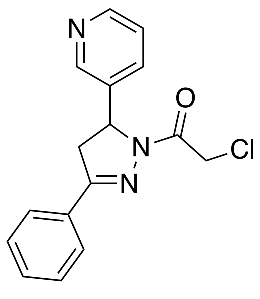

NJH-01-147

**2-chloro-1-(3-phenyl-5-(pyridin-3-yl)-4,5-dihydro-1*H*-pyrazol-1-yl)ethan-1-one(NJH-01-147):** General Procedure A was followed starting from pyridine-3-carbaldehyde (96 uL, 1.0 mmol). After stirring at room temperature for 15h, the solution was diluted with water and extracted with EtOAc (3x5mL). Extracts were combined, washed with NaCl solution (sat. aqueous), dried over $Na_2SO_4$, concentrated and purified by silica gel chromatography (0-70% EtOAc/Hex) to obtain the chalcone (50 mg, 0.24 mmol, 24%) as a clear colorless oil.

This chalcone (*E*)-1-phenyl-3-(pyridin-3-yl)prop-2-en-1-one (50 mg, 0.24 mmol) was converted via General Procedure F and purified by silica gel chromatography (0-80% EtOAc/Hex) to give **NJH-01-147** (21 mg, 0.070 mmol, 29%) as a white solid. **HRMS** calc. 300.0898, found 300.0917. **$^1$H NMR** (400 MHz, CDCl3) δ 8.62 (d, *J* = 2.4 Hz, 1H), 8.59 (d, *J* = 4.8 Hz, 1H), 7.80 (d, *J* = 7.7 Hz, 2H), 7.61 (d, *J* = 7.9 Hz, 1H), 7.56 – 7.47 (m, 3H), 7.32 (d, *J* = 4.9 Hz, 1H), 5.67 (dd, *J* = 11.9, 4.8 Hz, 1H), 4.63 (dd, *J* = 22.3, 13.8 Hz, 2H), 3.90 (dd, *J* = 17.9, 11.8 Hz, 1H), 3.29 (dd, *J* = 17.9, 4.9 Hz, 1H). **$^{13}$C NMR** (151 MHz, DMSO) δ 163.9, 156.3, 149.2, 148.1, 137.3, 133.9, 131.3, 131.0, 129.3, 127.5, 124.3, 58.5, 42.9, 42.1.

**NJH-01-158:**

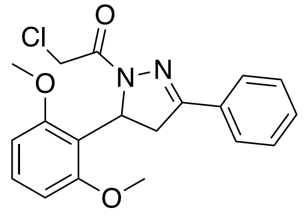

NJH-01-158

**2-chloro-1-(5-(2,6-dimethoxyphenyl)-3-phenyl-4,5-dihydro-1*H*-pyrazol-1-yl)ethan-1-one (NJH-01-158):** General Procedure A was followed starting from 2,6-dimethoxybenzaldehyde (332mg, 2.0 mmol). After stirring at rt 15h, the mixture was diluted with water and extracted with EtOAc (3x5mL). Extracts were combined, washed with NaCl solution (sat. aqueous), dried over $Na_2SO_4$, concentrated and purified by silica gel chromatography (0-30% EtOAc/Hex) to obtain the chalcone (525 mg, 1.96 mmol, 98%) as a colorless oil. This chalcone (*E*)-3-(2,6-dimethoxyphenyl)-1-phenylprop-2-en-1-one (100 mg, 0.37 mmol) was converted via General Procedure F and purified by silica gel chromatography (0-50% EtOAc/Hex) to give **NJH-01-158** (89 mg, 0.25 mmol, 67%) as a white solid. **HRMS** calc. 359.1157, found 359.1139. **$^1$H NMR** (400 MHz, CDCl3) δ 7.85 – 7.73 (m, 2H), 7.54 – 7.42 (m, 3H), 7.23 (t, *J* = 8.3 Hz, 1H), 6.58 (d, *J* = 8.3 Hz, 2H), 6.21 (dd, *J* = 12.6, 6.2 Hz, 1H), 4.58 (d, *J* = 2.1 Hz, 2H), 3.80 (s, 6H), 3.60 (dd, *J* = 17.3, 12.6 Hz, 1H), 3.27 (dd, *J* = 17.3, 6.3 Hz, 1H). **$^{13}$C NMR** (151 MHz, DMSO) δ 162.7, 158.4, 156.0, 131.8, 130.7, 129.4, 129.2, 127.1, 117.2, 105.2, 56.5, 51.7, 42.9.

**NJH-01-159:**

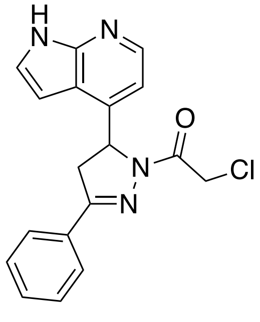

NJH-01-159

**2-chloro-1-(3-phenyl-5-(1H-pyrrolo[2,3-b]pyridin-4-yl)-4,5-dihydro-1H-pyrazol-1-yl)ethan-1-one (NJH-01-159):** 1*H*-pyrrolo[2,3-*b*]pyridine-4-carbaldehyde: n-BuLi (3.0 mL, 7.62 mmol, 2.5 M in hexanes) was added dropwise to a solution of 4-bromo-1H-pyrrolo[2,3-b]pyridine (500 mg, 2.54 mmol) in THF (20 mL) at -78 ºC and stirred at that temperature for 30 min. DMF (988 uL, 12.7 mmol) was added and the reaction was stirred at rt for 1h before being cooled to -78 ºC and quenched with sat. aq. $NH_4Cl$. The mixture was then warmed to rt, diluted with water and extracted with EtOAc (3x10 mL). Extracts were combined, washed with NaCl solution (sat. aqueous), dried over $Na_2SO_4$, concentrated and purified by silica gel chromatography (0-40% EtOAc/Hex) to obtain the aldehyde (178 mg, 1.22 mmol, 48%) as a yellow solid.

General Procedure A was followed starting from 1H-pyrrolo[2,3-b]pyridine-4-carbaldehyde (75 mg, 0.51 mmol). After stirring at rt 4h, water (15 mL) was added and the precipitate filtered to provide the chalcone (93 mg, 0.38 mmol, 74%) as a bright yellow solid. This chalcone (*E*)-1-phenyl-3-(1H-pyrrolo[2,3-b]pyridin-4-yl)prop-2-en-1-one was converted via General Procedure F and purified by silica gel chromatography (0-60% EtOAc/Hex) to give **NJH-01-159** (15 mg, 0.044 mmol, 22%) as a yellow solid. **LC/MS** calc. 339.1, found 339.1. **$^1$H NMR** (400 MHz, CDCl3) δ 9.55 (s, 1H), 8.31 (d, *J* = 4.9 Hz, 1H), 7.81 (d, *J* = 7.8 Hz, 2H), 7.55 – 7.46 (m, 3H), 7.35 (s, 1H), 7.04 (d, *J* = 5.0 Hz, 1H), 6.45 (d, *J* = 3.3 Hz, 1H), 5.94 (dd, *J* = 12.1, 5.4 Hz, 1H), 4.67 (s, 2H), 3.94 (dd, *J* = 17.8, 12.1 Hz, 1H), 3.39 (dd, *J* = 17.8, 5.4 Hz, 1H).

**NJH-01-161:**

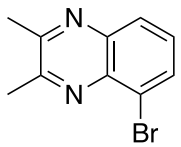

**5-bromo-2,3-dimethylquinoxaline**: 3-bromobenzene-1,2-diamine (1.0 g, 5.35 mmol) and diacetyl (517 mL, 5.89 mmol) were added to EtOH (20 mL), and the solution was stirred at reflux for 1h and cooled to rt. The mixture was then concentrated to dryness and the crude was redissolved in MeOH 95 mL. Water (90 mL) was then added and the precipitate filtered to provide the quinoxaline (1.17 g, 4.94 mmol, 92%) as a tan solid. **LC/MS calc.** 237.0, found 236.9. **$^1$H NMR** (400 MHz, CDCl3) δ 8.01 (t, *J* = 8.7 Hz, 2H), 7.57 (td, *J* = 7.9, 1.8 Hz, 1H), 2.85 (s, 3H), 2.81 (s, 3H).

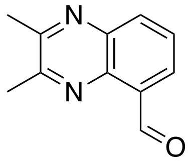

**2,3-dimethylquinoxaline-5-carbaldehyde:** 5-bromo-2,3-dimethylquinoxaline (474 mg, 2.0 mmol) was dissolved in THF (10 mL) and cooled to -78 ºC. n-BuLi in hexanes (960 mL, 2.4 mmol) was then added dropwise, and the mixture stirred at -78 ºC for 30 min. DMF (311 mL, 4.0 mmol) was then added at -78 ºC and the reaction stirred at rt for 1 hour. The reaction was then quenched with sat. aq. $NH_4Cl$ at -78 ºC, warmed to rt, diluted with water and extracted with EtOAc (3x10mL). Extracts were combined, washed with NaCl solution (sat. aqueous), dried over $Na_2SO_4$, concentrated and purified by silica gel chromatography (0-50% EtOAc/Hex) to obtain the aldehyde (169 mg, 0.91 mmol, 45%). **LC/MS** calc. 187.1, found 187.1. **$^1$H NMR** (400 MHz, CDCl3) δ 11.41 (s, 1H), 8.04 (dd, *J* = 6.3, 3.5 Hz, 1H), 7.83 (t, *J* = 7.7 Hz, 1H), 7.72 (dd, *J* = 6.4, 3.4 Hz, 1H), 2.84 (s, 3H), 2.82 (s, 3H).

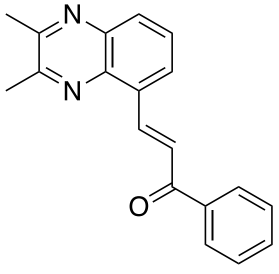

**(*E*)-3-(2,3-dimethylquinoxalin-5-yl)-1-phenylprop-2-en-1-one**: General Procedure A was followed starting from 2,3-dimethylquinoxaline-5-carbaldehyde (93 mg, 0.50 mmol). After stirring at room temperature for 5h, the mixture was diluted with water and extracted with EtOAc (3x5mL). Extracts were combined, washed with NaCl solution (sat. aqueous), dried over $Na_2SO_4$, concentrated and purified by silica gel chromatography (0-25% EtOAc/Hex) to obtain the chalcone (59 mg, 0.21 mmol, 41%). **LC/MS** calc. 289.1, found 289.1. **$^1$H NMR** (400 MHz, CDCl3) δ 8.96 (d, *J* = 16.0 Hz, 1H), 8.17 – 8.11 (m, 2H), 8.09 (d, *J* = 7.4 Hz, 2H), 8.04 (s, 1H), 7.76 (t, *J* = 7.9 Hz, 1H), 7.68 – 7.63 (m, 1H), 7.61 – 7.54 (m, 2H), 2.84 (s, 3H), 2.80 (s, 3H).

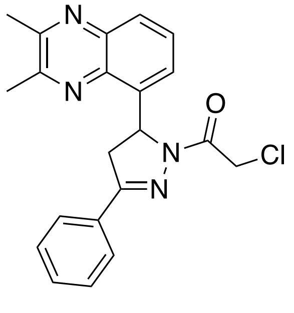

NJH-01-161

**2-chloro-1-(5-(2,3-dimethylquinoxalin-5-yl)-3-phenyl-4,5-dihydro-1*H*-pyrazol-1-yl)ethan-1-one (NJH-01-161):** (*E*)-3-(2,3-dimethylquinoxalin-5-yl)-1-phenylprop-2-en-1-one (30 mg, 0.10 mmol) was converted via General Procedure F and purified by silica gel chromatography (0-45% EtOAc/Hex) to give **NJH-01-161** (30 mg, 0.080 mmol, 80%) as a white solid. HRMS calc. 379.1320, found 379.1342. **$^1$H NMR** (400 MHz, CDCl3) δ 7.85 (d, *J* = 8.3 Hz, 1H), 7.71 (s, 2H), 7.54 (t, *J* = 7.7 Hz, 1H), 7.39 (s, 4H), 6.55 (d, *J* = 11.8 Hz, 1H), 4.62 (dd, *J* = 42.9, 13.4 Hz, 2H), 3.92 (dd, *J* = 17.9, 12.7 Hz, 1H), 3.18 (dd, *J* = 17.4, 4.2 Hz, 1H), 2.67 (s, 3H), 2.61 (s, 3H). **$^{13}$C NMR** (151 MHz, DMSO) δ 163.7, 156.8, 154.5, 153.6, 141.2, 138.5, 137.8, 131.4, 131.1, 129.2, 129.0, 127.9, 127.4, 57.0, 42.9, 42.3, 23.6, 23.2.

**NJH-01-166:**

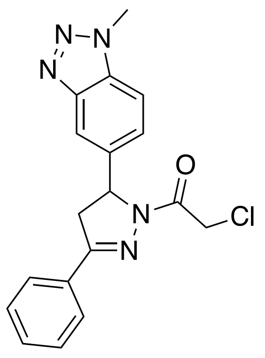

**(1-methyl-1*H*-benzo[*d*][1,2,3]triazol-5-yl)methanol:** $LiAlH_4$ (5.64 mmol, 1M in THF) was added dropwise to a solution of 1-methylbenzotriazole-5-carboxylic acid (500 mg, 2.52 mmol) in THF (10 mL) at 0 °C. The mixture was allowed to warm to rt and stirred for 5 h. The mixture was then diluted with $Et_2O$ (40mL), and 0.2 mL water was added at 0 °C, followed by 0.6 mL 1M NaOH aq., 0.2 mL water, and finally $Na_2SO_4$ after 15 min at rt. The mixture was filtered, concentrated and purified by silica gel chromatography (0-80% EtOAc/Hex) to obtain the alcohol (308 mg, 1.89 mmol, 67%). **LC/MS** calc. 164.1, found 164.1.

**1-methyl-1*H*-benzo[*d*][1,2,3]triazole-5-carbaldehyde:** (1-methyl-1*H*-benzo[*d*][1,2,3]triazol-5-yl)methanol (200 mg, 1.22 mmol) and 4-methylmorpholine N-oxide (500 mg, 4.27 mmol) were combined in DCM (5 mL) with 4Å MS. TPAP (42 mg, 0.12 mmol) was then added and the mixture stirred as 2 hours before the alcohol was consumed. The mixture was diluted with $Et_2O$ and filtered to remove NMO, filtrate concentrated and purified by silica gel chromatography (0-30% EtOAc/Hex) to obtain the aldehyde (138 mg, 0.86 mmol, 70%). **LC/MS** calc. 162.1, found 162.0.

**2-chloro-1-(5-(1-methyl-1H-benzo[d][1,2,3]triazol-5-yl)-3-phenyl-4,5-dihydro-1H-pyrazol-1-yl)ethan-1-one (NJH-01-166):** General Procedure A was followed starting from 1-methyl-1H-benzo[d][1,2,3]triazole-5-carbaldehyde (80 mg, 0.50 mmol). After 16 h., water was added and the precipitate filtered to obtain the chalcone (117 mg, 0.44 mmol, 88%) as a tan powder. This chalcone (*E*)-3-(1-methyl-1*H*-benzo[*d*][1,2,3]triazol-5-yl)-1-phenylprop-2-en-1-one (40 mg, 0.15 mmol) was converted via General Procedure F and purified by silica gel chromatography (0-60% EtOAc/Hex) to give **NJH-01-166** (16 mg, 0.046 mmol, 31%) as a light yellow foam. **HRMS** calc. 354.1116, found 354.1093. $^{1}$**H NMR** (400 MHz, CDCl3) δ 7.99 (s, 1H), 7.82 (d, *J* = 7.8 Hz, 2H), 7.59 – 7.43 (m, 5H), 5.85 – 5.76 (m, 1H), 4.64 (d, *J* = 4.8 Hz, 2H), 4.32 (s, 3H), 3.93 (dd, *J* = 18.0, 11.2 Hz, 1H), 3.34 (dd, *J* = 17.9, 5.0 Hz, 1H). $^{13}$**C NMR** (151 MHz, DMSO) δ 163.8, 156.2, 145.7, 138.0, 133.3, 131.2, 131.1, 129.3, 127.4, 125.8, 116.2, 111.7, 60.5, 43.0, 42.6, 34.7.

**NJH-01-181:**

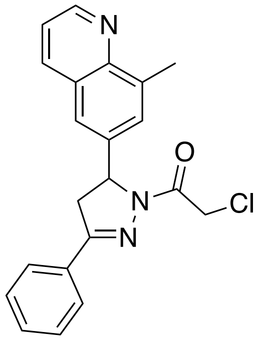

NJH-01-181

**2-chloro-1-(5-(8-methylquinolin-6-yl)-3-phenyl-4,5-dihydro-1H-pyrazol-1-yl)ethan-1-one (NJH-01-181):** 6-bromo-8-methylquinoline (500 mg, 2.25 mmol) was converted to 8-methylquinoline-6-carbaldehyde via General Procedure K and purified by silica gel chromatography (0-40% EtOAc/Hex) to provide the aldehyde (184 mg, 1.08 mmol, 48%) as a pale orange solid.

Then, General Procedure A was followed starting from 8-methylquinoline-6-carbaldehyde (100mg, 0.58 mmol). After stirring overnight water (15 mL) was added, and the solids filtered to provide the chalcone (134 mg, 0.49 mmol, 85%) as a tan solid. (*E*)-3-(8-methylquinolin-6-yl)-1-phenylprop-2-en-1-one (75 mg, 0.27 mmol) was converted via General Procedure F and purified by silica gel chromatography (0-45% EtOAc/Hex) to give **NJH-01-181** (49 mg, 0.13 mmol, 49%) as a white powder. **HRMS** calc. 364.1211, found 364.1219. $^{1}$**H NMR** (400 MHz, DMSO-$d_6$) δ 8.94 – 8.85 (m, 1H), 8.33 (d, *J* = 8.4 Hz, 1H), 7.90 – 7.81 (m, 2H), 7.67 (s, 1H), 7.56 – 7.47 (m, 5H), 5.74 (dd, *J* = 11.9, 5.0 Hz, 1H), 4.81 (dd, *J* = 32.2, 13.6 Hz, 2H), 3.99 (dd, *J* = 17.9, 11.4 Hz, 1H), 3.34 (dd, *J* = 17.9, 5.4 Hz, 1H), 2.71 (s, 3H). $^{13}$**C NMR** (151 MHz, DMSO) δ 163.9, 156.2, 150.0, 146.6, 139.6, 137.8, 136.8, 131.2, 131.1, 129.3, 128.2, 127.7, 127.5, 122.8, 122.1, 60.5, 43.0, 42.5, 18.3.

**NJH-01-182:**

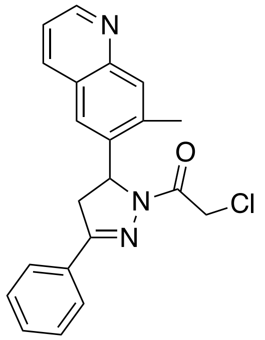

NJH-01-182

**2-chloro-1-(5-(7-methylquinolin-6-yl)-3-phenyl-4,5-dihydro-1H-pyrazol-1-yl)ethan-1-one (NJH-01-182):** 7-methylquinoline-6-carbaldehyde:6-bromo-8-methylquinoline (500 mg, 2.25 mmol) was converted to 7-methylquinoline-6-carbaldehyde via general procedure K and purified by silica gel chromatography (0-50% EtOAc/Hex) to provide the aldehyde (275 mg, 1.61 mmol, 71%) as a pale orange solid. LC/MS calc. 172.1, found 172.1.

General Procedure A was followed starting from 7-methylquinoline-6-carbaldehyde (100 mg, 0.58 mmol). The reaction mixture was stirred for 16 hours at rt and an immiscible orange oil formed. Water was added and the mixture extracted with EtOAc (3x5mL). Extracts were combined, washed with NaCl solution (sat. aqueous), dried over $Na_2SO_4$, concentrated, and purified by silica gel chromatography (0 to 50% EtOAc/Hex.) to obtain the chalcone (151 mg, 0.55 mmol, 95%) as a yellow gummy solid. This chalcone (*E)*-3-(7-methylquinolin-6-yl)-1-phenylprop-2-en-1-one (75 mg, 0.27 mmol) was converted via General Procedure F and purified by silica gel chromatography (0-65% EtOAc/Hex) to give **NJH-01-182** (25 mg, 0.069 mmol, 25%) as a white solid. **HRMS** calc. 364.1211, found 364.1222. **$^1$H NMR** (400 MHz, DMSO-$d_6$) δ 8.83 (dd, *J* = 4.2, 1.7 Hz, 1H), 8.26 (dd, *J* = 8.2, 1.8 Hz, 1H), 7.91 (s, 1H), 7.87 – 7.82 (m, 2H), 7.57 (s, 1H), 7.51 – 7.41 (m, 4H), 5.87 (dd, *J* = 11.7, 4.9 Hz, 1H), 4.87 (dd, *J* = 37.1, 13.8 Hz, 2H), 4.05 (dd, *J* = 18.1, 11.9 Hz, 1H), 3.24 (dd, *J* = 18.1, 4.9 Hz, 1H), 2.62 (s, 3H). **$^{13}$C NMR** (151 MHz, DMSO) δ 163.8, 156.3, 150.9, 147.5, 139.3, 137.4, 136.3, 131.2, 131.1, 130.0, 129.3, 127.4, 126.8, 123.2, 121.4, 58.0, 43.0, 41.8, 20.1.

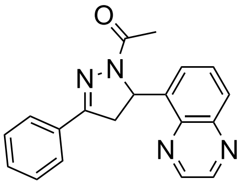

LEB-02-150

**1-(3-phenyl-5-(quinoxalin-5-yl)-4,5-dihydro-1H-pyrazol-1-yl)ethan-1-one (LEB-02-150).** General Procedure A was followed starting from acetophenone (0.40 mmol) and quinoxaline-5-carbaldehyde (0.40 mmol). The corresponding chalcone precipitated out during the reaction and was filtered by gravity filtration to yield product (0.20 mmol) as a light yellow solid. General Procedure F was followed starting from the above product (1.0 equiv, 0.20 mmol). Purification by flash column chromatography (EtOAc/hexane 50:50) yielded the chloroacetamide **LEB-02-150** (0.08 mmol, 22.1% yield across all steps). **$^1$H NMR** (400 MHz, CDCl3) δ 8.97 – 8.89 (m, 2H), 8.07 (d, *J* = 8.4 Hz, 1H), 7.82 – 7.71 (m, 3H), 7.56 – 7.50 (m, 1H), 7.48 – 7.40 (m, 3H), 7.30 (s, 3H), 6.73 (dd, *J* = 11.9, 4.8 Hz, 1H), 4.05 (dd, *J* = 17.9, 12.1 Hz, 1H), 3.18 (dd, *J* = 17.5, 4.9 Hz, 1H). **$^{13}$C NMR** (151 MHz, DMSO) δ 168.0, 155.2, 146.2, 145.2, 143.1, 140.3, 139.7, 131.6, 130.8, 130.6, 129.2, 128.7, 127.1, 125.5, 56.0, 42.4, 22.2. **HRMS** (ESI): exact mass calculated for $C_{19}H_{16}N_4O$ [$(M+H)^+$] 317.1397, found 317.1405.

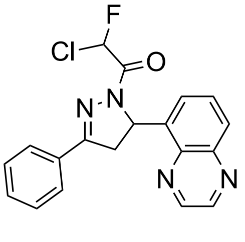

LEB-02-157

**2-chloro-2-fluoro-1-(3-phenyl-5-(quinoxalin-5-yl)-4,5-dihydro-1*H*-pyrazol-1-yl)ethan-1-one (LEB-02-157).** General Procedure A was followed starting from acetophenone (0.40 mmol) and quinoxaline-5-carbaldehyde (0.40 mmol). The corresponding chalcone precipitated out during the reaction and was filtered by gravity filtration to yield product (0.38 mmol) as a yellow solid. Hydrazine monohydrate (2.0 equiv) was added to a suspension of the above product (1.0 equiv, 0.38 mmol) in EtOH (0.3 M). The resulting reaction mixture was stirred at reflux temperature for 4 h before it was concentrated under reduced pressure. The crude pyrazoline (1.0 equiv, 0.36 mmol) was added to a screw-cap oven-dried vial charged with a stir-bar, chlorofluoroacetate (1.2 equiv, 0.44 mmol), EDC hydrochloride (1.2 equiv, 0.44 mmol), HOBt (1.2 equiv, 0.44 mmol), DIEA (2.5 equiv, 0.92 mmol), and 2mL DMF. Reaction proceeded overnight, and was extracted in EtOAc, washed 3X with NaCl solution (sat. aqueous), and purified via flash column chromatography (EtOAc/hexane 50:50), yielding the corresponding chlorofluoroacetamide **LEB-02-157** (0.026 mmol, 6.4% yield across all steps). **$^1$H NMR** (400 MHz, CDCl3) δ 8.97 – 8.84 (m, 2H), 8.13 – 8.08 (m, 1H), 8.06 – 7.95 (m, 1H), 7.83 – 7.75 (m, 3H), 7.63 – 7.57 (m, 1H), 7.54 – 7.44 (m, 3H), 6.70 (dd, *J* = 11.7, 5.1 Hz, 1H), 4.16 – 4.04 (m, 1H), 3.36 – 3.22 (m, 1H). **HRMS** (ESI): exact mass calculated for $C_{19}H_{14}ClFN_4O$ [(M+H)$^+$] 369.0913, found 369.0914.

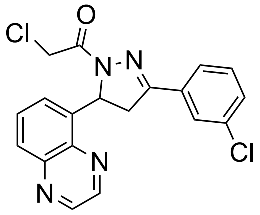


LEB-02-172

**2-chloro-1-(3-(3-chlorophenyl)-5-(quinoxalin-5-yl)-4,5-dihydro-1*H*-pyrazol-1-yl)ethan-1-one (LEB-02-172).** General Procedure A was followed starting from 3-chloroacetophenone (1.2 equiv, 0.76 mmol) and quinoxaline-5-carbaldehyde (1.0 equiv, 0.63 mmol). The corresponding chalcone precipitated out during the reaction and was filtered by gravity filtration to yield product (0.50 mmol) as a yellow solid. General Procedure F was followed starting from the above product (1.0 equiv, 0.50 mmol). Purification by flash column chromatography (EtOAc/hexane 50:50) yielded the chloroacetamide **LEB-02-172** (0.058 mmol, 9.2% yield across all steps). **$^1$H NMR** (400 MHz, CDCl3) δ 8.92 (dt, *J* = 21.3, 1.8 Hz, 2H), 8.10 (dd, *J* = 8.5, 1.7 Hz, 1H), 7.81 – 7.74 (m, 2H), 7.68 – 7.62 (m, 1H), 7.58 (d, *J* = 7.3 Hz, 1H), 7.49 – 7.43 (m, 1H), 7.43 – 7.37 (m, 1H), 6.74 – 6.66 (m, 1H), 4.79 (dd, *J* = 13.5, 1.6 Hz, 1H), 4.63 (dd, *J* = 13.7, 1.4 Hz, 1H), 4.10 – 3.99 (m, 1H), 3.28 – 3.19 (m, 1H). **$^{13}$C NMR** (151 MHz, DMSO) δ 164.0, 155.6, 146.3, 145.3, 143.1, 139.7, 139.3, 134.1, 133.3, 131.2, 130.8, 130.5, 129.0, 126.9, 126.1, 56.9, 43.0, 42.2. **HRMS** (ESI): exact mass calculated for $C_{19}H_{14}Cl_2N_4O$ [(M+H)$^+$] 385.0617, found 385.063.

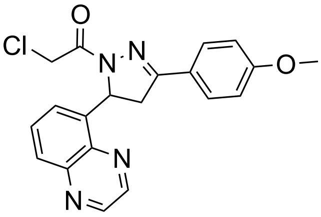


LEB-02-173

**2-chloro-1-(3-(4-methoxyphenyl)-5-(quinoxalin-5-yl)-4,5-dihydro-1*H*-pyrazol-1-yl)ethan-1-one (LEB-02-173).** General Procedure A was followed starting from 4-methoxyacetophenone (1.2 equiv, 0.76 mmol) and quinoxaline-5-carbaldehyde (1.0 equiv, 0.63 mmol). The corresponding chalcone precipitated out during the reaction and was filtered by gravity filtration to yield product (0.47 mmol) as a light yellow solid. General Procedure F was followed starting from the above product (1.0 equiv, 0.47 mmol). Purification by flash column chromatography (EtOAc/hexane 50:50) yielded the chloroacetamide **LEB-02-173** (0.025 mmol, 4.0% yield across all steps). **$^1$H NMR** (400 MHz, DMSO-$d_6$) δ 9.04 (q, *J* = 1.8 Hz, 2H), 8.05 (dd, *J* = 8.3, 1.4 Hz, 1H), 7.82 (dd, *J* = 8.5, 7.3 Hz, 1H), 7.77 – 7.70 (m, 2H), 7.53 (d, *J* = 7.2 Hz, 1H), 7.04 – 6.98 (m, 2H), 6.54 (dd, *J* = 11.9, 4.7 Hz, 1H), 4.90 (d, *J* = 13.8 Hz, 1H), 4.76 (d, *J* = 13.8 Hz, 1H), 4.04 (dd, *J* = 18.0, 11.9 Hz, 1H), 3.80 (s, 3H), 3.21 (dd, *J* = 18.0, 4.8 Hz, 1H). **HRMS** (ESI): exact mass calculated for $C_{20}H_{17}ClN_4O_2$ [(M+H)$^+$] 381.1113, found 381.1088.

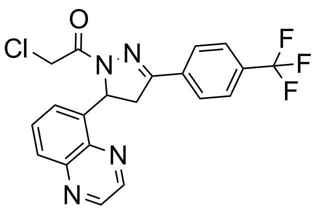

LEB-02-174

**2-chloro-1-(5-(quinoxalin-5-yl)-3-(4-(trifluoromethyl)phenyl)-4,5-dihydro-1*H*-pyrazol-1-yl)ethan-1-one (LEB-02-174).** General Procedure A was followed starting from 4-(trifluoromethyl)acetophenone (1.2 equiv, 0.76 mmol) and quinoxaline-5-carbaldehyde (1.0 equiv, 0.63 mmol). The corresponding chalcone precipitated out during the reaction and was filtered by gravity filtration to yield product (0.55 mmol) as a yellow solid. General Procedure F was followed starting from the above product (1.0 equiv, 0.55 mmol). Purification by flash column chromatography (EtOAc/hexane 50:50) yielded the chloroacetamide **LEB-02-174** (0.11 mmol, 18.9% yield across all steps). **$^1$H NMR** (400 MHz, DMSO-$d_6$) δ 9.05 – 9.00 (m, 2H), 8.00 (d, *J* = 8.2 Hz, 2H), 7.85 – 7.79 (m, 3H), 7.49 – 7.43 (m, 2H), 4.97 – 4.77 (m, 1H), 4.17 – 3.99 (m, 1H), 3.40 (s, 2H), 3.31 (dd, *J* = 18.2, 5.1 Hz, 1H). **HRMS** (ESI): exact mass calculated for $C_{20}H_{14}ClF_3N_4O$ [(M+H)$^+$] 419.0881, found 419.0865.

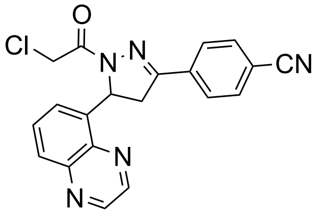

LEB-02-175

**4-(1-(2-chloroacetyl)-5-(quinoxalin-5-yl)-4,5-dihydro-1H-pyrazol-3-yl)benzonitrile (LEB-02-175).** General Procedure A was followed starting from 4-acetylbenzonitrile (1.2 equiv, 0.76 mmol) and quinoxaline-5-carbaldehyde (1.0 equiv, 0.63 mmol). The corresponding chalcone precipitated out during the reaction and was filtered by gravity filtration to yield product (0.57 mmol) as a yellow solid. General Procedure F was followed starting from the above product (1.0 equiv, 0.57 mmol). Purification by flash column chromatography (EtOAc/hexane 50:50) yielded the chloroacetamide **LEB-02-175** (0.050 mmol, 7.9% yield across all steps). **$^1$H NMR** (400 MHz, DMSO-$d_6$) δ 9.06 – 9.00 (m, 2H), 8.06 (dd, *J* = 8.4, 1.4 Hz, 1H), 8.00 – 7.91 (m, 4H), 7.83 (dd, *J* = 8.5, 7.3 Hz, 1H), 7.58 (dd, *J* = 7.3, 1.4 Hz, 1H), 4.15 – 3.99 (m, 2H), 3.39 (s, 2H), 3.31 (dd, *J* = 18.2, 5.2 Hz, 1H). **HRMS** (ESI): exact mass calculated for $C_{20}H_{14}ClN_5O$ [(M+H)$^+$] 376.096, found 376.0977.

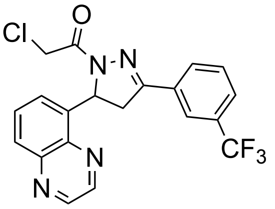

LEB-02-176

**2-chloro-1-(5-(quinoxalin-5-yl)-3-(3-(trifluoromethyl)phenyl)-4,5-dihydro-1H-pyrazol-1-yl)ethan-1-one (LEB-02-176).** General Procedure A was followed starting from 3-(trifluoromethyl)acetophenone (1.2 equiv, 0.76 mmol) and quinoxaline-5-carbaldehyde (1.0 equiv, 0.63 mmol). The corresponding chalcone precipitated out during the reaction and was filtered by gravity filtration to yield product (0.57 mmol) as a yellow solid. General Procedure F was followed starting from the above product (1.0 equiv, 0.57 mmol). Purification by flash column chromatography (EtOAc/hexane 50:50) yielded the chloroacetamide **LEB-02-176** (0.22 mmol, 35.3% yield across all steps). **$^1$H NMR** (400 MHz, DMSO-$d_6$) δ 9.06 – 9.03 (m, 2H), 8.12 – 8.04 (m, 3H), 7.84 (dt, *J* = 8.5, 6.7 Hz, 2H), 7.71 (t, *J* = 7.8 Hz, 1H), 7.57 (dd, *J* = 7.2, 1.4 Hz, 1H), 6.58 (dd, *J* = 12.0, 5.0 Hz, 1H), 4.97 (d, *J* = 14.1 Hz, 1H), 4.84 (d, *J* = 14.1 Hz, 1H), 4.12 (dd, *J* = 18.3, 12.1 Hz, 1H), 3.37 – 3.34 (m, 1H). **$^{13}$C NMR** (151 MHz, DMSO) δ 164.1, 155.7, 146.3, 145.3, 143.1, 139.7, 139.3, 132.3, 131.3, 130.5, 130.5, 130.4, 130.2, 130.0, 129.8, 129.0, 127.5, 127.4, 127.4, 127.1, 125.9, 125.3, 123.8, 123.8, 123.7, 123.5, 121.7, 57.0, 43.0, 42.1. **HRMS** (ESI): exact mass calculated for $C_{20}H_{14}ClF_3N_4O$ [(M+H)$^+$] 419.0881, found 419.0857.

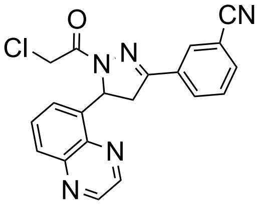

LEB-02-182

**3-(1-(2-chloroacetyl)-5-(quinoxalin-5-yl)-4,5-dihydro-1H-pyrazol-3-yl)benzonitrile (LEB-02-182).** General Procedure A was followed starting from 3-acetylbenzonitrile (1.2 equiv, 0.76 mmol) and quinoxaline-5-carbaldehyde (1.0 equiv, 0.63 mmol). The corresponding chalcone precipitated out during the reaction and was filtered by gravity filtration to yield product (0.46 mmol) as a light yellow solid. General Procedure F was followed starting from the above product (1.0 equiv, 0.46 mmol). Purification by flash column chromatography (EtOAc/hexane 50:50) yielded the chloroacetamide **LEB-02-182** (0.025 mmol, 4.0% yield across all steps). **$^1$H NMR** (400 MHz, DMSO-$d_6$) δ 9.09 – 9.00 (m, 2H), 8.28 – 8.23 (m, 1H), 8.14 (dt, *J* = 8.0, 1.4 Hz, 1H), 8.07 (dd, *J* = 8.4, 1.4 Hz, 1H), 7.96 (dt, *J* = 7.8, 1.4 Hz, 1H), 7.83 (dd, *J* = 8.4, 7.2 Hz, 1H), 7.68 (t, *J* = 7.9 Hz, 1H), 7.58 – 7.55 (m, 1H), 6.57 (dd, *J* = 11.9, 5.0 Hz, 1H), 4.96 (d, *J* = 14.1 Hz, 1H), 4.83 (d, *J* = 14.1 Hz, 1H), 4.14 – 4.01 (m, 1H), 3.34 – 3.27 (m, 1H). **HRMS** (ESI): exact mass calculated for $C_{20}H_{14}ClN_5O$ [(M+H)$^+$] 376.096, found 376.097.

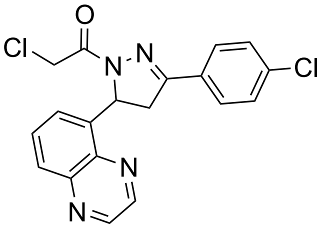

LEB-02-187

**2-chloro-1-(3-(4-chlorophenyl)-5-(quinoxalin-5-yl)-4,5-dihydro-1*H*-pyrazol-1-yl)ethan-1-one (LEB-02-187).** General Procedure A was followed starting from 4-chloroacetophenone (1.2 equiv, 0.76 mmol) and quinoxaline-5-carbaldehyde (1.0 equiv, 0.63 mmol). The corresponding chalcone precipitated out during the reaction and was filtered by gravity filtration to yield product (0.63 mmol) as a light yellow solid. General Procedure F was followed starting from the above crude product (1.0 equiv, 0.63 mmol). Purification by flash column chromatography (EtOAc/hexane 50:50) yielded the chloroacetamide **LEB-02-187** (0.12 mmol, 19.2% yield across all steps). **$^1$H NMR** (400 MHz, DMSO-$d_6$) δ 9.08 – 8.98 (m, 2H), 8.06 (dd, *J* = 8.4, 1.4 Hz, 1H), 7.87 – 7.77 (m, 3H), 7.58 – 7.49 (m, 3H), 6.55 (dd, *J* = 11.9, 5.0 Hz, 1H), 4.91 (d, *J* = 13.9 Hz, 1H), 4.78 (d, *J* = 13.9 Hz, 1H), 4.07 (dd, *J* = 18.2, 12.0 Hz, 1H), 3.26 (dd, *J* = 18.1, 5.0 Hz, 1H). **HRMS** (ESI): exact mass calculated for $C_{19}H_{14}Cl_2N_4O$ [(M+H)$^+$] 385.0617, found 385.0628.

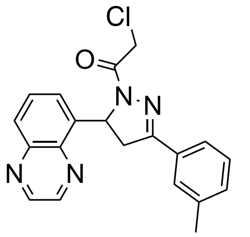

LEB-03-001c

**2-chloro-1-(5-(quinoxalin-5-yl)-3-(*m*-tolyl)-4,5-dihydro-1*H*-pyrazol-1-yl)ethan-1-one (LEB-03-001c).** General Procedure A was followed starting from 3-methylacetophenone (1.2 equiv, 0.76 mmol) and quinoxaline-5-carbaldehyde (1.0 equiv, 0.63 mmol). The corresponding chalcone precipitated out during the reaction and was filtered by gravity filtration to yield product (0.53 mmol) as a yellow solid. General Procedure F was followed starting from the above crude product (1.0 equiv, 0.53 mmol). Purification by flash column chromatography (EtOAc/hexane 50:50) yielded the chloroacetamide **LEB-03-001c** (0.29 mmol, 46.7% yield across all steps). **$^1$H NMR** (400 MHz, DMSO-$d_6$) δ 9.28 – 8.93 (m, 2H), 8.17 – 7.87 (m, 2H), 7.85 – 7.79 (m, 1H), 7.69 – 7.49 (m, 1H), 7.40 – 7.13 (m, 2H), 5.01 – 4.71 (m, 2H), 3.27 – 3.04 (m, 1H), 2.51 (s, 3H), 2.35 (d, *J* = 14.1 Hz, 3H). **HRMS** (ESI): exact mass calculated for $C_{20}H_{17}ClN_4O$ [(M+H)$^+$] 365.1164, found 365.1185.

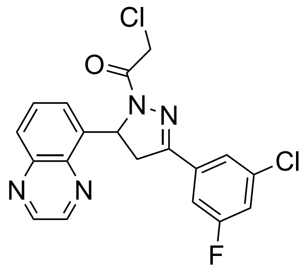

LEB-03-004c

**2-chloro-1-(3-(4-fluoro-2-methoxyphenyl)-5-(quinoxalin-5-yl)-4,5-dihydro-1*H*-pyrazol-1-yl)ethan-1-one (LEB-03-004c).** General Procedure A was followed starting from 3-chloro-5-fluoroacetophenone (1.2 equiv, 0.76 mmol) and quinoxaline-5-carbaldehyde (1.0 equiv, 0.63 mmol). The corresponding chalcone precipitated out during the reaction and was filtered by gravity filtration to yield product (0.51 mmol) as a light yellow solid. General Procedure F was followed starting from the above crude product (1.0 equiv, 0.51 mmol). Purification by flash column chromatography (EtOAc/hexane 50:50) yielded the chloroacetamide **LEB-03-004c** (0.23 mmol, 37.2% yield across all steps). $^{1}$**H NMR** (400 MHz, DMSO-$d_6$) δ 9.04 (dd, *J* = 4.3, 1.8 Hz, 2H), 8.08 – 7.91 (m, 2H), 7.85 (dt, *J* = 15.5, 7.9 Hz, 1H), 7.54 (d, *J* = 7.2 Hz, 1H), 6.89 (qd, *J* = 8.7, 2.4 Hz, 1H), 6.50 (dd, *J* = 11.9, 4.8 Hz, 1H), 4.91 – 4.71 (m, 1H), 4.15 – 4.00 (m, 1H), 3.72 (s, 2H), 3.18 (dd, *J* = 18.6, 4.9 Hz, 1H), 2.51 (dt, *J* = 3.7, 1.9 Hz, 3H). $^{13}$**C NMR** (151 MHz, DMSO) δ 164.2, 163.5, 161.8, 154.8, 154.8, 146.3, 145.3, 143.1, 139.7, 139.2, 135.1, 135.0, 134.9, 134.8, 130.5, 129.1, 126.0, 123.5, 123.5, 118.4, 118.2, 113.3, 113.1, 57.2, 51.8, 43.0, 42.1, 25.0, 16.7. **HRMS** (ESI): exact mass calculated for $C_{20}H_{16}ClFN_4O$ [(M+H)$^+$] 403.0523, found 403.0546.

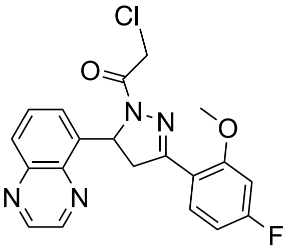

LEB-03-005c

**2-chloro-1-(3-(4-fluoro-2-methoxyphenyl)-5-(quinoxalin-5-yl)-4,5-dihydro-1*H*-pyrazol-1-yl)ethan-1-one (LEB-03-005c).** General Procedure A was followed starting from 4-fluoro-2-methoxyacetophenone (1.2 equiv, 0.76 mmol) and quinoxaline-5-carbaldehyde (1.0 equiv, 0.63 mmol). The corresponding chalcone precipitated out during the reaction and was filtered by gravity filtration to yield product (0.51 mmol) as a white solid. General Procedure F was followed starting from the above crude product (1.0 equiv, 0.51 mmol). Purification by flash column chromatography (EtOAc/hexane 50:50) yielded the chloroacetamide **LEB-03-005c** (0.13 mmol, 20.0% yield across all steps). $^{1}$**H NMR** (400 MHz, DMSO-$d_6$) δ 9.04 (dd, *J* = 4.3, 1.8 Hz, 2H), 8.08 – 7.91 (m, 2H), 7.85 (dt, *J* = 15.5, 7.9 Hz, 1H), 7.54 (d, *J* = 7.2 Hz, 1H), 6.89 (qd, *J* = 8.7, 2.4 Hz, 1H), 6.50 (dd, *J* = 11.9, 4.8 Hz, 1H), 4.91 – 4.71 (m, 1H), 4.15 – 4.00 (m, 1H), 3.72 (s, 2H), 3.18 (dd, *J* = 18.6, 4.9 Hz, 1H), 2.51 (dt, *J* = 3.7, 1.9 Hz, 3H). $^{13}$**C NMR** (151 MHz, DMSO) δ 165.8, 164.1, 163.7, 160.3, 160.2, 155.1, 146.3, 145.3, 143.1, 139.6, 131.2, 131.1, 130.6, 128.9, 125.7, 116.7, 116.7, 108.1, 108.0, 101.1, 101.0, 56.8, 56.3, 45.3, 43.0. **HRMS** (ESI): exact mass calculated for $C_{20}H_{16}ClFN_4O$ [(M+H)$^+$] 399.1019, found 399.1042

.

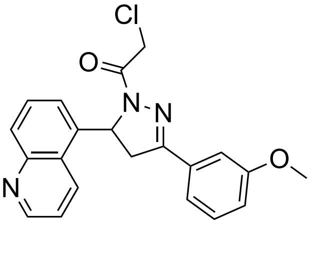

**2-chloro-1-(3-(3-methoxyphenyl)-5-(quinolin-5-yl)-4,5-dihydro-1H-pyrazol-1-yl)ethan-1-one (LEB-03-008c).** General Procedure A was followed starting from 3-methoxyacetophenone (1.2 equiv, 1.14 mmol) and quinoline-5-carbaldehyde (1.0 equiv, 0.95 mmol). The reaction was run for 15 minutes at 0°C. The corresponding chalcone was purified via flash chromatography (EtOAc/Hexane 50:50) to yield a light yellow solid (0.69 mmol). General Procedure F was followed starting from the above product (1.0 equiv, 0.69 mmol). Purification by flash column chromatography (EtOAc/hexane 50:50) yielded the chloroacetamide **LEB-03-008c** (0.29 mmol, 30.3% yield across all steps). **$^{1}$H NMR** (400 MHz, DMSO-d6) δ 8.99 (dd, J = 4.2, 1.6 Hz, 1H), 8.67 (d, J = 8.6 Hz, 1H), 7.98 (d, J = 8.5 Hz, 1H), 7.75 – 7.61 (m, 2H), 7.44 – 7.27 (m, 4H), 4.89 (d, J = 30.0 Hz, 1H), 4.11 (dd, J = 18.2, 11.9 Hz, 1H), 3.79 (s, 3H), 3.36 (s, 2H), 3.26 (d, J = 4.9 Hz, 1H). **$^{13}$C NMR** (151 MHz, DMSO) δ 167.9, 164.3, 164.0, 162.4, 160.0, 159.9, 157.8, 156.5, 156.4, 151.5, 151.2, 150.8, 148.6, 148.6, 148.0, 144.8, 143.9, 140.8, 140.7, 139.7, 139.6, 137.9, 137.6, 137.5, 135.9, 133.3, 132.5, 132.4, 132.2, 132.1, 132.0, 131.7, 130.6, 130.4, 129.7, 129.6, 129.3, 129.1, 128.7, 127.0, 125.8, 125.7, 125.1, 122.8, 122.6, 122.2, 122.0, 120.1, 119.9, 117.4, 117.2, 112.8, 112.6, 112.4, 59.9, 57.4, 56.9, 55.9, 55.8, 55.5, 43.0, 42.7, 42.6, 42.5, 42.3, 24.8. **HRMS** (ESI): exact mass calculated for $C_{21}H_{18}ClN_3O_2$ [(M+H)$^+$] 380.116, found 380.1185.

**1-(3-Phenyl-5-(quinoxalin-5-yl)-4,5-dihydro-1*H*-pyrazol-1-yl)but-2-yn-1-one (CMZ 07)**

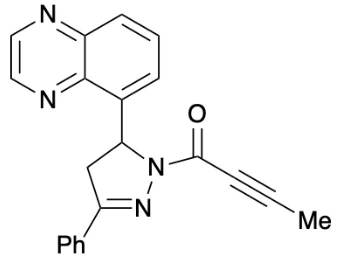


*Step 1*: Following General Procedure A, the chalcone was prepared from quinoxaline-5-carbaldehyde (500 mg, 3.16 mmol), acetophenone (443 μL, 3.79 mmol) and 5% (w/w) aq NaOH (3.8 mL, 4.74 mmol) in EtOH (10 mL). Filtration under reduced pressure gave the chalcone as a pale yellow solid (823 mg, quant) which was used without purification.
*Step 2*: Hydrazine hydrate (136 μL, 1.54 mmol, 50–60% wt solution) was added to a solution of the chalcone (200 mg, 0.77 mmol) in EtOH (2 mL) at rt and the resultant mixture was heated at reflux for 4 h before being cooled to rt. One half of the reaction mixture was then concentrated *in vacuo* and the crude pyrazoline was used immediately without purification.
*Step 3*: A solution of tetrolic acid (32 mg, 0.38 mmol) in DMF (0.5 mL) was added to a solution of the crude pyrazoline (0.38 mmol) in DMF (0.5 mL) at 0 °C. The reaction mixture was then treated sequentially with NMM (422 μL, 3.84 mmol) and T3P (642 μL, 1.08 mmol, 50 wt% in EtOAc) and allowed to warm to rt and stirred at rt for 16 h. Water (2 mL) and EtOAc (2 mL) were then added and the aqueous layer was extracted with EtOAc (3 × 2 mL). The combined organic extracts were washed sequentially with satd aq $NaHCO_3$ (6 mL) and NaCl solution (sat. aqueous) (6 mL), then dried and concentrated *in vacuo*. Purification *via* flash column chromatography (**Sfär Silica HC D**, 0% grading to 100% EtOAc in hexane, product eluted at 55%) gave **CMZ 7** as a white solid (47 mg, 36% over 2 steps from the respective chalcone); **$^{1}$H NMR** (400 MHz, $CDCl_3$) $\delta_H$ 2.08 (s, 3H), 3.11 (dd, *J* = 17.8, 4.8 Hz, 1H), 3.98 (dd, *J* = 17.8, 11.8 Hz, 1H), 6.62 (dd, *J* = 11.8, 4.8 Hz, 1H), 7.25 – 7.40 (m, 3H), 7.46 (d, *J* = 7.2 Hz, 1H), 7.59 – 7.75 (m, 3H), 7.92 – 7.99 (m, 1H), 8.77 – 8.83 (m, 2H). **$^{13}$C NMR** (151 MHz, $CDCl_3$) $\delta_C$ 4.5, 42.5, 56.7, 73.9, 90.3, 125.6, 126.9, 128.7, 129.0, 130.1, 130.7, 131.1, 138.3, 140.2, 143.2, 143.9, 144.7, 151.3, 156.8. **HRMS (ESI):** exact mass calculated for $C_{21}H_{16}N_4O$ [(M+H)$^+$] 341.1397, found 341.1376.

**2-Chloro-1-(3-phenyl-5-(quinolin-5-yl)-4,5-dihydro-1*H*-pyrazol-1-yl)ethan-1-one (CMZ 13)**

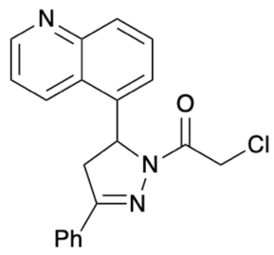


*Step 1*: Following General Procedure A, the chalcone was prepared from quinoline-5-carbaldehyde (55 mg, 0.35 mmol), acetophenone (49 μL, 0.42 mmol) and 5% (w/w) aq NaOH (0.4 mL, 0.52 mmol) in EtOH (1 mL). Filtration under reduced pressure gave the chalcone as a brown solid (58 mg, 65%).
*Step 2*: Following General Procedure F, the pure title compound was prepared from chalcone (58 mg, 0.22 mmol) and hydrazine hydrate (40 μL, 0.45 mmol, 50–60% wt solution) in EtOH (0.9 mL); then chloroacetyl chloride (27 μL, 0.34 mmol) and triethylamine (94 μL, 0.67 mmol) in DCM (0.9 mL). Purification *via* flash column chromatography (**Sfär Silica HC D**, 0% grading to 100% EtOAc in hexane, product eluted at 61%) gave **CMZ 13** as a white solid (35 mg, 44% over 2 steps from the respective chalcone). **$^1$H NMR** (400 MHz, $CDCl_3$) $\delta_H$ 3.13 (dd, *J* = 17.6, 4.9 Hz, 1H), 3.90 (dd, *J* = 17.6, 11.9 Hz, 1H), 4.55 (d, *J* = 13.4 Hz, 1H), 4.65 (d, *J* = 13.4 Hz, 1H), 6.21 (dd, *J* = 11.9, 4.9 Hz, 1H), 7.27 (d, *J* = 7.2 Hz, 1H), 7.31 – 7.44 (m, 4H), 7.57 (dd, *J* = 8.6, 7.2 Hz, 1H), 7.63 – 7.71 (m, 2H), 7.98 (d, *J* = 8.6 Hz, 1H), 8.29 (d, *J* = 8.6 Hz, 1H), 8.89 (dd, *J* = 4.3, 1.6 Hz, 1H). **$^{13}$C NMR** (151 MHz, $CDCl_3$) $\delta_C$ 42.1, 42.4, 57.1, 121.4, 123.3, 125.2, 127.0, 129.0, 129.2, 130.0, 130.6, 131.2, 132.3, 136.6, 147.9, 149.6, 155.9, 164.5. **HRMS (ESI):** exact mass calculated for $C_{20}H_{16}ClN_3O$ [$(M+H)^+$] 350.1055, found 350.1029.

**2-Chloro-1-(3-phenyl-5-(quinoxalin-6-yl)-4,5-dihydro-1*H*-pyrazol-1-yl)ethan-1-one (CMZ 15)**

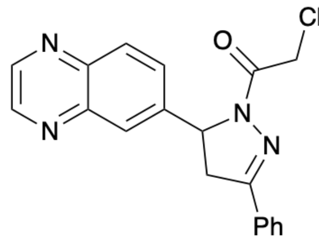


*Step 1*: Following General Procedure A, the chalcone was prepared from quinoxaline-6-carbaldehyde (301 mg, 1.90 mmol), acetophenone (267 μL, 2.28 mmol) and 5% (w/w) aq NaOH (2.28 mL, 2.85 mmol) in EtOH (9 mL). Filtration under reduced pressure gave the chalcone as a pale brown solid (445 mg, 90%).
*Step 2*: Following General Procedure F, the pure title compound was prepared from chalcone (100 mg, 0.38 mmol) and hydrazine hydrate (68 μL, 0.77 mmol, 50–60% wt solution) in EtOH (1.6 mL); then chloroacetyl chloride (46 μL, 0.58 mmol) and triethylamine (161 μL, 1.15 mmol) in DCM (1.5 mL). Purification *via* flash column chromatography (**Sfär Silica HC D**, 0% grading to 100% EtOAc in hexane, product eluted at 59%) gave **CMZ 15** as a white solid (50 mg, 37% over 2 steps from the respective chalcone); **$^1$H NMR** (400 MHz, $CDCl_3$) $\delta_H$ 3.23 (dd, *J* = 17.9, 4.9 Hz, 1H), 3.84 (dd, *J* = 17.9, 11.8 Hz, 1H), 4.49 – 4.61 (m, 2H), 5.74 (dd, *J* = 11.8, 4.9 Hz, 1H), 7.32 – 7.44 (m, 3H), 7.60 (dd, *J* = 8.7, 2.1 Hz, 1H), 7.61 – 7.72 (m, 2H), 7.91 (d, *J* = 2.1 Hz, 1H), 8.02 (d, *J* = 8.7 Hz, 1H), 8.73 (app s, 2H). **$^{13}$C NMR** (151 MHz, $CDCl_3$) $\delta_C$ 42.1, 42.1, 60.2, 126.0, 126.8, 128.0, 128.8, 130.4, 130.5, 131.0, 142.4, 142.8, 142.9, 145.0, 145.2, 155.3, 164.2. **HRMS (ESI):** exact mass calculated for $C_{19}H_{15}ClN_4O$ [$(M+MeCN+H)^+$] 392.1272, found 392.1243.

**2-(3-Phenyl-5-(35uinoxaline-6-yl)-4,5-dihydro-1*H*-pyrazol-1-yl)ethane-1-sulfonyl fluoride (CMZ 17)**

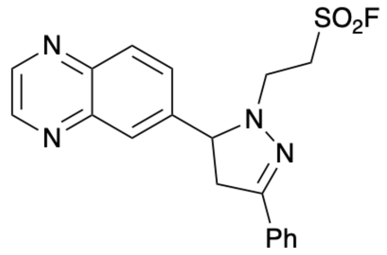

*Step 1*: Following General Procedure A, the chalcone was prepared from quinoxaline-6-carbaldehyde (301 mg, 1.90 mmol), acetophenone (267 μL, 2.28 mmol) and 5% (w/w) aq NaOH (2.3 mL, 2.85 mmol) in EtOH (9 mL). Filtration under reduced pressure gave the chalcone as a pale brown solid (495 mg, 90%) which was used without purification.
*Step 2*: Hydrazine hydrate (48 μL, 0.54 mmol, 50–60% wt solution) was added to a solution of the chalcone (71 mg, 0.27 mmol) in EtOH (1 mL) at rt and the resultant mixture was heated at reflux for 1.5 h before being cooled to rt. One third of the reaction mixture was then concentrated *in vacuo* and the crude pyrazoline was used immediately without purification.
*Step 3*: Ethene sulfonyl fluoride (25 μL, 0.30 mmol) was added to a solution of the crude pyrazoline (0.27 mmol) in THF (0.6 mL) at rt and stirred at 80°C for 4 h, then concentrated *in vacuo*. Purification *via* flash column chromatography (**Sfär Silica HC D**, 0% grading to 100% EtOAc in hexane, product eluted at 43%) gave **CMZ 17** as a pale yellow oil (63 mg, 60% over 2 steps from the respective chalcone); **$^{1}$H NMR** (400 MHz, $CDCl_3$) $\delta_H$ 3.06 (dd, *J* = 16.3, 13.8 Hz, 1H), 3.22 – 3.33 (m, 1H), 3.39 (app dt, *J* = 12.6, 7.7 Hz, 1H), 3.54 (dd, *J* = 16.3, 10.0 Hz, 1H), 3.84 – 3.99 (m, 2H), 4.45 (dd, *J* = 13.8, 10.0 Hz, 1H), 7.26 – 7.42 (m, 3H), 7.55 – 7.63 (m, 2H), 7.89 (dd, *J* = 8.7, 2.0 Hz, 1H), 8.05 – 8.13 (m, 2H), 8.80 (app s, 2H). **$^{13}$C NMR** (151 MHz, $CDCl_3$) $\delta_C$ 43.2, 47.7, 49.5, 49.6, 71.2, 126.1, 128.2, 128.6, 129.0, 129.5, 130.3, 131.8, 141.6, 142.7, 142.9, 145.1, 145.2, 151.6. **HRMS (ESI)**: exact mass calculated for $C_{19}H_{17}FN_4O_2S$ [$(M+H)^+$] 385.1129, found 385.1111.

**3-Oxo-3-(3-phenyl-5-(quinoxaline-5-yl)-4,5-dihydro-1*H*-pyrazol-1-yl)propanenitrile (CMZ 22)**

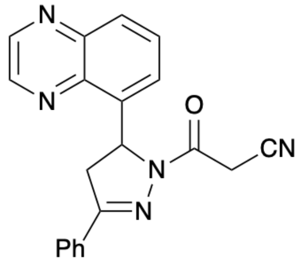

*Step 1*: Following General Procedure A, the chalcone was prepared from quinoxaline-5-carbaldehyde (500 mg, 3.16 mmol), acetophenone (443 μL, 3.79 mmol) and 5% (w/w) aq NaOH (3.8 mL, 4.74 mmol) in EtOH (10 mL). Filtration under reduced pressure gave the chalcone as a pale yellow solid (823 mg, quant) which was used without purification.
*Step 2*: Hydrazine hydrate (68 μL, 0.77 mmol, 50–60% wt solution) was added to a solution of the chalcone (100 mg, 0.38 mmol) in EtOH (2 mL) at rt and the resultant mixture was heated at reflux for 2 h before being cooled to rt. The reaction mixture was then concentrated *in vacuo* and the crude pyrazoline was used immediately without purification.
*Step 3*: A solution of cyanoacetic acid (33 mg, 0.38 mmol) in DMF (0.5 mL) was added to a solution of the crude pyrazoline (0.38 mmol) in DMF (0.5 mL) at 0 °C. The reaction mixture was then treated sequentially with NMM (422 μL, 3.84 mmol) and T3P (643 μL, 1.08 mmol, 50 wt% in EtOAc) and allowed to warm to rt and stirred at rt for 16 h. Water (2 mL) and EtOAc (2 mL) were then added and the aqueous layer was extracted with EtOAc (3 × 2 mL). The combined organic extracts were washed sequentially with satd aq $NaHCO_3$ (6 mL) and NaCl solution (sat. aqueous) (6 mL), then dried and concentrated *in vacuo*. Purification *via* flash column chromatography (**Sfär Silica HC D**, 0% grading to 100% EtOAc in hexane, product eluted at 48%) gave **CMZ 22** as a white solid (11 mg, 8% over 2 steps from the respective chalcone); **$^{1}$H NMR** (600 MHz, DMSO-*d*$_6$) $\delta_H$ 3.27 (dd, *J* = 18.2, 5.0 Hz, 1H), 4.10 (dd, *J* = 18.2, 11.9 Hz, 1H), 4.36 (d, *J* = 19.2 Hz, 1H), 4.48 (d, *J* = 19.1 Hz, 1H), 6.55 (dd, *J* = 11.9, 5.0 Hz, 1H), 7.43 – 7.53 (m, 3H), 7.59 (dd, *J* = 7.4, 1.3 Hz, 1H), 7.78 – 7.82 (m, 2H), 7.82 – 7.86 (m, 1H), 8.06 (dd, *J* = 8.4, 1.3 Hz, 1H), 9.04 (dd, *J* = 10.5, 1.8 Hz, 2H). **$^{13}$C NMR** (151 MHz, $CDCl_3$) $\delta_C$ 25.5, 40.1, 42.3, 56.3, 116.0, 125.5, 127.0, 128.7, 128.9, 130.1, 130.7, 130.9, 138.9, 139.3, 142.7, 144.9, 145.9, 156.7, 160.5. **HRMS (ESI)**: exact mass calculated for $C_{20}H_{15}N_5O$ [$(M+H)^+$] 342.135, found 342.1336.

**5-(3-Phenyl-1-(vinylsulfonyl)-4,5-dihydro-1*H*-pyrazol-5-yl)quinoxaline (CMZ 23)**

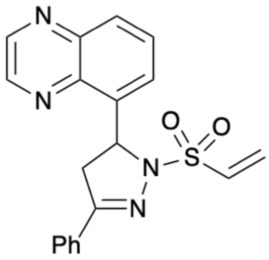

*Step 1*: Following General Procedure A, the chalcone was prepared from quinoxaline-5-carbaldehyde (500 mg, 3.16 mmol), acetophenone (443 μL, 3.79 mmol) and 5% (w/w) aq NaOH (3.8 mL, 4.74 mmol) in EtOH (10 mL). Filtration under reduced pressure gave the chalcone as a pale yellow solid (823 mg, quant) which was used without purification.
*Step 2*: Hydrazine hydrate (204 μL, 2.31 mmol, 50–60% wt solution) was added to a solution of the chalcone (300 mg, 1.15 mmol) in EtOH (4.5 mL) at rt and the resultant mixture was heated at reflux for 2 h before being cooled to rt. The reaction mixture was then concentrated *in vacuo* and the crude pyrazoline was used immediately without purification.
*Step 3*: Following General Procedure G, the pure title compound was prepared from crude pyrazoline (1.15 mmol), triphenylphosphine oxide (712 mg, 2.56 mmol), trifluoromethanesulfonic anhydride (194 μL, 1.15 mmol), vinyl sulfonic acid (125 mg, 1.15 mmol), pyridine (93 μL, 1.15 mmol) and triethylamine (321 μL, 2.31 mmol) in DCM (17 mL). Purification *via* flash column chromatography (**Sfär Silica HC D**, 0% grading to 100% EtOAc in hexane, product eluted at 53%; then SNAP Ultra C18, 20% grading to 100% MeCN in water, product eluted at 43%) gave **CMZ 23** as a white solid (33 mg, 24% over 2 steps from the respective chalcone); **$^1$H NMR** (400 MHz, DMSO-$d_6$) $\delta_H$ 3.29 (dd, *J* = 17.9, 9.0 Hz, 1H), 4.14 (dd, *J* = 17.9, 11.9 Hz, 1H), 6.18 (dd, *J* = 11.9, 9.0 Hz, 1H), 6.26 – 6.40 (m, 2H), 7.02 (dd, *J* = 16.5, 10.0 Hz, 1H), 7.42 – 7.54 (m, 3H), 7.73 – 7.79 (m, 2H), 7.90 – 7.97 (m, 1H), 7.98 – 8.02 (m, 1H), 8.10 (dd, *J* = 8.4, 1.7 Hz, 1H), 8.95 – 9.08 (m, 2H). **HRMS (ESI)**: exact mass calculated for $C_{19}H_{16}N_4O_2S$ [(M+H)$^+$] 365.1067, found 365.1061.

**3-Phenyl-5-(quinoxalin-5-yl)-4,5-dihydro-1*H*-pyrazole-1-sulfonyl fluoride (CMZ 24)**

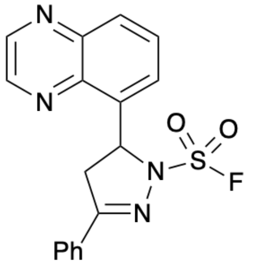

*Step 1*: Following General Procedure A, the chalcone was prepared from quinoxaline-5-carbaldehyde (500 mg, 3.16 mmol), acetophenone (443 μL, 3.79 mmol) and 5% (w/w) aq NaOH (3.8 mL, 4.74 mmol) in EtOH (10 mL). Filtration under reduced pressure gave the chalcone as a pale yellow solid (823 mg, quant) which was used without purification.
*Step 2*: Hydrazine hydrate (204 μL, 2.31 mmol, 50–60% wt solution) was added to a solution of the chalcone (300 mg, 1.15 mmol) in EtOH (4.5 mL) at rt and the resultant mixture was heated at reflux for 4 h before being cooled to rt. One third of the reaction mixture was then concentrated *in vacuo* and the crude pyrazoline was used immediately without purification.
*Step 3*: 1-(Fluorosulfuryl)-2,3-dimethyl-1H-imidazol-3-ium trifluoromethanesulfonate (126 mg, 0.38 mmol) was added to a solution of the crude pyrazoline (0.38 mmol) in MeCN (0.7 mL) at rt and stirred at this temperature for 16 h. Water (2 mL) was then added and the aqueous layer was extracted with EtOAc (3 × 2 mL). The combined organic extracts were washed sequentially with NaCl solution (sat. aqueous) (2 × 2 mL), then dried and concentrated *in vacuo*. Purification *via* flash column chromatography (**Sfär Silica HC D**, 0% grading to 100% EtOAc in hexane, product eluted at 25%) gave **CMZ 24** as a white solid (34 mg, 25% over 2 steps from the respective chalcone); **$^1$H NMR** (400 MHz, $CDCl_3$) $\delta_H$ 3.24 (dd, *J* = 17.7, 8.2 Hz, 1H), 4.18 (dd, *J* = 17.7, 11.4 Hz, 1H), 6.50 (dd, *J* = 11.4, 8.2 Hz, 1H), 7.30 – 7.46 (m, 3H), 7.68 – 7.74 (m, 2H), 7.74 – 7.78 (m, 1H), 7.84 – 7.92 (m, 1H), 8.04 (dd, *J* = 8.5, 1.6 Hz, 1H), 8.73 – 8.88 (m, 2H). **$^{13}$C NMR** (151 MHz, $CDCl_3$) $\delta_C$ 44.3, 61.4, 127.2, 127.6, 129.0, 129.8, 130.0, 130.3, 131.7, 138.0, 140.3, 143.2, 144.3, 145.4, 160.2. **HRMS (ESI)**: exact mass calculated for $C_{17}H_{13}FN_4O_2S$ [(M+MeCN+H)$^+$] 398.1081, found 398.1097.

**1-(3-Phenyl-5-(quinoxalin-5-yl)-4,5-dihydro-1*H*-pyrazole-1-carbonyl)cyclopropane-1-carbonitrile (CMZ 29)**

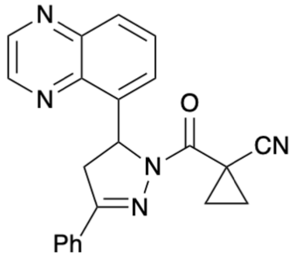

*Step 1*: Following General Procedure A, the chalcone was prepared from quinoxaline-5-carbaldehyde (500 mg, 3.16 mmol), acetophenone (443 μL, 3.79 mmol) and 5% (w/w) aq NaOH (3.8 mL, 4.74 mmol) in EtOH (10 mL). Filtration under reduced pressure gave the chalcone as a pale yellow solid (823 mg, quant) which was used without purification.
*Step 2*: Hydrazine hydrate (204 μL, 2.31 mmol, 50–60% wt solution) was added to a solution of the chalcone (300 mg, 1.15 mmol) in EtOH (4.5 mL) at rt and the resultant mixture was heated at reflux for 4 h before being cooled to rt. One third of the reaction mixture was then concentrated *in vacuo* and the crude pyrazoline was used immediately without purification.
*Step 3*: A solution of 1-cyano-1-cyclopropane carboxylic acid (85 mg, 0.77 mmol) in DMF (1 mL) was added to a solution of the crude pyrazoline (0.77 mmol) in DMF (1 mL) at 0 °C. The reaction mixture was then treated sequentially with NMM (845 μL, 7.68 mmol) and T3P (1.3 mL, 2.15 mmol, 50 wt% in EtOAc) and allowed to warm to rt and stirred at rt for 16 h. Water (5 mL) and EtOAc (5 mL) were then added and the aqueous layer was extracted with EtOAc (3 × 5 mL). The combined organic extracts were washed sequentially with satd aq $NaHCO_3$ (15 mL) and NaCl solution (sat. aqueous) (15 mL), then dried and concentrated *in vacuo*. Purification *via* flash column chromatography (**Sfär Silica HC D**, 0% grading to 100% EtOAc in hexane, product eluted at 39%) gave **CMZ 29** as a white solid (41 mg, 15% over 2 steps from the respective chalcone); **$^1$H NMR** (400 MHz, $CDCl_3$) $\delta_H$ 1.57 – 1.66 (m, 3H), 1.72 (ddd, *J* = 9.2, 4.7, 3.2 Hz, 1H), 3.13 (dd, *J* = 17.9, 5.4 Hz, 1H), 3.95 (dd, *J* = 17.9, 11.8 Hz, 1H), 6.63 (dd, *J* = 11.8, 5.4 Hz, 1H), 7.27 – 7.42 (m, 3H), 7.48 (d, *J* = 7.2 Hz, 1H), 7.67 (app t, *J* = 8.0 Hz, 1H), 7.74 – 7.87 (m, 2H), 7.93 – 8.04 (m, 1H), 8.60 – 8.95 (m, 2H). **$^{13}$C NMR** (151 MHz, $CDCl_3$) $\delta_C$ 14.2, 17.9, 18.6, 42.1, 57.4, 120.9, 125.7, 127.3, 129.0, 129.4, 130.3, 131.0, 131.1, 138.9, 140.2, 143.2, 144.3, 145.1, 156.4, 162.5. **HRMS (ESI)**: exact mass calculated for $C_{22}H_{17}N_5O$ [$(M+H)^+$] 368.1506, found 368.1524.

**2-Chloro-1-(3-phenyl-5-(quinolin-4-yl)-4,5-dihydro-1*H*-pyrazol-1-yl)ethan-1-one (CMZ 35)**

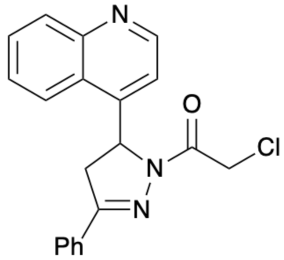

*Step 1*: Following General Procedure A, the chalcone was prepared from quinoline-4-carbaldehyde (137 mg, 0.87 mmol), acetophenone (204 μL, 1.74 mmol) and 5% (w/w) aq NaOH (0.3 mL, 0.38 mmol) in EtOH (5 mL). Extraction with EtOAc and purification *via* flash column chromatography (**Sfär Silica HC D**, 0% grading to 100% EtOAc in hexane, product eluted at 42%) gave the chalcone as a yellow solid (39 mg, 17%).
*Step 2*: Following General Procedure F, the pure title compound was prepared from chalcone (39 mg, 0.15 mmol) and hydrazine hydrate (27 μL, 0.30 mmol, 50–60% wt solution) in EtOH (1.5 mL); then chloroacetyl chloride (18 μL, 0.23 mmol) and triethylamine (63 μL, 0.45 mmol) in DCM (1.5 mL). Purification *via* flash column chromatography (**Sfär Silica HC D**, 0% grading to 100% EtOAc in hexane, product eluted at 46%) gave **CMZ 35** as a white solid (31 mg, 58% over 2 steps from the respective chalcone); **$^1$H NMR** (400 MHz, $CDCl_3$) $\delta_H$ 3.12 (dd, *J* = 17.6, 5.1 Hz, 1H), 3.95 (dd, *J* = 17.6, 12.0 Hz, 1H), 4.53 (d, *J* = 13.2 Hz, 1H), 4.69 (d, *J* = 13.2 Hz, 1H), 6.23 (dd, *J* = 12.0, 5.1 Hz, 1H), 7.09 (d, *J* = 4.5 Hz, 1H), 7.28 – 7.41 (m, 4H), 7.52 – 7.59

(m, 1H), 7.61 – 7.67 (m, 2H), 7.67 – 7.71 (m, 1H), 7.88 – 7.93 (m, 1H), 8.07 – 8.12 (m, 1H), 8.77 (d, *J* = 4.5 Hz, 1H). **$^{13}$C NMR** (151 MHz, $CDCl_3$) $\delta_C$ 41.9, 42.0, 57.1, 116.5, 122.9, 125.1, 127.0, 127.8, 129.1, 129.9, 130.3, 130.3, 131.4, 146.6, 147.5, 149.5, 156.0, 164.7. **HRMS (ESI)**: exact mass calculated for $C_{20}H_{16}ClN_3O$ [$(M+H)^+$] 350.1055, found 350.1066.

**2-Fluoro-1-(3-phenyl-5-(quinoxalin-5-yl)-4,5-dihydro-1*H*-pyrazol-1-yl)ethan-1-one (CMZ 38)**

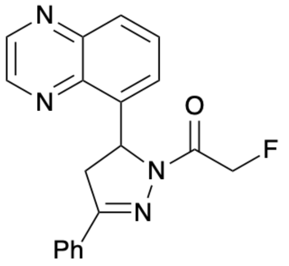


*Step 1*: Following General Procedure A, the chalcone was prepared from quinoxaline-5-carbaldehyde (500 mg, 3.16 mmol), acetophenone (443 μL, 3.79 mmol) and 5% (w/w) aq NaOH (3.8 mL, 4.74 mmol) in EtOH (10 mL). Filtration under reduced pressure gave the chalcone as a pale yellow solid (823 mg, quant) which was used without purification.
*Step 2*: Hydrazine hydrate (94 μL, 1.06 mmol, 50–60% wt solution) was added to a solution of the chalcone (138 mg, 0.53 mmol) in EtOH (2.2 mL) at rt and the resultant mixture was heated at reflux for 2 h before being cooled to rt. The reaction mixture was then concentrated *in vacuo* and the crude pyrazoline was used immediately without purification.
*Step 3*: A solution of fluoroacetic acid (41 mg, 0.53 mmol) in DMF (0.7 mL) was added to a solution of the crude pyrazoline (0.53 mmol) in DMF (0.7 mL) at 0 °C. The reaction mixture was then treated sequentially with NMM (583 μL, 5.30 mmol) and T3P (887 μL, 1.48 mmol, 50 wt% in EtOAc) and allowed to warm to rt and stirred at rt for 16 h. Water (2 mL) and EtOAc (2 mL) were then added and the aqueous layer was extracted with EtOAc (3 × 2 mL). The combined organic extracts were washed sequentially with satd aq $NaHCO_3$ (6 mL) and NaCl solution (sat. aqueous) (6 mL), then dried and concentrated *in vacuo*. Purification *via* flash column chromatography (**Sfär Silica HC D**, 0% grading to 100% EtOAc in hexane, product eluted at 46%) gave **CMZ 38** as a white solid (20 mg, 11% over 2 steps from the respective chalcone); **$^1$H NMR** (400 MHz, $CDCl_3$) $\delta_H$ 3.12 (dd, *J* = 17.9, 5.3 Hz, 1H), 3.94 (dd, *J* = 17.9, 11.9 Hz, 1H), 5.21 – 5.63 (m, 2H), 6.59 (dd, *J* = 11.9, 5.3 Hz, 1H), 7.28 – 7.41 (m, 3H), 7.42 – 7.50 (m, 1H), 7.58 – 7.69 (m, 3H), 7.98 (dd, *J* = 8.4, 1.4 Hz, 1H), 8.69 – 8.89 (m, 2H). **$^{13}$C NMR** (151 MHz, $CDCl_3$) $\delta_C$ 42.1, 56.9, 78.7, 79.8, 126.1, 126.9, 129.0, 129.2, 130.3, 130.9, 131.0, 138.6, 140.4, 143.1, 144.1, 144.7, 156.7, 165.2, 165.3. **HRMS (ESI):** exact mass calculated for $C_{19}H_{15}FN_4O$ [$(M+H)^+$] 335.1303, found 335.1304.

**2-Chloro-1-(5-(4-fluorophenyl)-3-phenyl-4,5-dihydro-1*H*-pyrazol-1-yl)ethan-1-one (CMZ 47)**

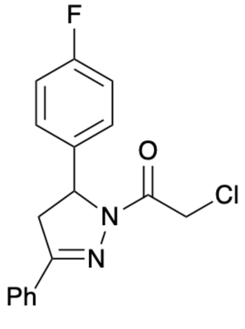


*Step 1*: Following General Procedure A, the chalcone was prepared from 4-fluorobenzaldehyde (259 μL, 2.42 mmol), acetophenone (339 μL, 2.90 mmol) and 5% (w/w) aq NaOH (2.9 mL, 3.63 mmol) in EtOH (3 mL). Filtration under reduced pressure gave the chalcone as a pale yellow solid (547 mg, quant).
*Step 2*: Following General Procedure F, the pure title compound was prepared from chalcone (100 mg, 0.44 mmol) and hydrazine hydrate (78 μL, 0.88 mmol, 50–60% wt solution) in EtOH (3 mL); then chloroacetyl chloride (53 μL, 0.66 mmol) and triethylamine (185 μL, 1.33 mmol) in DCM (1.5 mL). Purification *via* flash column chromatography (**Sfär Silica HC D**, 0% grading to 100% EtOAc in hexane, product eluted at 16%) gave **CMZ 47** as a white solid (44 mg, 31% over 2 steps from the respective chalcone); **$^1$H NMR** (400 MHz, $CDCl_3$) $\delta_H$ 3.15 (dd, *J* = 17.9, 4.7 Hz, 1H), 3.73 (dd, *J* = 17.9, 11.8 Hz, 1H), 4.44 – 4.57 (m, 2H), 5.51 (dd, *J* = 11.8, 4.7 Hz, 1H), 6.90 – 7.00 (m, 2H), 7.11 – 7.23 (m, 2H), 7.33 – 7.45 (m, 3H), 7.65 – 7.72 (m, 2H). **$^{13}$C**

**NMR** (151 MHz, $CDCl_3$) $\delta_C$ 42.3, 42.3, 60.0, 115.9, 116.1, 127.0, 127.0, 127.8, 129.0, 129.0, 130.8, 131.1, 136.8, 136.8, 155.5, 161.6, 163.3, 164.1. **HRMS (ESI):** exact mass calculated for $C_{17}H_{14}ClFN_2O$ $[(M+H)^+]$ 317.0852, found 317.0845.

**2-Chloro-1-(3-(2,5-dichlorophenyl)-5-(quinolin-5-yl)-4,5-dihydro-1*H*-pyrazol-1-yl)ethan-1-one (CMZ 51)**

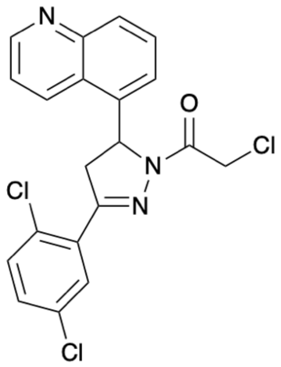


*Step 1*: Following General Procedure A, the chalcone was prepared from quinoline-5-carbaldehyde (200 mg, 1.27 mmol), 2′5′-dichloroacetophenone (367 μL, 2.55 mmol) and 5% (w/w) aq NaOH (438 μL, 0.55 mmol) in EtOH (4 mL). Filtration under reduced pressure and purification *via* flash column chromatography (**Sfär Silica HC D**, 0% grading to 100% EtOAc in hexane, product eluted at 32%) gave the chalcone as a yellow oil (168 mg, 40%).
*Step 2*: Following General Procedure F, the pure title compound was prepared from chalcone (168 mg, 0.51 mmol) and hydrazine hydrate (91 μL, 1.03 mmol, 50–60% wt solution) in EtOH (4 mL); then chloroacetyl chloride (61 μL, 0.77 mmol) and triethylamine (214 μL, 1.54 mmol) in DCM (1.5 mL). Purification *via* flash column chromatography (**Sfär Silica HC D**, 0% grading to 100% EtOAc in hexane, product eluted at 45%) gave **CMZ 51** as a white solid (53 mg, 25% over 2 steps from the respective chalcone); $^1$**H NMR** (400 MHz, $CDCl_3$) $\delta_H$ 3.28 (dd, *J* = 18.1, 4.9 Hz, 1H), 4.10 (dd, *J* = 18.1, 12.0 Hz, 1H), 4.51 (d, *J* = 13.5 Hz, 1H), 4.62 (d, *J* = 13.5 Hz, 1H), 6.23 (dd, *J* = 12.0, 4.9 Hz, 1H), 7.23 – 7.34 (m, 3H), 7.43 (dd, *J* = 8.6, 4.2 Hz, 1H), 7.61 (dd, *J* = 8.6, 7.3 Hz, 1H), 7.69 (d, *J* = 2.3 Hz, 1H), 8.01 (d, *J* = 8.6 Hz, 1H), 8.29 (d, *J* = 8.6 Hz, 1H), 8.90 (dd, *J* = 4.2, 1.6 Hz, 1H). $^{13}$**C NMR** (151 MHz, $CDCl_3$) $\delta_C$ 42.1, 44.9, 57.6, 121.5, 122.8, 125.0, 129.6, 129.8, 130.2, 131.2, 131.4, 131.5, 131.6, 132.4, 133.3, 136.0, 148.6, 150.1, 154.1, 164.7. **HRMS (ESI):** exact mass calculated for $C_{20}H_{14}Cl_3N_3O$ $[(M+H)^+]$ 418.0275, found 418.029.

**2-Chloro-1-(3-(5-chloro-2-fluorophenyl)-5-(quinolin-5-yl)-4,5-dihydro-1*H*-pyrazol-1-yl)ethan-1-one (CMZ 53)**

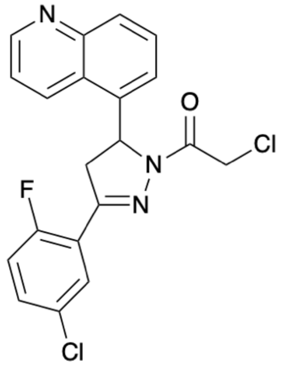


*Step 1*: Following General Procedure A, the chalcone was prepared from quinoline-5-carbaldehyde (207 mg, 1.31 mmol), 5′-chloro-2′-fluoroacetophenone (352 μL, 2.63 mmol) and 5% (w/w) aq NaOH (452 μL, 0.57 mmol) in EtOH (4 mL). Filtration under reduced pressure and purification *via* flash column chromatography (**Sfär Silica HC D**, 0% grading to 100% EtOAc in hexane, product eluted at 31%) gave the chalcone as a yellow solid (56 mg, 14%).
*Step 2*: Following General Procedure F, the pure title compound was prepared from chalcone (54 mg, 0.17 mmol) and hydrazine hydrate (31 μL, 0.35 mmol, 50–60% wt solution) in EtOH (4 mL); then chloroacetyl chloride (21 μL, 0.26 mmol) and triethylamine (72 μL, 0.52 mmol) in DCM (1.5 mL). Purification *via* flash column chromatography (**Sfär Silica HC D**, 0% grading to 100% EtOAc in hexane, product eluted at 52%) gave **CMZ 53** as a white solid (31 mg, 44% over 2 steps from the respective chalcone); $^1$**H NMR** (400 MHz,

$CDCl_3$) $\delta_H$ 3.09 – 3.31 (m, 1H), 3.88 – 4.04 (m, 1H), 4.52 (app dd, *J* = 13.4, 1.8 Hz, 1H), 4.63 (app dd, *J* = 13.4, 1.8 Hz, 1H), 6.22 (dd, *J* = 12.2, 5.0 Hz, 1H), 6.99 (ddd, *J* = 10.7, 8.7, 1.9 Hz, 1H), 7.24 (d, *J* = 7.3 Hz, 1H), 7.28 – 7.36 (m, 1H), 7.43 (ddd, *J* = 8.7, 4.1, 1.9 Hz, 1H), 7.52 – 7.66 (m, 1H), 7.86 – 7.95 (m, 1H), 8.00 (d, *J* = 8.7 Hz, 1H), 8.28 (d, *J* = 8.7 Hz, 1H), 8.90 (dt, *J* = 4.1, 1.9 Hz, 1H). **$^{13}$C NMR** (151 MHz, $CDCl_3$) $\delta_C$ 42.0, 44.4, 44.5, 57.3, 118.2, 118.4, 120.2, 120.3, 121.5, 122.9, 125.1, 128.5, 128.5, 129.6, 129.8, 130.2, 130.2, 131.9, 132.4, 132.5, 136.2, 148.3, 149.9, 151.5, 151.5, 159.0, 160.7, 164.7. **HRMS (ESI):** exact mass calculated for $C_{20}H_{14}Cl_2FN_3O$ [$(M+H)^+$] 402.0571, found 402.0560.

**2-Chloro-1-(3-(2,3-dichlorophenyl)-5-(quinolin-5-yl)-4,5-dihydro-1*H*-pyrazol-1-yl)ethan-1-one (CMZ 56)**

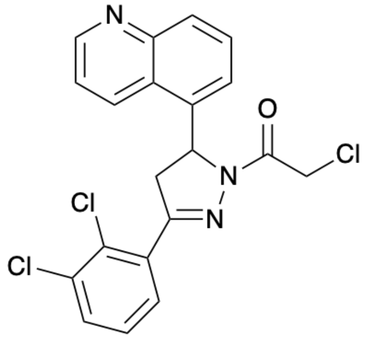


*Step 1*: Following General Procedure A, the chalcone was prepared from quinoline-5-carbaldehyde (213 mg, 1.36 mmol), 2′3′-dichloroacetophenone (396 μL, 2.71 mmol) and 5% (w/w) aq NaOH (466 μL, 0.58 mmol) in EtOH (4 mL). Filtration under reduced pressure and purification gave the chalcone as a pale brown solid (445 mg, quant).
*Step 2*: Following General Procedure F, the pure title compound was prepared from chalcone (306 mg, 0.93 mmol) and hydrazine hydrate (165 μL, 1.87 mmol, 50–60% wt solution) in EtOH (4 mL); then chloroacetyl chloride (111 μL, 1.40 mmol) and triethylamine (390 μL, 2.80 mmol) in DCM (4.7 mL). Purification *via* flash column chromatography (**Sfär Silica HC D**, 0% grading to 100% EtOAc in hexane, product eluted at 40%; then SNAP Ultra C18, 20% grading to 100% MeCN in water, product eluted at 63%) gave **CMZ 56** as a white solid (9 mg, 2.4% over 2 steps from the respective chalcone); **$^1$H NMR** (400 MHz, $CDCl_3$) $\delta_H$ 3.26 (dd, *J* = 17.7, 5.1 Hz, 1H), 4.09 (dd, *J* = 17.7, 11.9 Hz, 1H), 4.49 (d, *J* = 12.8 Hz, 1H), 4.65 (d, *J* = 12.8 Hz, 1H), 5.89 (dd, *J* = 11.9, 5.1 Hz, 1H), 7.03 (dd, *J* = 8.0, 1.6 Hz, 1H), 7.14 (t, *J* = 8.0 Hz, 1H), 7.36 (dd, *J* = 8.0, 1.6 Hz, 1H), 7.53 (dd, *J* = 8.7, 4.2 Hz, 1H), 7.58 (dd, *J* = 7.3, 1.5 Hz, 1H), 7.65 (dd, *J* = 8.7, 7.3 Hz, 1H), 8.14 (app dt, *J* = 8.4, 1.1 Hz, 1H), 8.95 (dd, *J* = 4.2, 1.5 Hz, 1H), 9.49 - 9.59 (m, 1H). **$^{13}$C NMR** (151 MHz, $CDCl_3$) $\delta_C$ 41.9, 43.5, 57.6, 122.6, 124.0, 126.2, 127.4, 128.0, 129.5, 129.6, 130.1, 130.2, 131.6, 134.1, 137.2, 129.5, 146.8, 149.2, 155.1, 164.3**. HRMS (ESI):** exact mass calculated for $C_{20}H_{14}Cl_3N_3O$ [$(M+H)^+$] 418.0275, found 418.0287.

**2-Chloro-1-(3-(2,5-difluorophenyl)-5-(quinolin-5-yl)-4,5-dihydro-1*H*-pyrazol-1-yl)ethan-1-one (CMZ 63)**

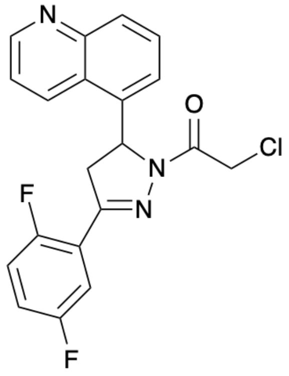


*Step 1*: Following General Procedure A, the chalcone was prepared from quinoline-5-carbaldehyde (200 mg, 1.27 mmol), 2′5′-difluoroacetophenone (161 μL, 1.27 mmol) and 5% (w/w) aq NaOH (438 μL, 0.55 mmol) in EtOH (4 mL). Filtration under reduced pressure and purification *via* flash column chromatography (**Sfär Silica HC D**, 0% grading to 100% EtOAc in hexane, product eluted at 44%) gave the chalcone as a yellow solid (59 mg, 16%).

*Step 2*: Following General Procedure F, the pure title compound was prepared from chalcone (59 mg, 0.20 mmol), hydrazine hydrate (35 μL, 0.40 mmol, 50–60% wt solution) in EtOH (4 mL); then chloroacetyl chloride (24 μL, 0.30 mmol) and triethylamine (84 μL, 0.60 mmol) in DCM (1.5 mL). Purification *via* flash column chromatography (**Sfär Silica HC D**, 0% grading to 100% EtOAc in hexane, product eluted at 56%) gave **CMZ 63** as a white solid (30 mg, 38% over 2 steps from the respective chalcone); **$^1$H NMR** (400 MHz, $CDCl_3$) $\delta_H$ 3.23 (ddd, *J* = 18.5, 5.1, 3.0 Hz, 1H), 4.00 (ddd, *J* = 18.5, 12.0, 3.0 Hz, 1H), 4.51 (d, *J* = 13.4 Hz, 1H), 4.62 (d, *J* = 13.4 Hz, 1H), 6.23 (dd, *J* = 12.0, 5.1 Hz, 1H), 6.94 – 7.12 (m, 2H), 7.26 (d, *J* = 7.2 Hz, 1H), 7.45 (dd, *J* = 8.6, 4.3 Hz, 1H), 7.58 – 7.69 (m, 2H), 8.03 (d, *J* = 8.6 Hz, 1H), 8.31 (d, *J* = 8.6 Hz, 1H), 8.91 (dd, *J* = 4.3, 1.6 Hz, 1H). **HRMS (ESI)**: exact mass calculated for $C_{20}H_{14}ClF_2N_3O$ [$(M+H)^+$] 386.0866, found 386.0881.
